# Bipolar multiplex families have an increased burden of common risk variants for psychiatric disorders

**DOI:** 10.1101/468975

**Authors:** Till F. M. Andlauer, Jose Guzman-Parra, Fabian Streit, Jana Strohmaier, Maria José González, Susana Gil Flores, Francisco J. Cabaleiro Fabeiro, Francisco del Río Noriega, Fermin Perez Perez, Jesus Haro González, Guillermo Orozco Diaz, Yolanda de Diego-Otero, Berta Moreno-Kuestner, Georg Auburger, Franziska Degenhardt, Stefanie Heilmann-Heimbach, Stefan Herms, Per Hoffmann, Josef Frank, Jerome C. Foo, Jens Treutlein, Stephanie H. Witt, Sven Cichon, Manolis Kogevinas, Bipolar Disorder Working Group of the Psychiatric Genomics Consortium, Major Depressive Disorder Working Group of the Psychiatric Genomics Consortium, Fabio Rivas, Fermín Mayoral, Bertram Müller-Myhsok, Andreas J. Forstner, Markus M. Nöthen, Marcella Rietschel

## Abstract

Multiplex families with a high prevalence of a psychiatric disorder are often examined to identify rare genetic variants with large effect sizes. In the present study, we analysed whether the risk for bipolar disorder (BD) in BD multiplex families is influenced by common genetic variants. Furthermore, we investigated whether this risk is conferred mainly by BD-specific risk variants or by variants also associated with the susceptibility to schizophrenia or major depression. In total, 395 individuals from 33 Andalusian BD multiplex families as well as 438 subjects from an independent, sporadic BD case-control cohort were analysed. Polygenic risk scores (PRS) for BD, schizophrenia, and major depression were calculated and compared between the cohorts. Both the familial BD cases and unaffected family members had significantly higher PRS for all three psychiatric disorders than the independent controls, suggesting a high baseline risk for several psychiatric disorders in the families. Moreover, familial BD cases showed significantly higher BD PRS than unaffected family members and sporadic BD cases. A plausible hypothesis is that, in multiplex families with a general increase in risk for psychiatric disease, BD development is attributable to a high burden of common variants that confer a specific risk for BD. The present analyses, therefore, demonstrated that common genetic risk variants for psychiatric disorders are likely to contribute to the high incidence of affective psychiatric disorders in the multiplex families. The PRS explained only part of the observed phenotypic variance and rare variants might have also contributed to disease development.

## Introduction

Bipolar disorder (BD), characterized by alternating episodes of mania and depression, has a lifetime prevalence of approximately 1% and is a substantial contributor to disability throughout the world^1^. Nevertheless, reliable data concerning the aetiology of BD remain scarce. The heritability of BD is estimated to be above 70%^2-4^, thus demonstrating an important genetic component in the development of the disorder. Genome-wide association studies (GWAS) in case/control samples have reported that common genetic risk factors with minor allele frequencies (MAF) of ≥1% explain a substantial proportion of the genetic risk for BD^5-11^: the heritability explained by such common variants is estimated to be 0.17-0.23 on a liability scale^12^. Common variants also make a substantial contribution to the development of schizophrenia (SCZ) and major depressive disorder (MDD)^13,14^. These three psychiatric disorders have a shared genetic component, whereby relatives of patients with BD have, in addition to BD, an increased risk for MDD and SCZ^15^. In fact, GWAS have shown that many genetic risk variants are associated with all three disorders^16-20^.

## Materials and Methods

### Sample description

Besides common variants with small individual effects, rare variants with large effect sizes may also contribute to BD development^21,22^. In theory, such highly penetrant variants should be enriched in families with a high prevalence of illness, termed multiplex families, in comparison to sporadic BD cases. However, it remains unclear whether and to what extent disease incidence in multiplex families is caused by rare variants, a high load of common variants, or a combination of both.

To elucidate the molecular genetic causes of BD, we established the Andalusian Bipolar Family (ABiF) study in 1997, which recruited BD multiplex families^23-25^. In the present analyses, we first investigated whether common genetic variants make a significant contribution to the occurrence of BD in ABiF families. Next, we examined whether BD development was attributable to (a) BD-specific risk variants, (b) variants conferring risk to all three disorders BD, MDD, and SCZ, or, (c), a combination of both. To this end, polygenic risk scores (PRS) based on GWAS of BD, MDD, and SCZ were calculated for and compared between ABiF family members and sporadic BD cases and unscreened controls from the same population.

The ABiF study recruited BD multiplex families in Andalusia, Spain^23-25^. The present analyses included 395 members of 33 ABiF families. Diagnoses were assigned by two trained clinicians according to DSM IV criteria using the best estimate approach^23^. Diagnoses comprised (Table 1): BD, n=166 (FAM_BD_); MDD, n=78 MDD (FAM_MDD_); no history of an affective disorder n=151 (FAM_unaffected_). Six unaffected individuals with a history of substance abuse were excluded from the analyses. Forty-four subjects have married into the families and had no parent in the ABiF cohort (36 unaffected; 8 MDD). Furthermore, an independent, previously reported Spanish BD case/control (CC) sample was analyzed^9^.

**Table 1:** Characteristics of 389 individuals from the 33 ABiF families.

|  | **BD**<br>(n=166) | **MDD**<br>(n=78) | **Unaffected**<br>(n=145) | **Significance level of differences between diagnostic groups** |
| --- | --- | --- | --- | --- |
| **Age at interview** | 40.5 (18.53) | 44.5 (17.05) | 44 (22.24) | $\chi^2=3.892$; $p=0.143$ |
| **Age at onset** | 20 (7.41)<br>*missing=3* | 26 (11.86)<br>*missing=1* | | $\chi^2=17.482$; $p<0.001$ |
| **Gender (% female)** | 103 (62.05) | 54 (69.23) | 51 (35.17) | $\chi^2=32.210$; $p<0.001$ |
| **Married-in** | 0 (100.0) | 8 (10.03) | 36 (24.83) | $\chi^2=47.665$; $p<0.001$ |
| **Educational level** | | | | $\chi^2=2.092$; $p=0.143$ |
| Primary school | 118 (71.51) | 53 (67.95) | 102 (70.34) |  |
| Secondary school | 39 (23.63) | 18 (23.08) | 36 (24.83) |  |
| University degree | 8 (4.85)<br>*missing=1* | 7 (8.97) | 7 (4.83) |  |
| **Severe impairment during the disorder** | 105 (65.62)<br>*missing=6* | 4 (5.55)<br>*missing=6* | | $\chi^2=71.931$; $p<0.001$ |
| **History of psychosis** | 110 (66.26) | 4 (5.13) | | $\chi^2=101.218$; $p<0.001$ |
| **History of suicide attempts** | 41 (24.70) | 2 (2.56) | 1 (0.69)<br>*missing=1* | $\chi^2=51.670$; $p<0.001$ |

Six unaffected individuals with a history of substance abuse were excluded from the analyses and are not shown in this table. *Missing*: number of individuals with missing data. Age was analysed using the Kruskal-Wallis test; median and median absolute deviation are shown. All other values were analysed using chi-squared ( $\chi^2$ ) tests with two degrees of freedom; number (n) and percentage (%) of subjects are shown. Note that of the 44 married-in individuals listed here, only 35 unaffected, married-in family members were part of the combined FAM+CC dataset. These subjects were excluded from analyses of the combined dataset unless specified otherwise.

After quality control (QC), the combined dataset of both cohorts comprised data from 384 FAM (163 FAM_BD_, 73 FAM _MDD_, 142 FAM_unaffected_, and 6 FAM_unaffected_ with a history of substance abuse) and 438 CC subjects (161 sporadic BD cases and 277 unscreened controls). This dataset contained 35 unaffected, married-in family members who were excluded from analyses using the combined sample (unless specified otherwise). A detailed description of QC procedures is provided in the Supplementary Methods.

The study was approved by the respective local ethics committees, and all participants provided written informed consent. For five adolescents (age 15-17 years), written informed consent was also obtained from parents.

### Genotyping and Imputation

Genome-wide genotyping of the FAM sample was carried out using the Illumina Infinium PsychArray BeadChip (PsychChip). QC and population substructure analyses were performed in PLINK^26^, as described in the Supplementary Methods. Genotyping and basic QC of the CC sample were conducted previously and are described elsewhere^9^. The genotypes of the CC dataset were merged with those of the FAM sample.

Genotype data were imputed to the 1000 Genomes phase 3 reference panel using SHAPEIT and IMPUTE2^27-29^. After imputation and post-imputation QC, the combined dataset of both cohorts contained 6862461 variants with an INFO metric of ≥0.8 and a MAF of ≥1%. The imputed FAM dataset without the CC subjects contained 8628089 variants.

### Calculation of polygenic risk scores

PRS were calculated in *R* v3.3^30^ using imputed genetic data. For each PRS, the effect sizes of variants below a selected p-value threshold, both obtained from large GWAS (training data), were multiplied by the imputed SNP dosage in the test data and then summed to produce a single PRS per threshold. The weighted PRS thus represent cumulative, additive risk. For each disorder, ten PRS based on different GWAS p-value thresholds were calculated, ranging from *p*_*PRS*_<5×10^−8^ to *p*_*PRS*_<0.2. The number of SNPs used for each PRS is shown in Supplementary Table S1. For additional details, see the Supplementary Methods.

For BD, MDD, and SCZ diagnoses, summary statistics of GWAS by the Psychiatric Genomics Consortium (PGC) were used as training data. For BD, the data freeze contained 20352 cases and 31358 controls, after exclusion of the Spanish samples^12^. For MDD and SCZ, published datasets were used. These contained 130664 cases and 330470 controls for MDD^14^ and 33640 cases and 43456 controls for SCZ^13^. Variants with an INFO metric of <0.6 in the GWAS summary statistics were removed. For comparison, PRS for late-onset Alzheimer’s disease (LOAD) were calculated, based on a GWAS by the International Genomics of Alzheimer’s Project (IGAP) with 17008 cases and 37154 controls^31^. For further details, see the Supplementary Methods.

*Shared* psychiatric PRS were generated using all variants showing an association at *p*<0.05 in the GWAS of BD, SCZ, and MDD and for which effect sizes pointed in the same direction across studies. For this shared set of variants, *p*-values were calculated using random-effects meta-analysis. Genome-wide inferred statistics (GWIS) were calculated as published elsewhere^32^. As recommended for this method, variants with an INFO metric of <0.9 or >1.1 were removed. Furthermore, 10000 random PRS for each of the ten p-value thresholds were calculated. To this end, random variants from across the genome were drawn, using the same number of variants as for the BD PRS at each threshold and random effect sizes from the pool of all available BD, SCZ, and MDD effects.

### Statistical Analysis

PRS analyses were conducted in *R* using the function *polygenic* of the package *GenABEL*^*33*^, which implements a linear mixed model that takes family structure into account. Test statistics, including 95% confidence intervals (CI), were calculated using bootstrapping (package *boot*^34,35^). Following the hypothesis that family members or subjects with a psychiatric diagnosis have increased PRS for psychiatric disorders, one-sided *p*-values were calculated for all PRS-based analyses. Since the residuals were not normally distributed, *p*-values were confirmed using permutation analysis (10000 permutations). Significance thresholds were corrected for multiple testing using the Bonferroni method. For further details see the Supplementary Methods.

In analyses of the combined FAM and CC dataset, sex was used as a covariate. In the analysis of FAM data alone, sex, the age at the interview, and whether an individual had married into the family were used as covariates. Age was not available for the CC controls.

## Results

### FAM_BD_ cases had higher psychiatric PRS than controls from the general population

We first examined whether FAM_BD_ cases, compared to independent CC_controls_, had higher PRS for the three psychiatric disorders BD, SCZ, and MDD. Selected results for these comparisons are shown in Fig. 1 *(i*.*e*., for each type of PRS, the test statistics for the PRS with the threshold *p*_*PRS*_ that showed the strongest association); full results for all ten thresholds per PRS type are provided in Supplementary Figs. S1-S2 and Supplementary Table S2. On average, FAM_BD_ cases had higher BD PRS than CC_controls_ at all ten thresholds. The most substantial support for an increased BD PRS was found with the threshold *p*_*PRS*_=0.1 (one-sided p=1.9×10^−20^, significance threshold α=0.05/10=5×10^−3^). FAM_BD_ cases also had significantly higher SCZ and MDD PRS than CC_controls_. While the BD and SCZ PRS were increased across all thresholds, the association using MDD PRS was only significant for *p*_*PRS*_*≥* 0.1.

**Figure 1:**
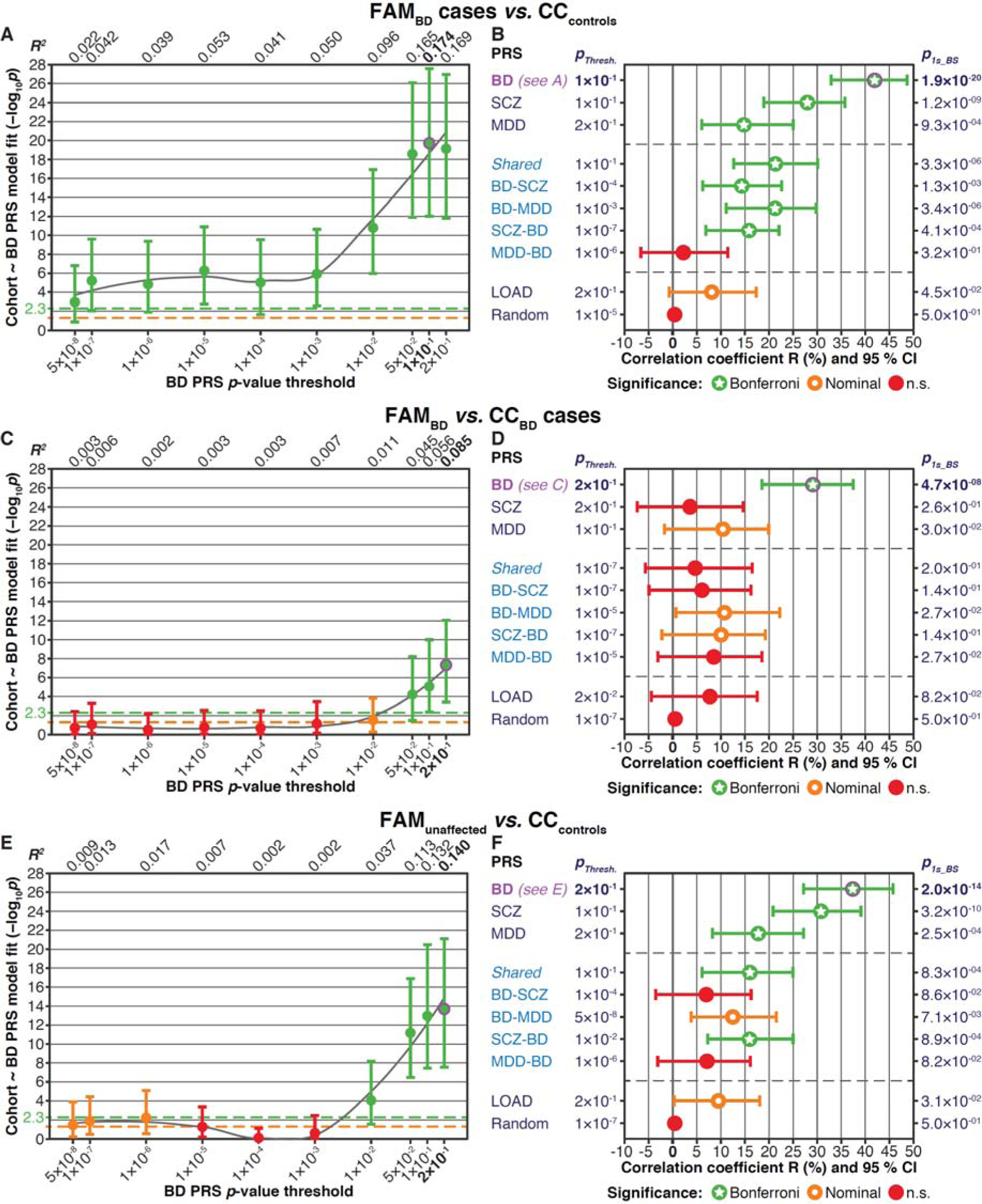
Association analyses comparing the various PRS between the FAM and CC samples. Married-in family members were excluded from these analyses. The plots show one-sided *p*-values, following the hypothesis that family members have higher PRS than individuals from the CC samples. **A-B:** Comparison of FAM_BD_ cases to CC_controls_. **A:** FAM_BD_ cases had higher BD PRS across all ten *p*_*PRS*_ thresholds. The plot shows one-sided *p*-values (filled circles), and 95% CI calculated using bootstrapping. The various *pPRS* thresholds are shown on the x-axis, and the association strength (-log_10_ *p*) is shown on the y-axis. The coefficient of determination *R*^*2*^ of the linear models is indicated at the top of the plot. Orange dashed line: nominal significance threshold (α=0.05), green dashed line: significance threshold after Bonferroni correction for multiple testing (α=0.05/10=0.005). Results foreach threshold are coloured in accordance with their degree of significance: red = not significant, orange = nominally significant, green = significant after correction for multiple testing. The top-associated PRS (*p*_*PRS*_=0.1) is indicated in bold font and was marked by a magenta circle (also in B). **B:** For ten different PRS, this plot shows association statistics for the top-associated *p*_*PRS*_ thresholds. The x-axis shows the association strength as the coefficient of multiple correlation *R* and 95% CI calculated by bootstrapping. Note that *R* instead of *R*^*2*^ has been used in this plot to allow for directionality. BD, SCZ, MDD: Standard PRS using the respective PGC GWAS summary statistics. *Shared:* Shared psychiatric PRS (SNPs with BD, MDD, SCZ *p*<0.05, random effects meta-analysis). BD-SCZ, BD-MDD: BD-specific GWIS PRS corrected for SCZ and MDD, respectively. SCZ-BD and MDD-BD: GWIS PRS for SCZ and MDD, each corrected for BD. LOAD: PRS for late-onset Alzheimer’s disease. Random: Mean and CI of the 10000 random PRS at the *p*_*PRS*_ with the lowest mean association *p*-value of all random PRS across *p*_*PRS*_. The column to the left of the plot: *p*_*PRS*_ with the strongest association. Supplementary Fig. S2 shows plots for all *p*_*PRS*_. Column to the right: *p*_*1s*_*_* _*BS*_ = one-sided *p*-value calculated using bootstrapping. For full association test statistics, see Supplementary Table S2. Abbreviations: Bonferroni = significant after Bonferroni correction for multiple testing; nominal = nominally significant (*p*<0.05); n.s. = not significant. **C-D:** Comparison of FAM_BD_ cases and sporadic CC_BD_ cases. See Supplementary Fig. S3 and Table S3 for more detailed plots and full association test statistics. **E-F:** Comparison of FAM_unaffected_ and CC_controls_. See Supplementary Fig. S4 and Table S4 for more detailed plots and full association test statistics.

BD and SCZ, and, to a lesser degree, BD and MDD, show a strong genetic correlation^14,16,18-20^. We, therefore, analysed whether variants associated with all three disorders contributed to the increased psychiatric PRS in FAMBD cases. *Shared* PRS generated from the variants associated with BD, SCZ, and MDD were increased at *p*_*PRS*_*≥* 0.01 in FAM_BD_ cases compared to CC_controls_. The additional contribution of disorder-specific variants was examined using GWIS^32^, which is a method previously applied to the analysis of genetic factors unique to either BD or SCZ. BD GWAS summary statistics were thus corrected for the MDD GWAS results to calculate a BD-MDD PRS less biased by the genetic correlation of BD with MDD. The BD-MDD PRS were significantly increased in FAM_BD_ cases compared to CC_controls_ across all thresholds. When correcting BD summary statistics for SCZ, BD-SCZ PRS were only associated at the single *p*_*PRS*_=1×10^−4^. Furthermore, the SCZ-BD GWIS PRS was significantly increased in FAM_BD_ cases at *p*_*PRS*_<1×10^−7^, whereas the MDD-BD PRS was not. These results indicate that FAM_BD_ cases had a higher load of common risk variants for BD, SCZ, and MDD than the population-based controls. This difference was mainly attributable to variants specific to BD. However, the increased risk of FAM_BD_ cases was also influenced by variants that confer risk for all three psychiatric diagnoses.

Finally, to confirm that the FAMBD cases had an increased PRS specifically for the tested psychiatric disorders but not for unrelated diseases, a PRS for late-onset Alzheimer’s disease (LOAD) was calculated. No significant increase was found at any threshold. In addition, 10000 random PRS were generated for each of the ten thresholds. The threshold at which the lowest mean association *p*-value across all 10000 random PRS was observed (*p*_*PRS*_=1 *10^−5^; *p*=0.496) was selected for comparisons. Associations of the PRS for BD, MDD, and SCZ, and of the *Shared* PRS were significantly stronger than this random PRS in FAMBD compared to CC_controls_ (Table 2). These results demonstrate that the observed increase in the risk burden of FAM_BD_ cases compared to CC_controls_ was not attributable to systematic, genome-wide genetic differences between the cohorts.

**Table 2:**
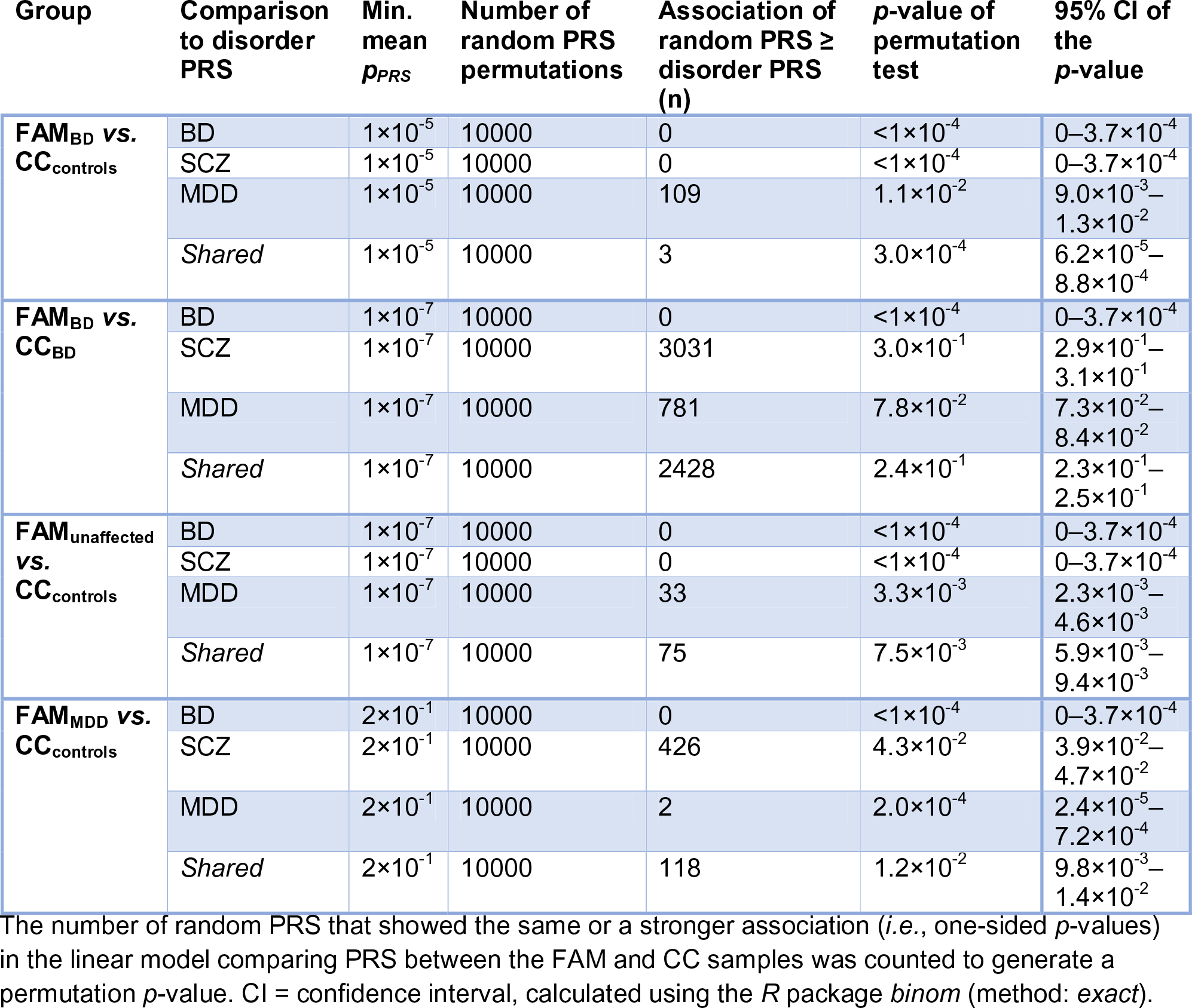
Permutation test of 10000 random PRS at the *p*-value threshold showing the lowest mean association *p*-value (min. mean *p*_*PRS*_).

| Group | Comparison to disorder PRS | Min. mean $p_{PRS}$ | Number of random PRS permutations | Association of random PRS ≥ disorder PRS (n) | $p$-value of permutation test | 95% CI of the $p$-value |
| --- | --- | --- | --- | --- | --- | --- |
| **FAM<sub>BD</sub> vs. CC<sub>controls</sub>** | BD | $1 \times 10^{-5}$ | 10000 | 0 | $<1 \times 10^{-4}$ | $0-3.7 \times 10^{-4}$ |
| | SCZ | $1 \times 10^{-5}$ | 10000 | 0 | $<1 \times 10^{-4}$ | $0-3.7 \times 10^{-4}$ |
| | MDD | $1 \times 10^{-5}$ | 10000 | 109 | $1.1 \times 10^{-2}$ | $9.0 \times 10^{-3}-1.3 \times 10^{-2}$ |
| | *Shared* | $1 \times 10^{-5}$ | 10000 | 3 | $3.0 \times 10^{-4}$ | $6.2 \times 10^{-5}-8.8 \times 10^{-4}$ |
| **FAM<sub>BD</sub> vs. CC<sub>BD</sub>** | BD | $1 \times 10^{-7}$ | 10000 | 0 | $<1 \times 10^{-4}$ | $0-3.7 \times 10^{-4}$ |
| | SCZ | $1 \times 10^{-7}$ | 10000 | 3031 | $3.0 \times 10^{-1}$ | $2.9 \times 10^{-1}-3.1 \times 10^{-1}$ |
| | MDD | $1 \times 10^{-7}$ | 10000 | 781 | $7.8 \times 10^{-2}$ | $7.3 \times 10^{-2}-8.4 \times 10^{-2}$ |
| | *Shared* | $1 \times 10^{-7}$ | 10000 | 2428 | $2.4 \times 10^{-1}$ | $2.3 \times 10^{-1}-2.5 \times 10^{-1}$ |
| **FAM<sub>unaffected</sub> vs. CC<sub>controls</sub>** | BD | $1 \times 10^{-7}$ | 10000 | 0 | $<1 \times 10^{-4}$ | $0-3.7 \times 10^{-4}$ |
| | SCZ | $1 \times 10^{-7}$ | 10000 | 0 | $<1 \times 10^{-4}$ | $0-3.7 \times 10^{-4}$ |
| | MDD | $1 \times 10^{-7}$ | 10000 | 33 | $3.3 \times 10^{-3}$ | $2.3 \times 10^{-3}-4.6 \times 10^{-3}$ |
| | *Shared* | $1 \times 10^{-7}$ | 10000 | 75 | $7.5 \times 10^{-3}$ | $5.9 \times 10^{-3}-9.4 \times 10^{-3}$ |
| **FAM<sub>MDD</sub> vs. CC<sub>controls</sub>** | BD | $2 \times 10^{-1}$ | 10000 | 0 | $<1 \times 10^{-4}$ | $0-3.7 \times 10^{-4}$ |
| | SCZ | $2 \times 10^{-1}$ | 10000 | 426 | $4.3 \times 10^{-2}$ | $3.9 \times 10^{-2}-4.7 \times 10^{-2}$ |
| | MDD | $2 \times 10^{-1}$ | 10000 | 2 | $2.0 \times 10^{-4}$ | $2.4 \times 10^{-5}-7.2 \times 10^{-4}$ |
| | *Shared* | $2 \times 10^{-1}$ | 10000 | 118 | $1.2 \times 10^{-2}$ | $9.8 \times 10^{-3}-1.4 \times 10^{-2}$ |

The number of random PRS that showed the same or a stronger association (*i.e.*, one-sided $p$ -values) in the linear model comparing PRS between the FAM and CC samples was counted to generate a permutation $p$ -value. CI = confidence interval, calculated using the *R* package *binom* (method: *exact*).

### FAM_BD_ cases had higher BD PRS than sporadic BD cases

We next compared the FAM_BD_ cases to sporadic BD cases from Andalusia. The BD PRS was significantly higher in FAM_BD_ than in CC_BD_ cases at *p*_*PRS*_*≥* 0.05 (Fig. 1C-D, Supplementary Figs. S1 and S3, Supplementary Table S3). However, no other type of PRS was increased in FAM_BD_ compared to CC_BD_ cases. The association of BD PRS was significantly stronger than the association of any random PRS in this analysis (Table 2), which confirmed that the FAM_BD_ cases carried an exceptionally high load of common risk variants for BD.

### Unaffected family members showed higher psychiatric PRS than CC controls

In the comparison of FAM_unaffected_ to CC_controls_, PRS for BD, SCZ, MDD, and the *Shared* PRS were significantly higher in unaffected family members (Fig. 1E-F, Supplementary Figs. S1 and S4, Supplementary Table S4). FAM_unaffected_ individuals showed no increase in BD/MDD-specific GWIS, LOAD, or random PRS. However, their SCZ-BD GWIS PRS were higher. The associations of the psychiatric PRS were significantly stronger than the association of random PRS (Table 2). FAM_unaffected_ individuals, therefore, had a substantial risk load for all three psychiatric disorders and did not show a specific increase in risk for BD.

### FAM_BD_ cases had an increased PRS specifically for BD

The preceding analyses have established that all family members showed an increased burden of common risk variants for BD, MDD, and SCZ and that FAM_BD_ cases appeared to carry a particularly high load of BD-associated variants.

To investigate FAM_BD_ cases further, we compared them directly to FAM_unaffected_. Here, only the BD PRS was significantly higher in FAM_BD_ (Fig. 2A-B, Supplementary Figs. S1 and S5, Supplementary Table S5).

**Figure 2:**
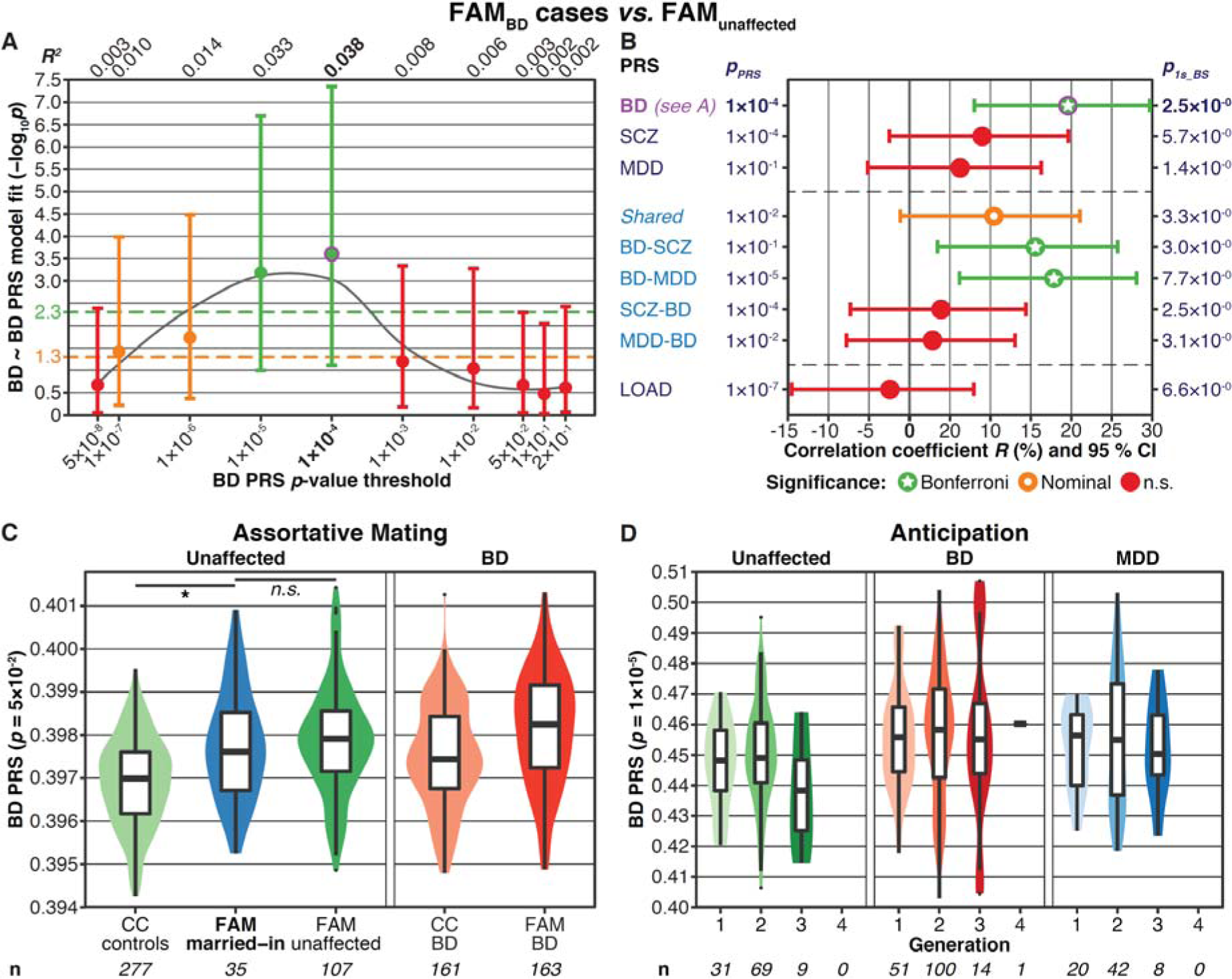
**A-B:** Association analyses comparing different PRS in FAM_BD_ cases to FAM_unaffected_. The plots show one-sided *p*-values, following the hypothesis that BD cases have higher PRS than unaffected individuals. Further details of the plots are as described in the legend for Fig. 1. See Supplementary Fig. S5 and Table S5 for more detailed plots and full association test statistics. **C-D:** Analyses of assortative mating (**C**) and anticipation (**D**). These plots were not adjusted for covariates; n = sample size. The y-axis shows the PRS values. **C:** Assortative mating. The plot shows violin- and boxplots of the BD PRS (*p*_PRS_=0.05), comparing unaffected, married-in individuals with no parent among the ABiF families to other FAM and CC subjects. At *p*_*PRS*_=0.05, married-in family members showed the highest BD PRS compared to CC_controls_ (*p*=1.1×10^−5^, Supplementary Fig. S6A and Table S6). The BD PRS of married-in individuals was not significantly higher than the PRS of FAM_unaffected_ at any *p_PRS_* (p≥0.186, Supplementary Fig. S6B and Table S6). Covariate used: gender. One-sided *p*-values were calculated, following the hypothesis that married-in individuals have higher PRS than other unaffected subjects. **D:** Anticipation: the BD PRS did not increase across generations. The plot shows violin- and boxplots of the BD PRS (*p*_*PRS*_=1 ×10^−5^) across different generations of the FAM sample for the three diagnosis groups. At *p*_*PRS*_*=1* ×10^−5^, the association of the BD PRS with generation was strongest (*p*=0.45, Supplementary Fig. S7A and Table S7). Married-in family members were excluded from this analysis. Covariates used: gender, age at the interview, diagnostic group. One-sided *p*-values were calculated, following the hypothesis that the PRS increase across generations.

Notably, while the BD-specific GWIS PRS were significantly higher, the *Shared* PRS were not. These results corroborate the observation that, while all family members had an increased load of risk variants for BD, SCZ, and MDD, the FAM_BD_ cases showed an especially high burden of variants that conferred a specific risk for BD.

### Effects of assortative mating on BD PRS in family members

The high burden of psychiatric risk in the ABiF families may be attributable to assortative mating. To investigate this hypothesis, we analysed the married-in family members without a parent within the FAM sample. Eight of the 44 individuals who had married into the families had a diagnosis of MDD and none of BD (Table 1). While the unaffected married-in individuals had higher BD PRS than CC_controls_, their BD PRS was not higher than the PRS of other FAM_unaffected_ (Fig 2C, Supplementary Fig. S6, Supplementary Table S6). We also examined possible anticipation of BD in the families: neither did the BD PRS increase significantly over generations nor did the age at onset decrease over time (Fig. 2D, Supplementary Fig. S7, Supplementary Table S7).

### FAM_MDD_ cases had higher psychiatric PRS than CC_controls_

Comparisons of the FAM_MDD_ cases to CC_controls_ demonstrated that FAM_MDD_ had significantly higher BD, MDD, and *Shared* PRS (Supplementary Figs. S8-S9, Supplementary Table S8). However, none of these PRS were significantly increased when comparing FAM_MDD_ to FAM_unaffected_ (Supplementary Fig. S10, Supplementary Table S9). Notably, in both comparisons, FAM_MDD_ showed an increase in SCZ-MDD GWIS PRS but not in SCZ-BD GWIS PRS.

## Discussion

The main aim of the present study was to investigate whether common genetic variants contribute to the occurrence of BD in 33 BD multiplex families from Andalusia, Spain. For this purpose, PRS were calculated for the family members and compared to PRS of independent controls from the Spanish population and sporadic BD cases. In comparison to CC_controls_, FAM_BD_ cases had significantly increased PRS for the three psychiatric disorders BD, SCZ, and MDD, higher BD+SCZ+MDD *Shared* PRS, and higher BD-specific PRS. Unaffected family members had, compared to CC_controls_, significantly increased PRS for BD, SCZ, and MDD, and higher *Shared* PRS. When comparing FAM_BD_ to sporadic CC_BD_ cases, only the BD PRS were significantly higher in the FAM sample. Also within the ABiF families, only the BD PRS were higher in FAMBD cases than in FAM_unaffected_. These findings indicate that, compared to non-familial samples from Spain, the family members had a strongly elevated baseline risk for the genetically correlated psychiatric disorders BD, MDD, and SCZ. While this genetic burden confers a higher risk for mental illness in general, FAM_BD_ cases were characterized by a particularly high number of BD-specific risk variants.

Although both the FAM and CC samples were recruited in Spain^9^, minor population differences may have influenced the present results. Even if such minor differences existed, it is unlikely that they caused the highly significant associations observed for the psychiatric PRS, given that the pairwise genetic relationship matrix was used as random effects in the association analyses. Nevertheless, three analyses were conducted to confirm that systematic differences between the genotype data of FAM, CC_controls_, and CC_BD_ samples did not distort our findings: First, we did not find significant differences between the cohorts in a population substructure analysis (see Supplementary Fig. S11 and Supplementary Methods). Second, PRS for LOAD were not significantly increased in family members in any analysis. Since LOAD shows no genetic correlation with BD, MDD, or SCZ^143637^, this adds further support to the specificity of our analyses. Third, we generated random PRS and conducted permutation tests to determine how often random PRS showed the same or a more extreme association strength than PRS for psychiatric disorders. These analyses confirmed that when a psychiatric disorder PRS was significantly increased in family members, this association was stronger than for random PRS. We thus conclude that the high psychiatric PRS observed in family members compared to controls cannot be attributed to population or technical differences between the cohorts.

While the ABiF families are characterized by a high prevalence of BD and MDD, the family members showed increased SCZ PRS compared to controls. This increase could be an indirect consequence of the genetic correlation between BD and SCZ^14,16,18-20^. Interestingly, the majority of FAM_BD_ cases were diagnosed with BD-I (115 of 166 cases), which is the BD subtype with the highest genetic correlation to SCZ^12,38^. Furthermore, family members also had higher *Shared* PRS than CC_controls_. SCZ shows a higher heritability and lower prevalence than MDD does, and the number of cases in the GWAS used for calculating the SCZ PRS was higher than the number of cases in the BD GWAS. The SCZ GWAS is therefore likely to have had higher statistical power than the GWAS of BD and MDD^39^. In consequence, the SCZ PRS may have included more cross-disorder signals with smaller effects than the PRS of BD and MDD. If family members had an increased *Shared* risk burden, this cross-disorder risk may have rendered them vulnerable to psychiatric disorders in general, with the high BD PRS then shaping the final BD diagnosis outcome. Notably, none of the family members included in the present analyses has been diagnosed with SCZ. Given that the recruitment strategy focused on BD multiplex families, this is likely to be attributable to ascertainment bias. Of note, the analyses of FAM_MDD_ cases are discussed in the Supplementary Data.

The BD PRS of FAM_BD_ cases were significantly higher than those of CC_controls_ across all *p*_*PRS*_ thresholds (Fig. 1A). Similarly, the BD PRS of CC_BD_ cases were higher than in CC_controls_ across all thresholds (Supplementary Fig. S12, Supplementary Table S10). By comparison, the increase of the BD PRS in FAM_BD_ cases relative to FAM_unaffected_ was less pronounced, albeit significant at two *p*_*PRS*_ thresholds (Fig. 2A). Depending on the genetic architecture of the disorder in question and the statistical power of GWAS and PRS analyses, PRS may be expected to differ less in a family-based cohort. In a previous study of BD multiplex families^40^, in which the PRS was calculated using a different GWAS^10^, the BD PRS was not significantly higher in cases compared to unaffected family members. In contrast, another study found that PRS for LOAD significantly distinguished between Alzheimer’s disease cases and controls in familial cohorts^41^.

Our analyses showed that ABiF family members carried high BD PRS compared to the average population. Although individuals who married into these families also had higher BD PRS than CC_controls_, their BD risk load was similar to other FAM_unaffected_. At the time of the interview, none of the married-in family members had a diagnosis of BD. Nevertheless, their increased BD PRS indicate that weak assortative mating may have occurred. Unaffected individuals with an above average BD PRS may display sub-threshold characteristics of BD, such as a broader range of emotions^42-44^. Consistent with the observation that married-in subjects did not have higher BD PRS than the other FAM_unaffected_, no increase in BD PRS was found across generations. However, assortative mating may have contributed to the establishment and maintenance of a high genetic risk load for BD in these families. Furthermore, assortative mating may have already occurred in previous generations, for which no DNA was available. Of note, DNA was neither available for all ABiF family members of the current generations, limiting the scope of the analysis of assortative mating.

The present study generated substantial evidence that members of the ABiF families, including unaffected subjects, carried a higher risk burden of common genetic risk variants for the psychiatric disorders BD, SCZ, and MDD than the average population. A plausible hypothesis is that this polygenic load of common risk variants is a major contributor to the high incidence of BD and MDD in these families. Although a previous investigation of a single ABiF pedigree already described a high BD PRS in cases, no rare causal variants were identified^25^. Research suggests that for high prevalence complex diseases, even cases accumulated within families may typically be influenced by a polygenic risk burden^45^. However, given that the PRS explained only a fraction of the phenotypic variance (up to an *R*^*2*^ of 17.4% in the present analyses), rare mutations likely also played an important role in each of the families. Sequencing studies carried out in multiplex families have suggested a function of rare variants in the aetiology of BD46-48. To date, however, it has proven difficult to identify replicable causal associations between rare variants and BD susceptibility. Of note, we analysed genotype data from the 33 ABiF families as an average across families and not separately per family. Thus, the degree to which common and rare variants shaped the emergence of psychiatric disorders may vary between families. To further enhance our understanding regarding the aetiology of BD, future studies should aim at identifying such rare variants in single ABiF families and determine the contribution of rare variants to disease development in comparison to the impact of common variants.

Supplementary information is available at bioRxiv.org.

## Acknowledgments

The study was supported by the German Federal Ministry of Education and Research (BMBF), through the Integrated Network IntegraMent, under the auspices of the e:Med programme (grants 01ZX1314A to MMN and SC; 01ZX1314G to MR; 01ZX1614J to BMM), by the German Research Foundation (DFG grants FOR2107; RI908/11-1 to MR; NO246/10-1 to MMN; MU1315/8-2 to BMM), and by the Swiss National Science Foundation (SNSF grant 156791 to SC). MMN is a member of the DFG-funded cluster of excellence ImmunoSensation. The PGC has received major funding from the US National Institute of Mental Health and the US National Institute of Drug Abuse (U01 MH109528 and U01 MH1095320). We thank the research participants and employees of 23andMe, Inc. for their contribution to the MDD meta-analysis published in (14). We thank the International Genomics of Alzheimer’s Project (IGAP) for providing summary results data for the present analyses. See the Supplementary Data for extended Acknowledgments.

## Conflicts of Interest

The authors have no conflicts of interest to declare.

## Supplementary Information

### Supplementary Material

PDF document containing the Supplementary Methods, Discussion, References, Acknowledgments, and Figures S1-S12, legends for the Supplementary Tables and the full list of PGC authors. A detailed table of contents is provided at the beginning of the document.

### Supplementary Tables

Excel worksheet containing the Supplementary Tables S1-S10. Detailed legends for the Supplementary Tables are provided in the Supplementary Material PDF document. The Supplementary Tables can be accessed under goo.gl/3u1C66.

## Notes

#### Summary of Updates

authors and affiliations updated

