## Supplementary Material for "Bipolar multiplex families have an increased burden of common risk variants for psychiatric disorders"

### *Table of Contents*

**Supplementary Methods:** Sample description, quality control, population substructure analysis, imputation, generation and analysis of PRS.

**Supplement. Discussion:** Discussion of the analyses of $FAM_{MDD}$ cases.

**Supplement. References**

**Supplementary Fig. S1:** Boxplots of PRS at different *p*-value thresholds for $CC_{controls}$, $FAM_{unaffected}$, and BD cases.

**Supplementary Fig. S2:** Association analysis comparing PRS in $FAM_{BD}$ cases and $CC_{controls}$.

**Supplementary Fig. S3:** Association analysis comparing PRS in $FAM_{BD}$ cases and sporadic $CC_{BD}$ cases.

**Supplementary Fig. S4:** Association analysis comparing PRS in $FAM_{unaffected}$ and $CC_{controls}$.

**Supplementary Fig. S5:** Association analysis comparing PRS in $FAM_{BD}$ cases and $FAM_{unaffected}$.

**Supplementary Fig. S6:** Analysis of assortative mating.

**Supplementary Fig. S7:** Analysis of anticipation in the FAM sample.

**Supplementary Fig. S8:** Boxplots of PRS at different *p*-value thresholds, including $FAM_{MDD}$ cases.

**Supplementary Fig. S9:** Association analysis comparing PRS in $FAM_{MDD}$ cases and $CC_{controls}$.

**Supplementary Fig. S10:** Association analysis comparing PRS in $FAM_{MDD}$ cases and $FAM_{unaffected}$.

**Supplementary Fig. S11:** Population substructure analysis.

**Supplementary Fig. S12:** Association analysis comparing BD PRS in sporadic $CC_{BD}$ cases and $CC_{controls}$.

**IGAP Supplementary Methods and Acknowledgments**

**Authors of the Bipolar Disorder Working Group of the PGC**

**Authors of the Major Depressive Disorder Working Group of the PGC**

*Supplementary Tables in the separate Excel file:*

**Supplementary Table S1:** The number of variants used for calculation of each PRS (after clumping).

**Supplementary Table S2:** Full association test results for the comparison of PRS between $FAM_{BD}$ cases and $CC_{controls}$. Covariate used: Sex. Married-in family members were excluded.

**Supplementary Table S3:** Full association test results for the comparison of PRS between $FAM_{BD}$ cases and sporadic $CC_{BD}$ cases. Covariate used: Sex. Married-in family members were excluded.

**Supplementary Table S4:** Full association test results for the comparison of PRS between $FAM_{unaffected}$ and $CC_{controls}$. Covariates used: Sex. Married-in family members were excluded.

**Supplementary Table S5:** Full association test results for the comparison of PRS between $FAM_{BD}$ cases and $FAM_{unaffected}$. Covariates used: Sex, age at interview, married-in status.

**Supplementary Table S6:** Full association test results for the analysis of assortative mating. Covariate used: Sex.

**Supplementary Table S7:** Full association test results for the analysis of anticipation. Covariates used: Sex, age at the interview, diagnostic group (BD/MDD/unaffected). Married-in family members were excluded.

**Supplementary Table S8:** Full association test results for the comparison of PRS between $FAM_{MDD}$ cases and $CC_{controls}$. Covariate used: Sex. Married-in family members were excluded.

**Supplementary Table S9:** Full association test results for the comparison of PRS between $FAM_{MDD}$ cases and $FAM_{unaffected}$. Covariates used: Sex, age at the interview, married-in status.

**Supplementary Table S10:** Full association test results for the comparison of BD PRS between sporadic $CC_{BD}$ cases and $CC_{controls}$. Covariate used: Sex.

### Supplementary Methods

#### Extended FAM sample description

We included 395 members of 33 families in the present analyses. 166 participants were diagnosed with BD (BD type I (BD-I), n=115; BD type II (BD-II), n=41; not otherwise specified (NOS) BD, n=10), 78 with MDD (recurrent MDD (R-MDD), n=53; single episode MDD (SE-MDD), n=17; NOS MDD, n=8), and 151 without a history of an affective disorder.

Diagnoses were assigned by two trained clinicians according to DSM IV using the best estimate approach. Diagnosis and clinical data were based on the Schedule for Affective Disorders and Schizophrenia (SADS)[1], the Operational Criteria Checklist for Psychotic and Affective Illness (OPCRIT)[2], the Family Informant Schedule and Criteria (FISC)[3], and on clinical records.

A severe impairment during the disorder (see Table 1) corresponded to a level of 3 in the OPCRIT item 87 (no function at all in a major life role for more than two days or in-patient treatment has been required or active psychotic symptoms such as delusions or hallucinations have occurred).

#### Quality control (QC)

QC of genotype data was conducted in PLINK v1.90b3.36. QC was carried out first on each of both cohorts separately (FAM and CC), followed by a second round of QC on the combined dataset.

**Sequence of QC steps:**

1. FAM (genotyped on Infinium PsychArray BeadChip (PsychChip))
   Before QC: 395 individuals and 588,454 variants
   1.1. Removal of SNPs with call rates <98% or a MAF <1%
   1.2. Check for individuals with genotyping rates <98% (*none removed*)
   1.3. Check for sex mismatches (*none removed*)
   1.4. Removal of non-autosomal variants
   1.5. Removal of SNPs with call rates <98%, a MAF <1%, or Hardy-Weinberg Equilibrium (HWE) test *p*-values $<1\times10^{-6}$
   1.6. Removal of A/T and G/C SNPs
   1.7. Update of variant IDs and positions to the IDs and positions in the 1000 Genomes Phase 3 reference panel
   1.8. Alignment of alleles to the reference panel
   1.9. Removal of duplicated variants and of variants not present in the reference panel
   After QC: 395 individuals and 258,046 variants

2. CC (Illumina HumanOmni1-Quad and Illumina Human610-Quad, combined and quality-controlled as previously published[4]; the QC described here was conducted on the published data)
   Before QC: 547 individuals and 333,353 variants
   2.1. Removal of SNPs with call rates <98% or a MAF <1%
   2.2. Check for individuals with genotyping rates <98% (*none removed*)
   2.3. Check for sex mismatches (*none removed*)
   2.4. Check for genetic duplicates (*none removed*)
   2.5. Removal of individuals where the autosomal or X-chromosomal heterozygosity deviated from the mean >4 SD (*six removed*)
   2.6. Removal of non-autosomal variants
   2.7. Removal of SNPs with call rates <98%, a MAF <1%, or HWE test *p*-values $<1\times10^{-6}$
   2.8. Removal of A/T and G/C SNPs
   2.9. Update of variant IDs and positions to the IDs and positions in the 1000 Genomes Phase 3 reference panel
   2.10. Alignment of alleles to the reference panel
   2.11. Removal of duplicated variants and variants not present in the reference panel
   After QC: 541 individuals and 315,634 variants

3. Combined dataset of both samples (936 individuals and 116,079 variants)
   3.1. Removal of SNPs with call rates <98% or a MAF <1%
   3.2. Removal of individuals with genotyping rates <98% (*two removed*)
   3.3. Removal of individuals duplicated between both datasets (*31 removed from the CC sample*)
   3.4. Removal of genetic outliers with a distance from the mean of >4 SD in the first eight MDS components (*15 removed from the CC sample*)
   3.5. Removal of individuals where the autosomal heterozygosity deviated from the mean >4 SD (*eleven removed from the FAM sample*)
   3.6. Removal of SNPs with call rates <98%, a MAF <1%, or HWE test *p*-values $<1\times10^{-6}$
   3.7. Removal of individuals from the CC sample that have been recruited as part of the FAM/ABiF cohort (*55 removed*)
   After QC: 822 individuals (384 FAM and 438 CC) and 116,067 variants

**Population substructure analysis**

For the population substructure analyses, pre-imputation genotype data was used, after the QC steps explained above had been applied. Additional variant filtering steps were: removal of variants with a MAF <0.05 or HWE *p*-value $<10^{-3}$; removal of variants mapping to the extended MHC region (chromosome 6, 25-35 Mbp) or to a typical inversion site on chromosome 8 (7-13 Mbp); LD pruning (command *--indep-pairwise 200 100 0.2*).

Next, the pairwise identity-by-state (IBS) matrix of all individuals was calculated using the command *--genome* on the filtered genotype data. Multidimensional scaling (MDS) analysis was performed on the IBS matrix using the eigendecomposition-based algorithm in PLINK v1.90b5.

In an MDS analysis, the high relatedness between family members leads to artifacts. To avoid such artifacts, only one person per family was included in population substructure analyses. For each of the 33 families, the individual with the highest absolute values in MDS components 1 and 2 was selected to represent the respective family. Afterwards, the MDS analysis was repeated, using only selected individuals from the FAM sample.

Whether MDS components differ between cohorts was analyzed with logistic regression using the following model without additional covariates:
*cohort (FAM/CC) ~ MDS components.*
The ten calculated MDS components showed association *p*-values with cohort ≥0.30, except for component 3, which was associated with cohort at nominal significance with $p=0.036$. After correction for multiple testing (ten comparisons), this difference observed for component 3 was not significant.

**Imputation of genotype data**

Genotypes were aligned to the 1000 Genomes Phase 3 reference panel using SHAPEIT v2 (r837) and PLINK v1.90b3.36. Pre-phasing (haplotype estimation) was conducted for each chromosome separately using SHAPEIT. Imputation was performed using IMPUTE2 v2.3.2 in 5 Mbp chunks with 500 kbp buffers, filtering out variants that are monomorphic in the EUR samples. Chunks with <51 genotyped variants or concordance rates <92 % were fused with neighboring chunks and re-imputed. Imputed variants with a MAF <1% or an INFO metric <0.8 were removed.

| | |
|---|---|
| Imputed variants in the combined sample after QC: | 6,862,461 |
| Imputed variants in the FAM sample after QC: | 8,628,089 |

Note that optimized imputation algorithms for pedigrees exist, for example, GIGI[5]. GIGI mainly improves imputation accuracy of rare variants but does not have a clear advantage over population-based methods regarding common variants. Moreover, GIGI can only impute one pedigree at a time and cannot impute unrelated individuals. As we were only interested in common variation (MAF ≥1%) and also wanted to analyse a mixed sample of related and unrelated subjects, we chose a population-based imputation method using SHAPEIT and IMPUTE2.

#### Generation and analysis of PRS

The GWAS test statistics and imputed variants in our data were merged based on chromosome, position, and alleles of each variant. Summary statistics were then clumped in PLINK v1.90b5.2, based on best-guess genotype data (hard-call threshold 0.3) using the following parameters:

--clump-kb 500 --clump-r2 0.1 --clump-p1 1 --clump-p2 1

PRS were then calculated in *R* v.3.3 based on imputed (dosage) data. Test statistics and alleles in the GWAS training data were flipped so that effect sizes were always positive. Thus, the weighted PRS represent cumulative, additive risk. PRS were scaled to represent the relative risk load (minimum possible cumulative risk load = 0, maximum = 1). For each disorder, ten PRS with different *p*-value thresholds were calculated: $<5\times10^{-8}$, $<1\times10^{-7}$, $<1\times10^{-6}$, $<1\times10^{-5}$, $<1\times10^{-4}$, <0.001, <0.01, <0.05, <0.1, <0.2.

In the first step of the PRS analyses, residuals were calculated with the GenABEL *polygenic* function using the formula *phenotype ~ covariates* (where the phenotype corresponded to the diagnosis/cohort groups contrasted in a given analysis), including the genetic relationship matrix as random effects. Residuals from this model were then used in a second linear model with the formula *residuals ~ PRS*. Test statistics including 95% CI were calculated using bootstrapping (*R* package boot, nonparametric bootstrapping using ordinary resampling with 2,000 replications).

### Supplementary Discussion

$FAM_{MDD}$ cases had higher BD, MDD, and *Shared* PRS than $CC_{controls}$. The SCZ-MDD GWIS PRS were also increased in $FAM_{MDD}$ but the SCZ and SCZ-BD GWIS PRS were not higher. This may be due to the lower genetic correlation of MDD and SCZ compared to the correlation of BD and SCZ[6-9]. However, when interpreting the results for $FAM_{MDD}$ cases, it is important to consider that much fewer MDD than BD cases have been analysed (Table 1). The power of MDD-based analyses in the present study was thus considerably lower than for BD. This lower statistical power is also a possible explanation for why $FAM_{MDD}$ cases only showed a nominally increased MDD PRS over $FAM_{unaffected}$ individuals. In addition to suggesting a cross-disorder illness burden, the increased BD PRS in $FAM_{MDD}$ cases may also indicate that, in some cases, the current MDD diagnosis constituted a prodromal stage of BD[10]. Furthermore, in ABiF families, MDD may be more strongly driven by BD risk variants and therefore have closer etiological proximity to BD than is the case for the average MDD patient from the general population.

**Supplementary Fig. S1:** Boxplots of PRS at different *p*-value thresholds for $CC_{controls}$, $FAM_{unaffected}$, and BD cases. FAM samples excluded from the analyses of the combined dataset are not shown in these plots, *i.e.*, family members with a history of substance abuse, married-in family members, or family members diagnosed with MDD. CC = Case/control sample.

**Supplementary Fig. S1A:** Boxplots of BD PRS.

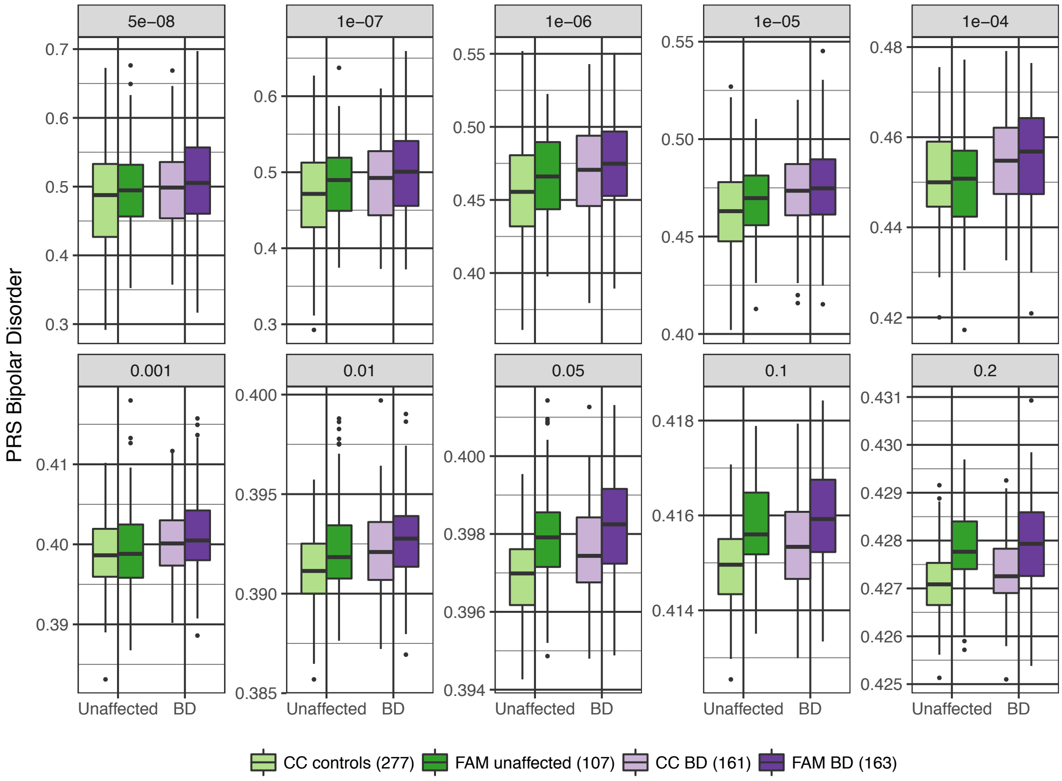


**Supplementary Fig. S1B:** Boxplots of SCZ PRS.

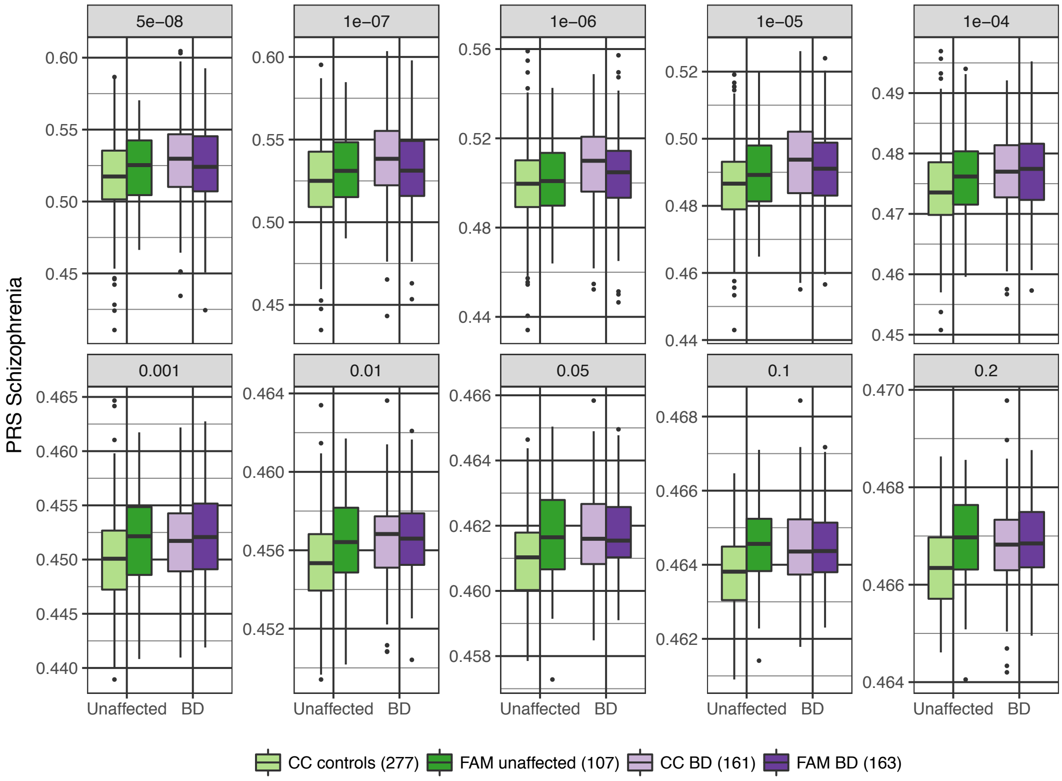

**Supplementary Fig. S1C:** Boxplots of MDD PRS.

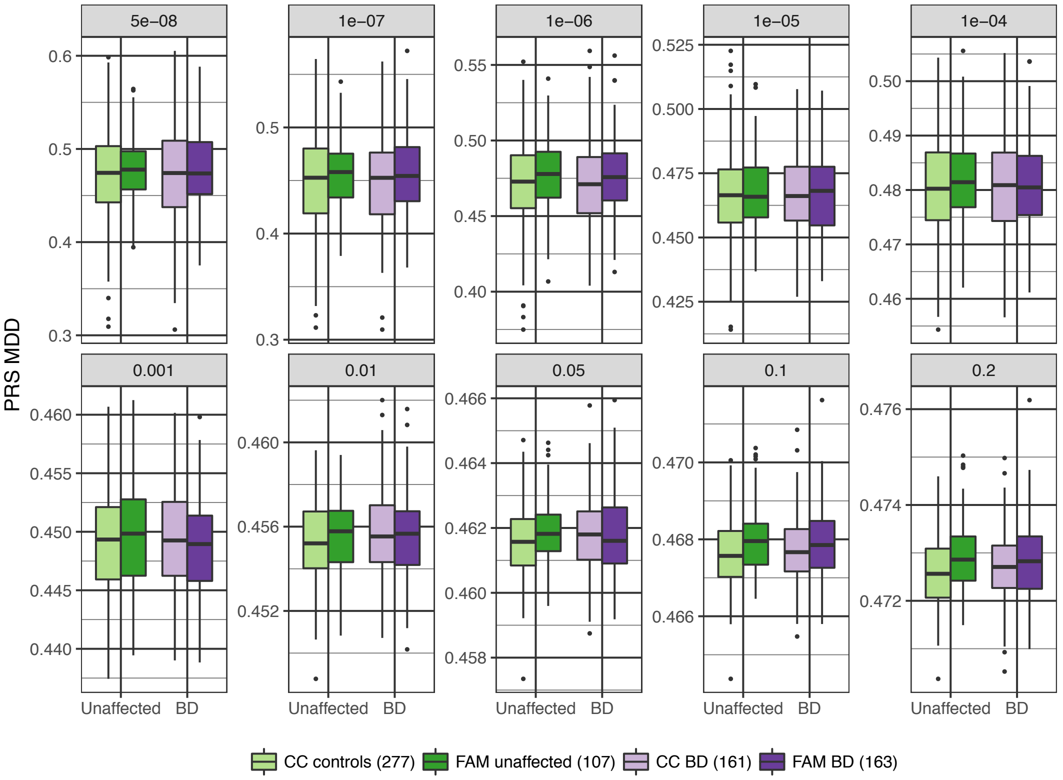


**Supplementary Fig. S1D:** Boxplots of the BD+SCZ+MDD *Shared* PRS. Note that because of the way this PRS was calculated, the maximum possible threshold was $p_{PRS}$=0.1.

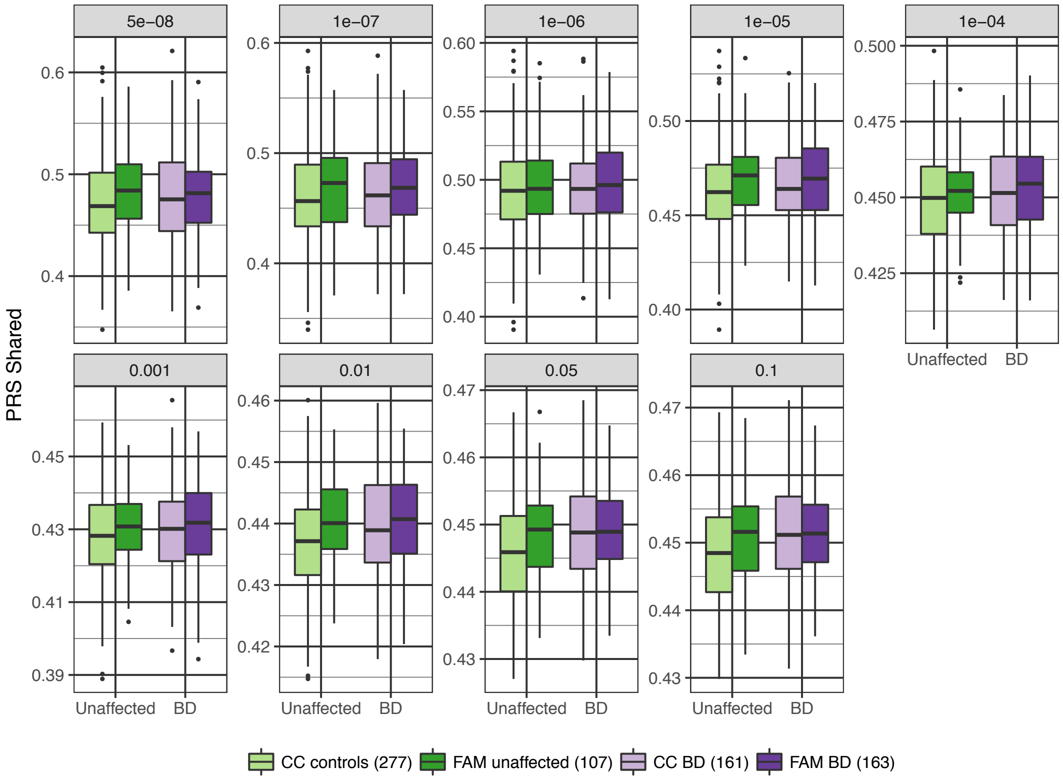

**Supplementary Fig. S1E:** Boxplots of the BD-SCZ GWIS PRS.

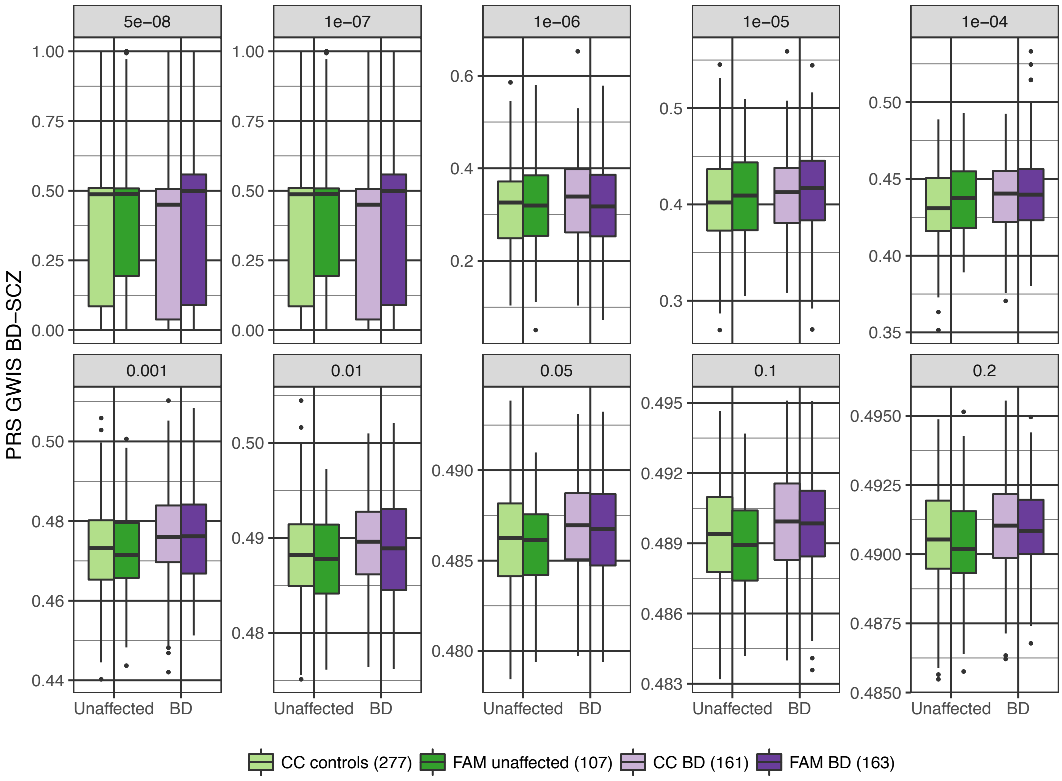


**Supplementary Fig. S1F:** Boxplots of the BD-MDD GWIS PRS.

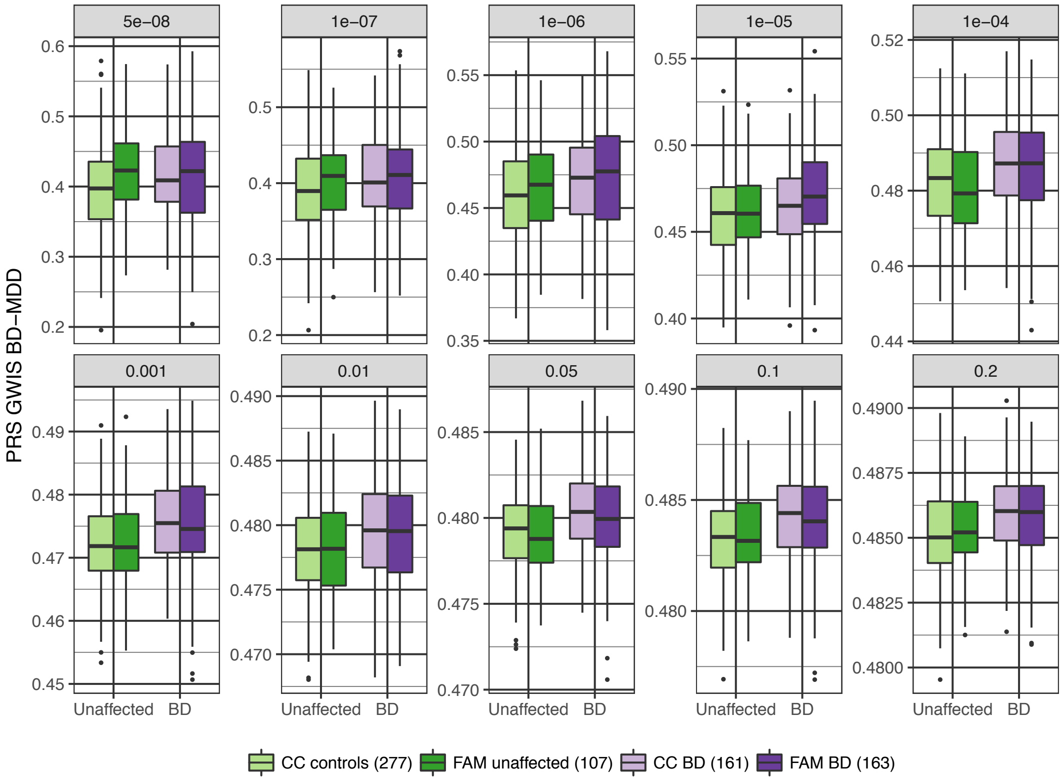

**Supplementary Fig. S1G:** Boxplots of the SCZ-BD GWIS PRS.

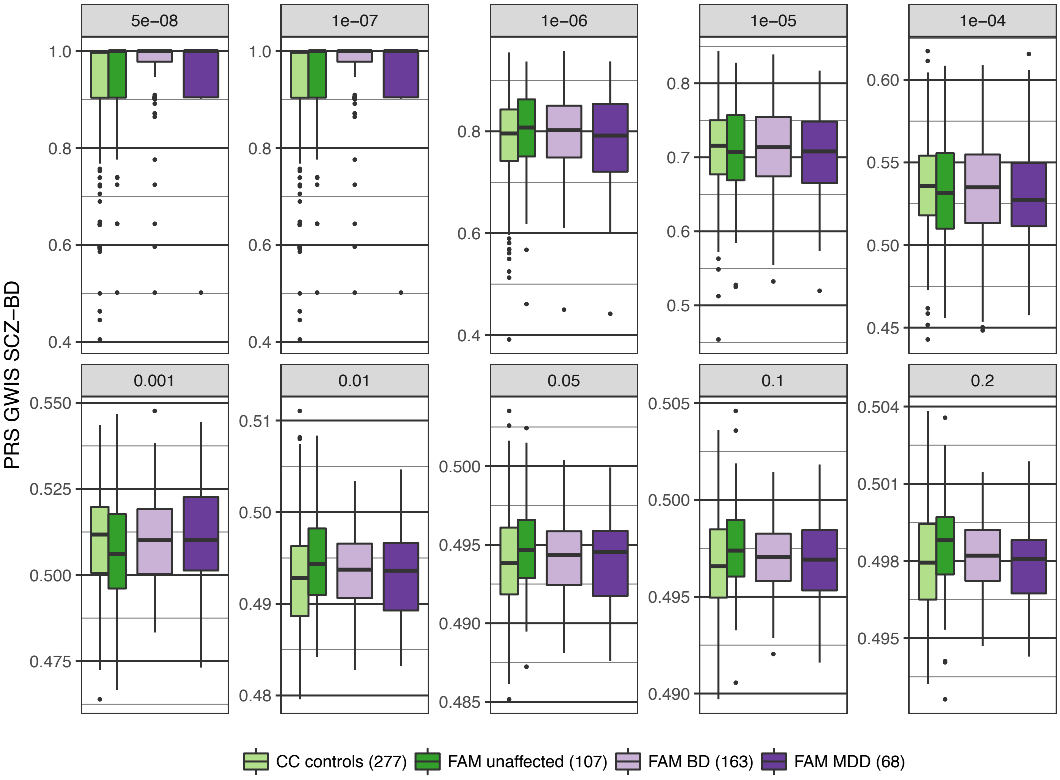


**Supplementary Fig. S1H:** Boxplots of the MDD-BD GWIS PRS.

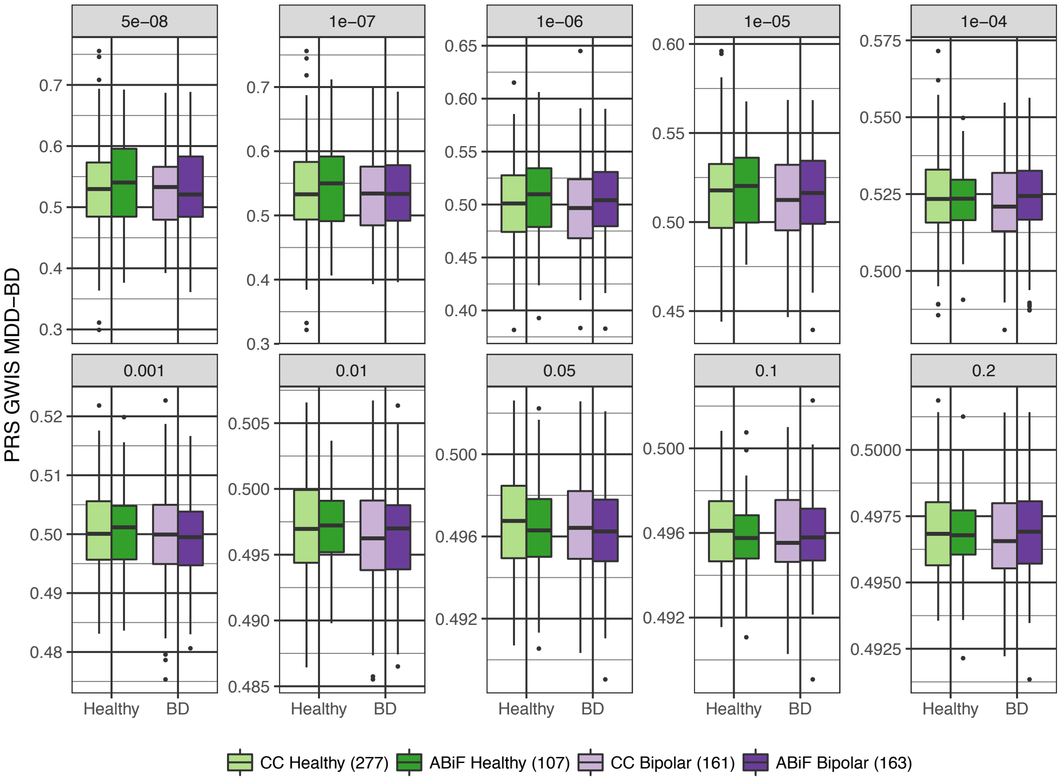

**Supplementary Fig. S1I:** Boxplots of the LOAD (Alzheimer) PRS.

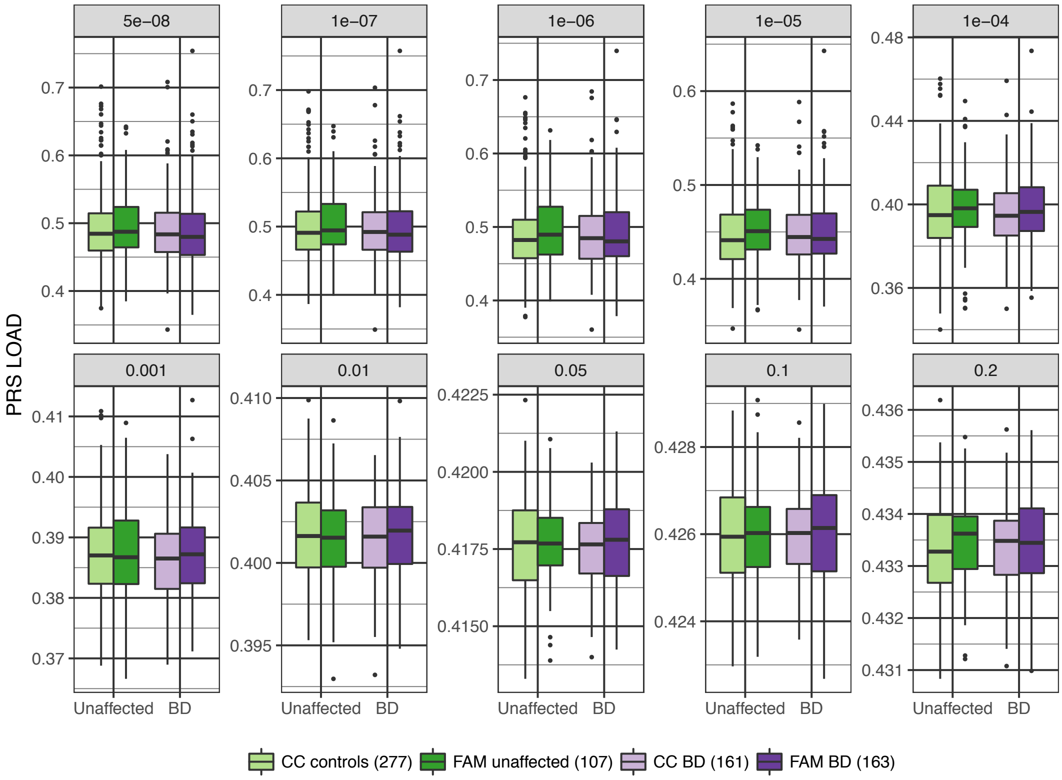


**Supplementary Fig. S2:** Association analysis comparing PRS in $FAM_{BD}$ cases and $CC_{controls}$. Further details of the plots are described in the legend for Fig. 1. Full association test statistics are shown in Supplementary Table S2.

**Supplementary Fig. S2A:** Association of the BD PRS (data is identical to Fig. 1A).

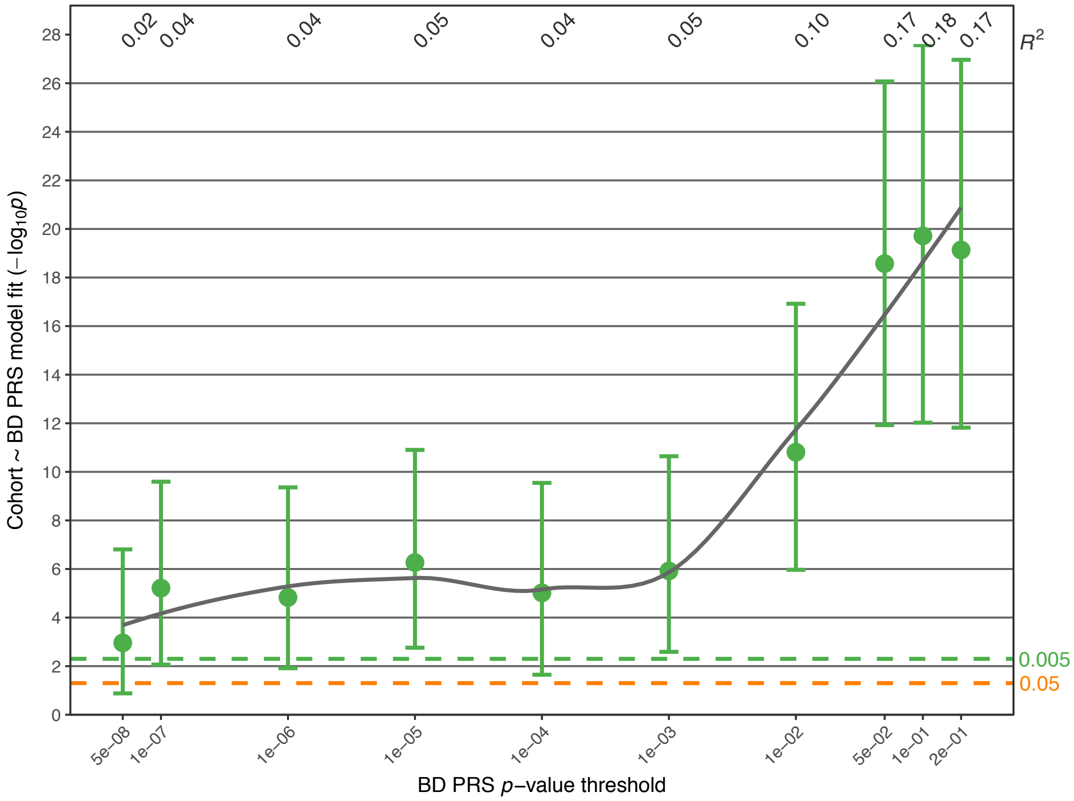

**Supplementary Fig. S2B:** Association of the SCZ PRS.

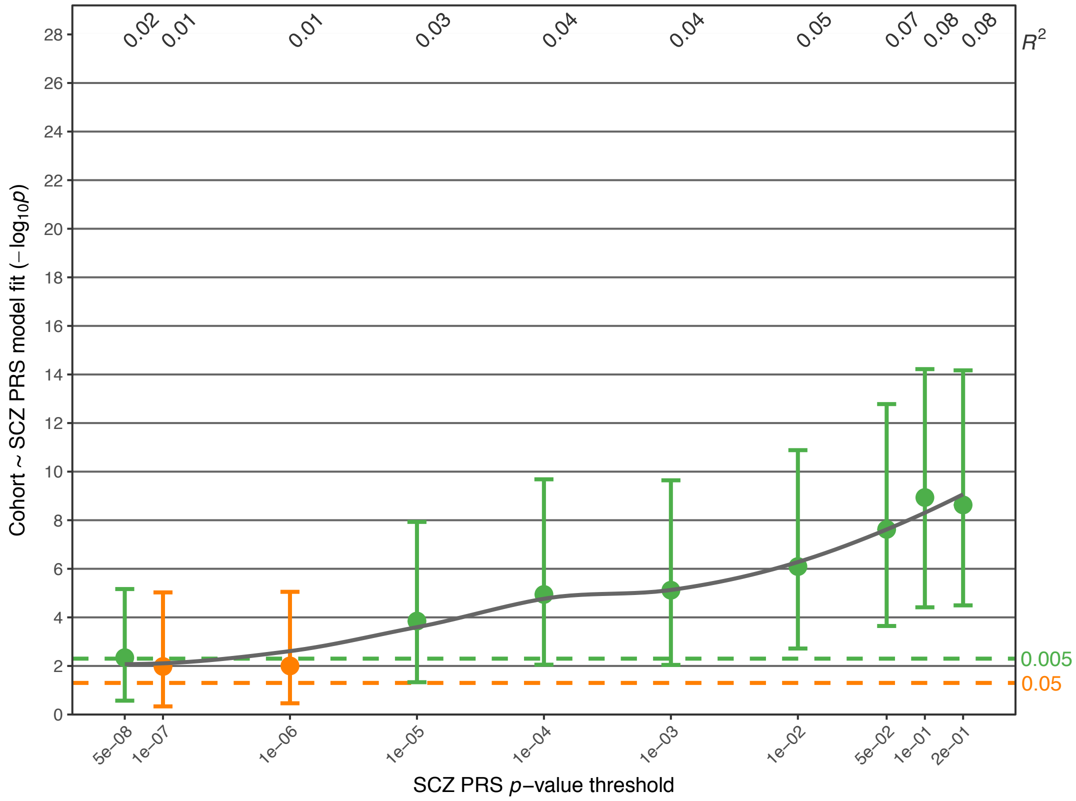


**Supplementary Fig. S2C:** Association of the MDD PRS.

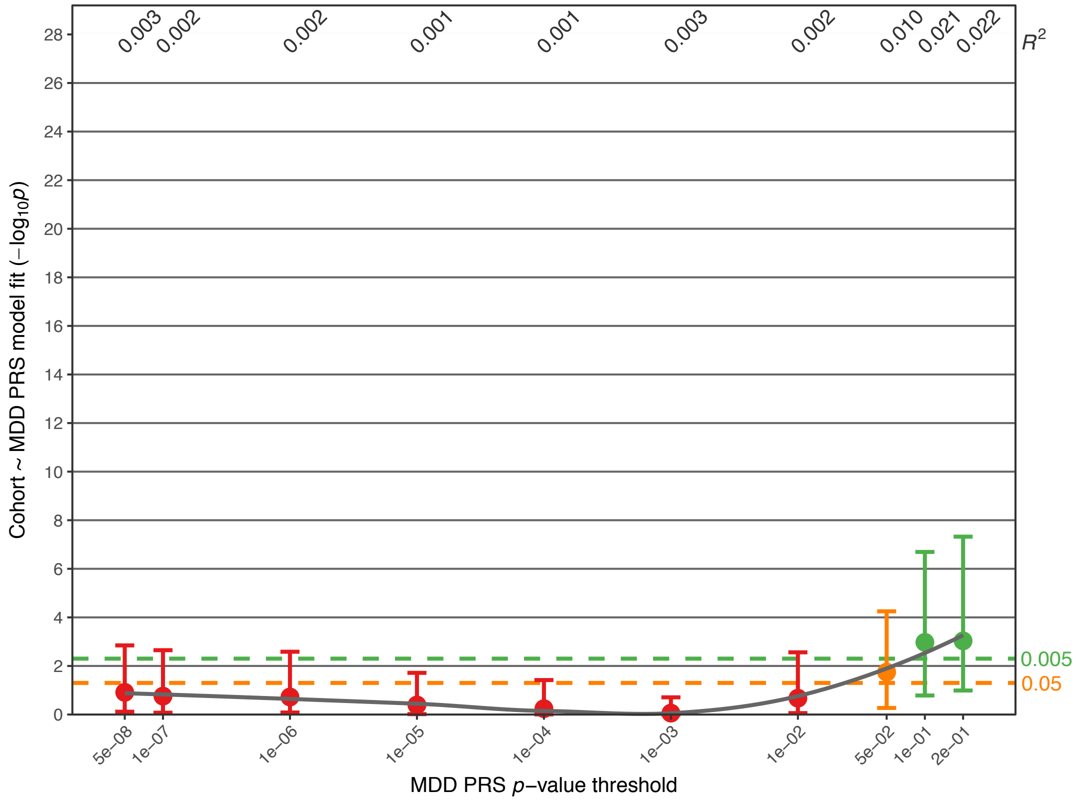

**Supplementary Fig. S2D:** Association of the *Shared* PRS.

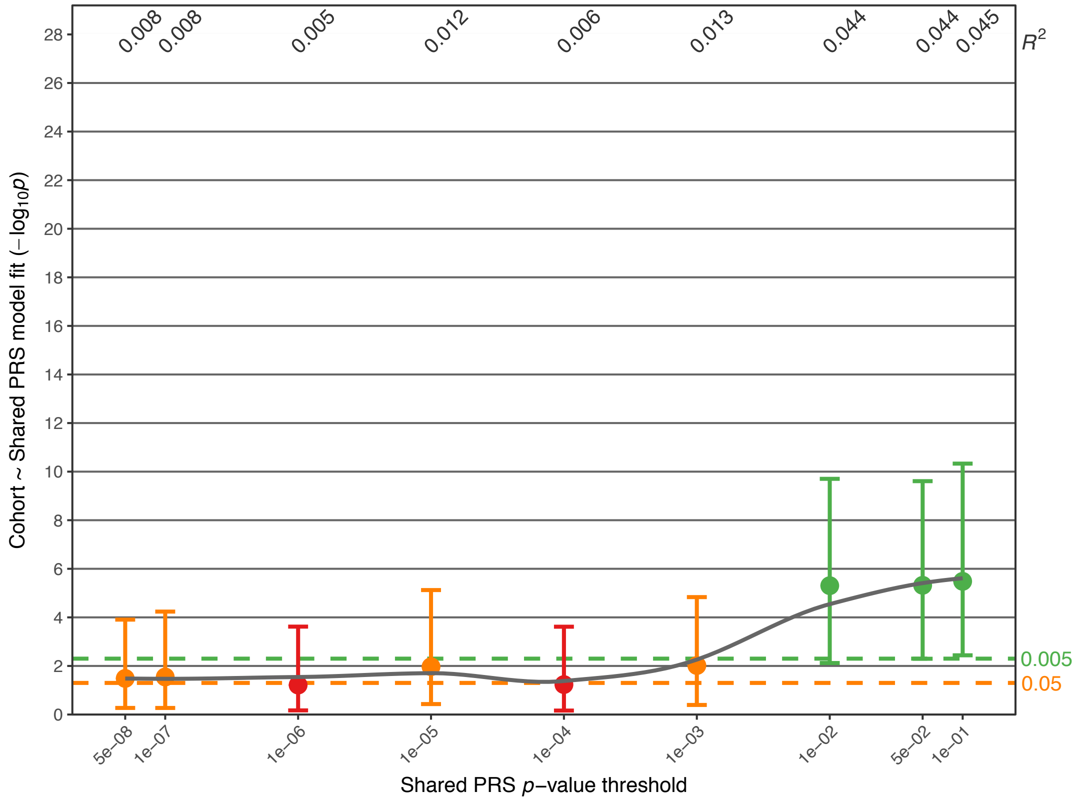


**Supplementary Fig. S2E:** Association of the BD-SCZ GWIS PRS.

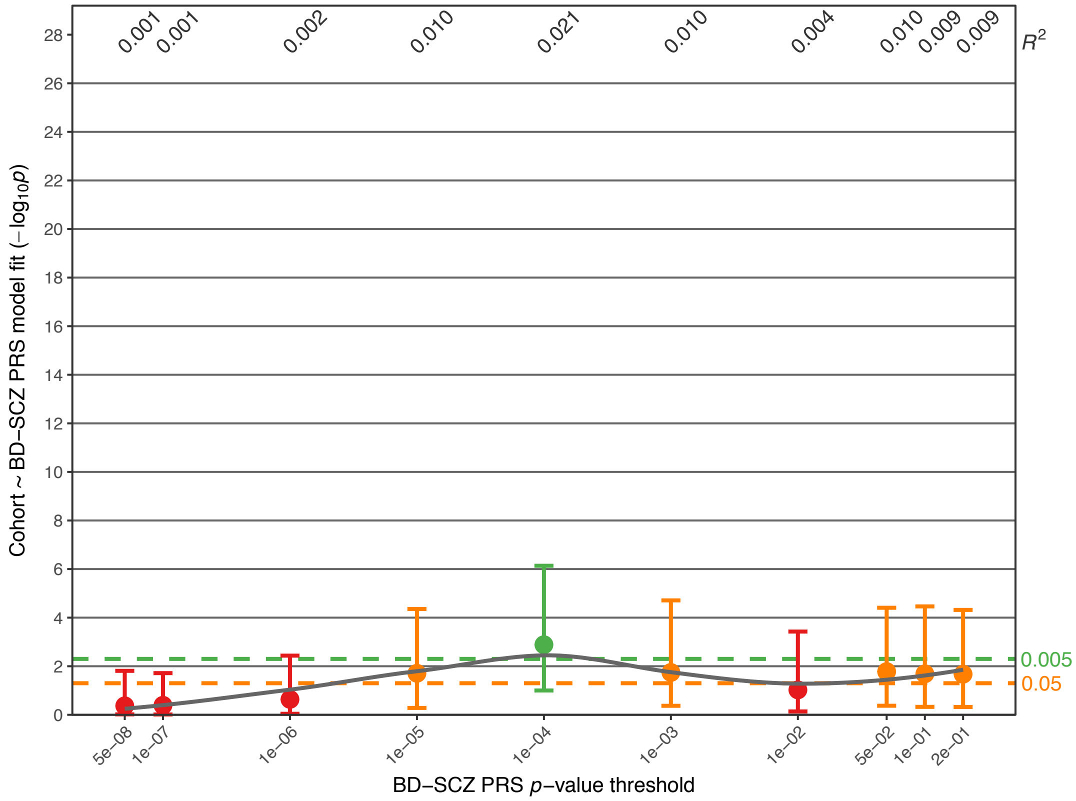

**Supplementary Fig. S2F:** Association of the BD-MDD GWIS PRS.

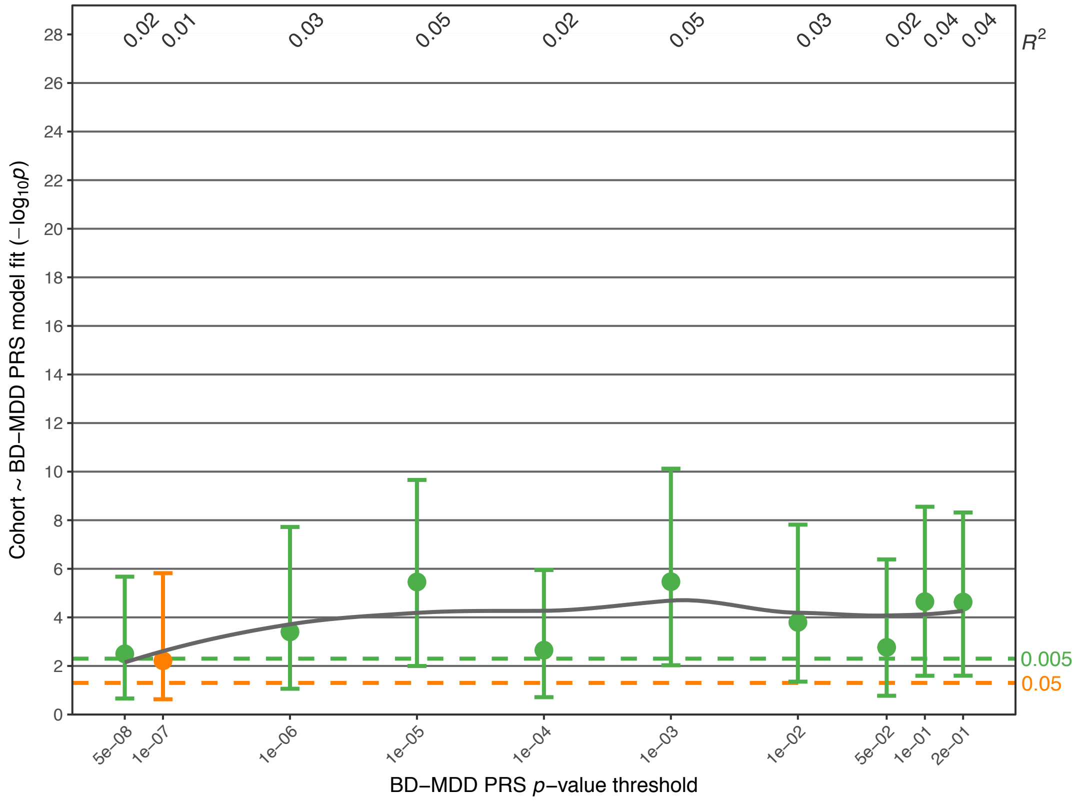


**Supplementary Fig. S2G:** Association of the SCZ-BD GWIS PRS.

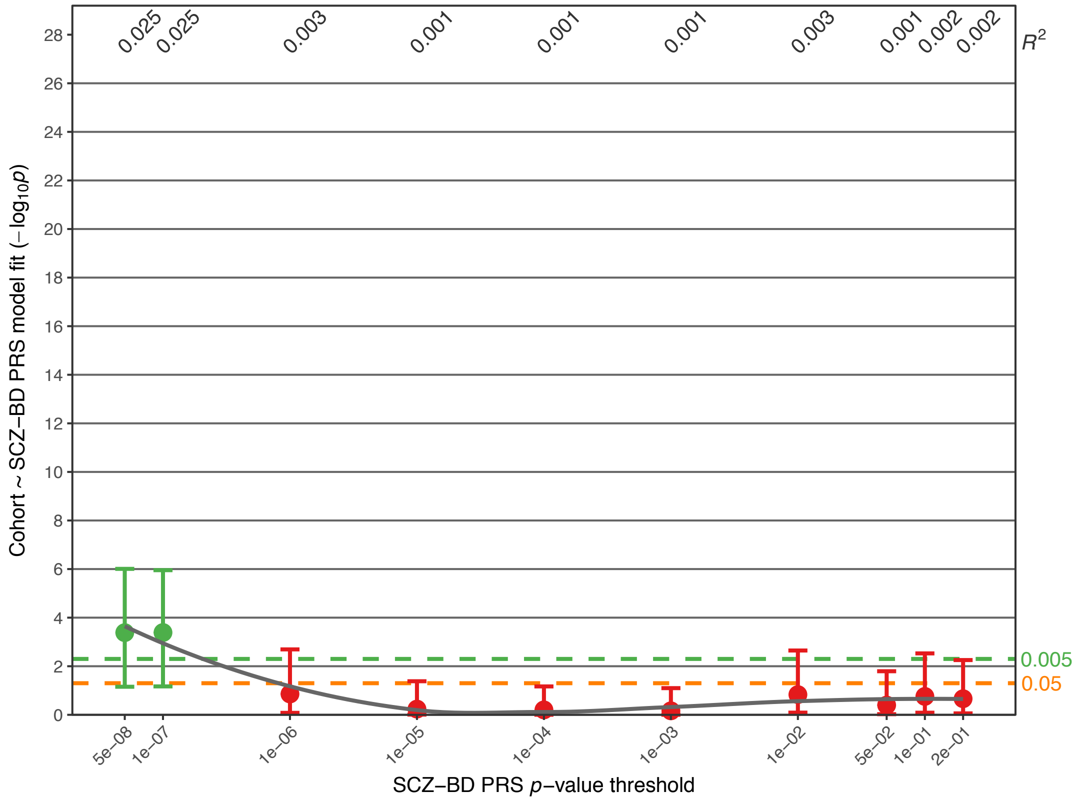

**Supplementary Fig. S2H:** Association of the MDD-BD GWIS PRS.

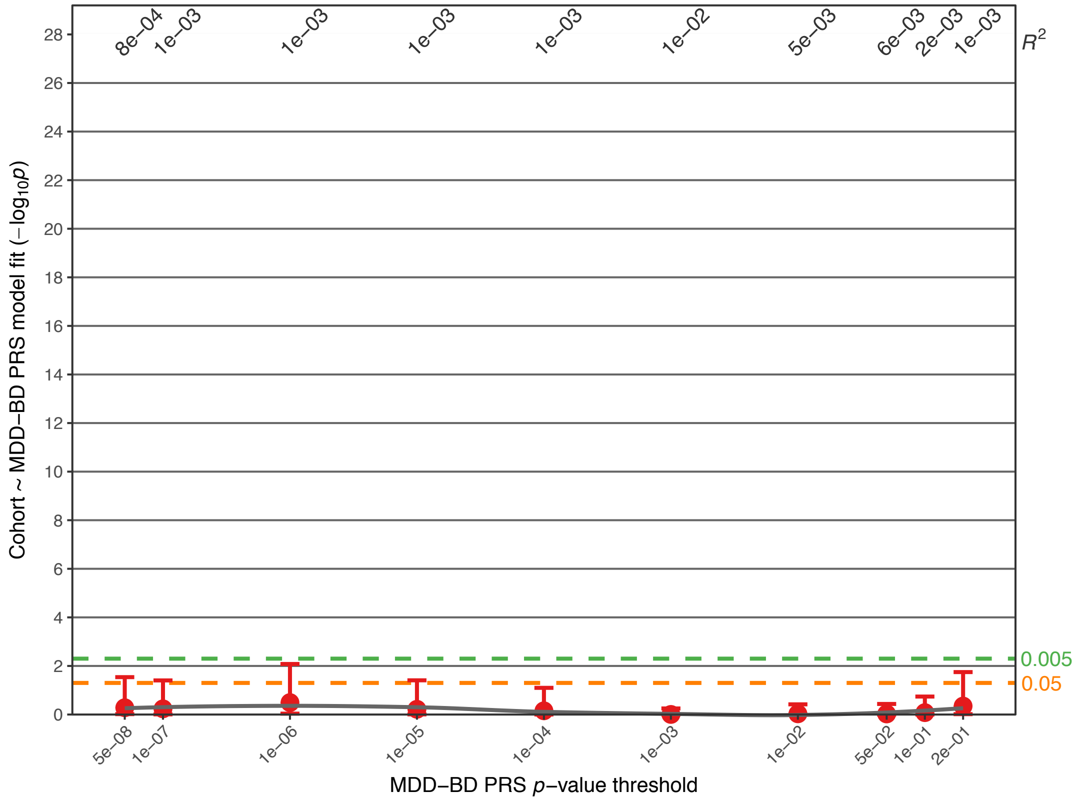


**Supplementary Fig. S2I:** Association of the LOAD PRS.

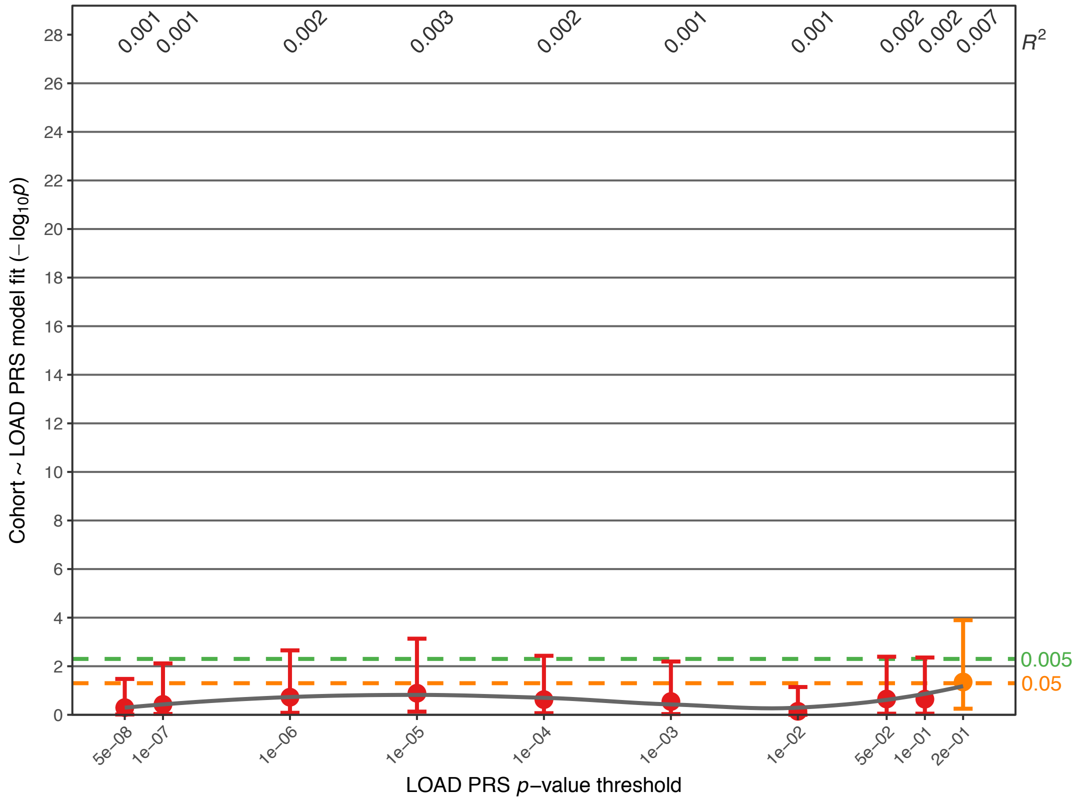

**Supplementary Fig. S3:** Association analysis comparing PRS in $FAM_{BD}$ cases and sporadic $CC_{BD}$ cases. Further details of the plots are described in the legend for Fig. 1. Full association test statistics are shown in Supplementary Table S3.

**Supplementary Fig. S3A:** Association of the BD PRS (data is identical to Fig. 1C).

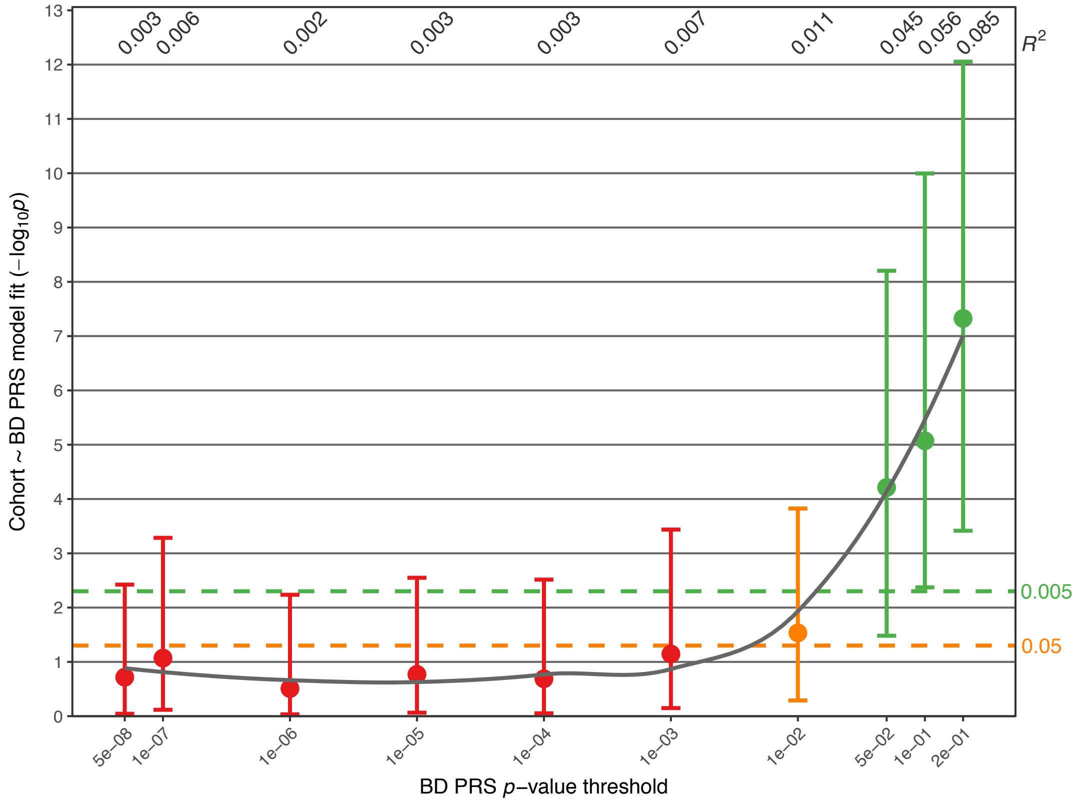


**Supplementary Fig. S3B:** Association of the SCZ PRS.

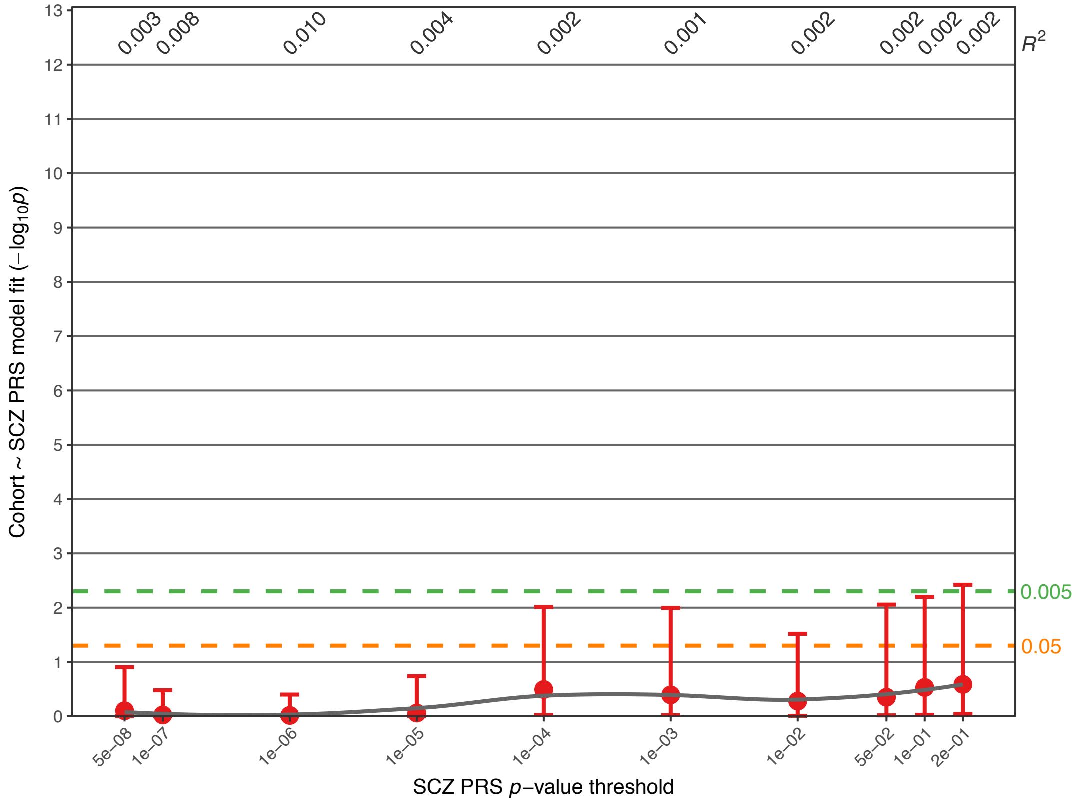

**Supplementary Fig. S3C:** Association of the MDD PRS.

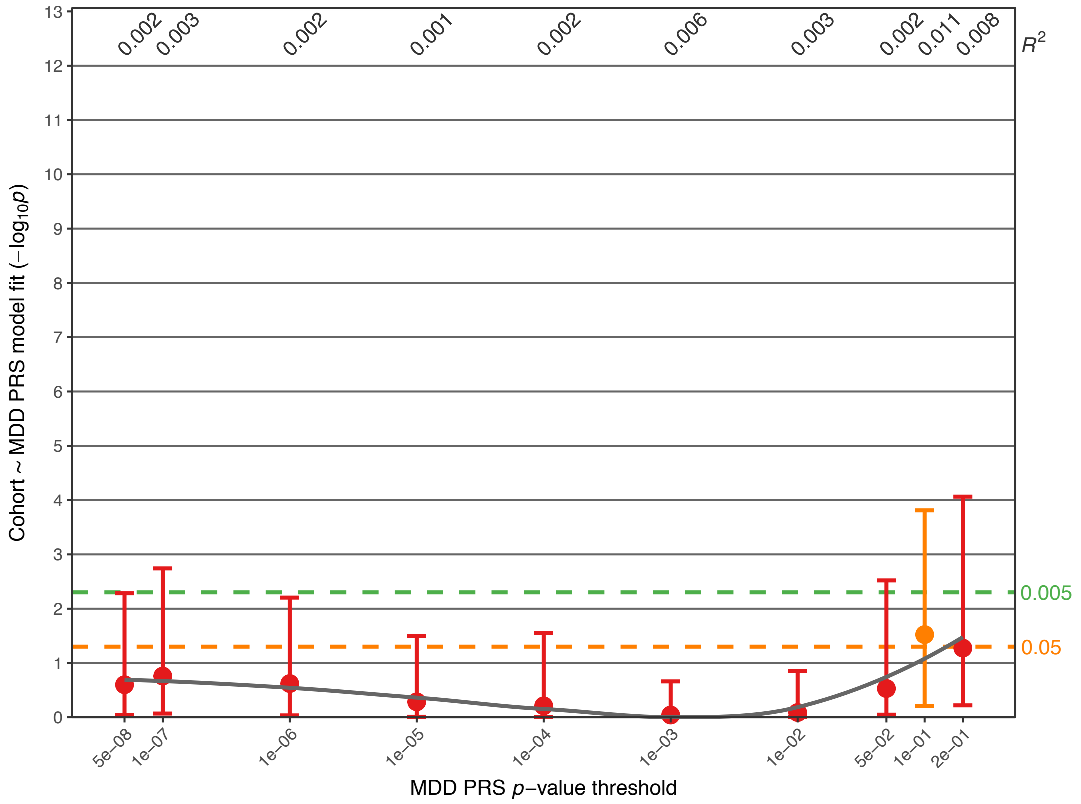


**Supplementary Fig. S3D:** Association of the *Shared* PRS.

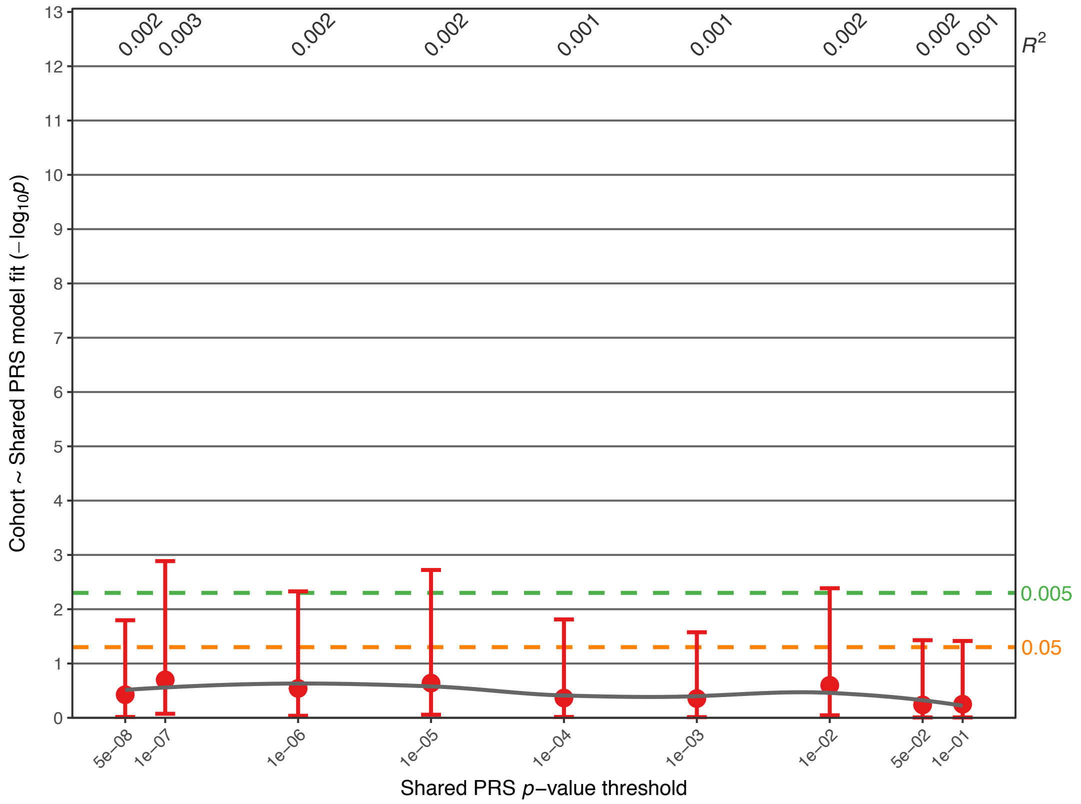

**Supplementary Fig. S3E:** Association of the BD-SCZ GWIS PRS.

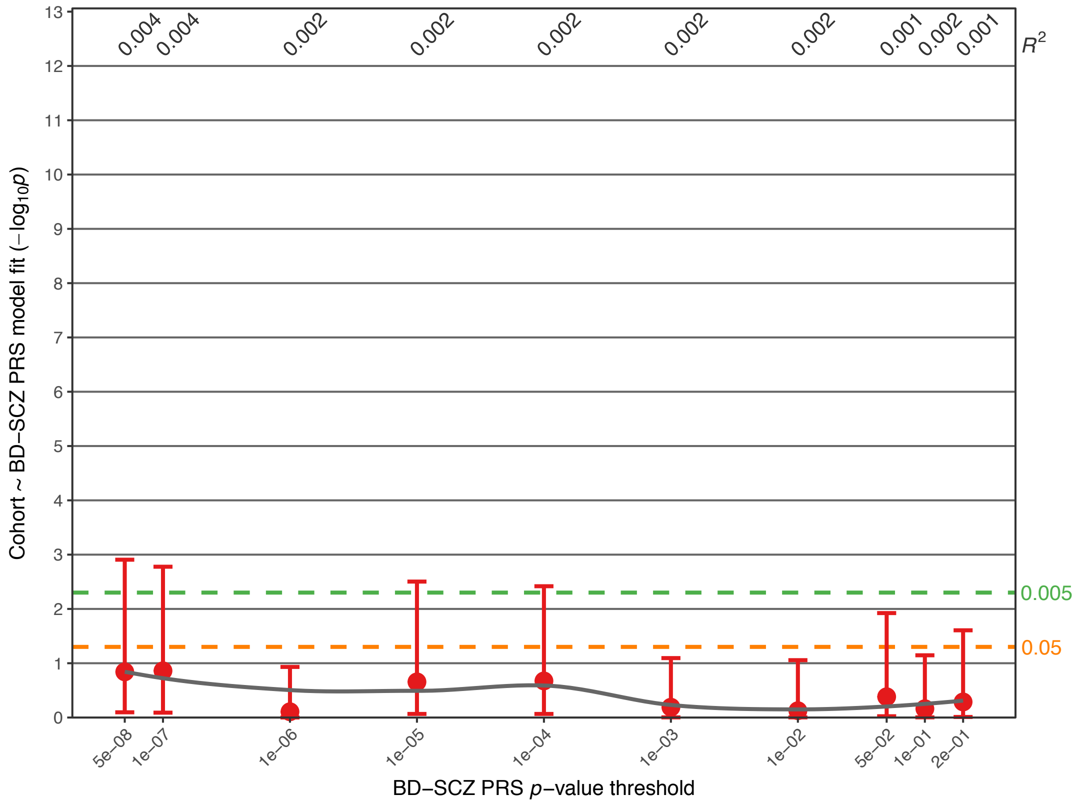


**Supplementary Fig. S3F:** Association of the BD-MDD GWIS PRS.

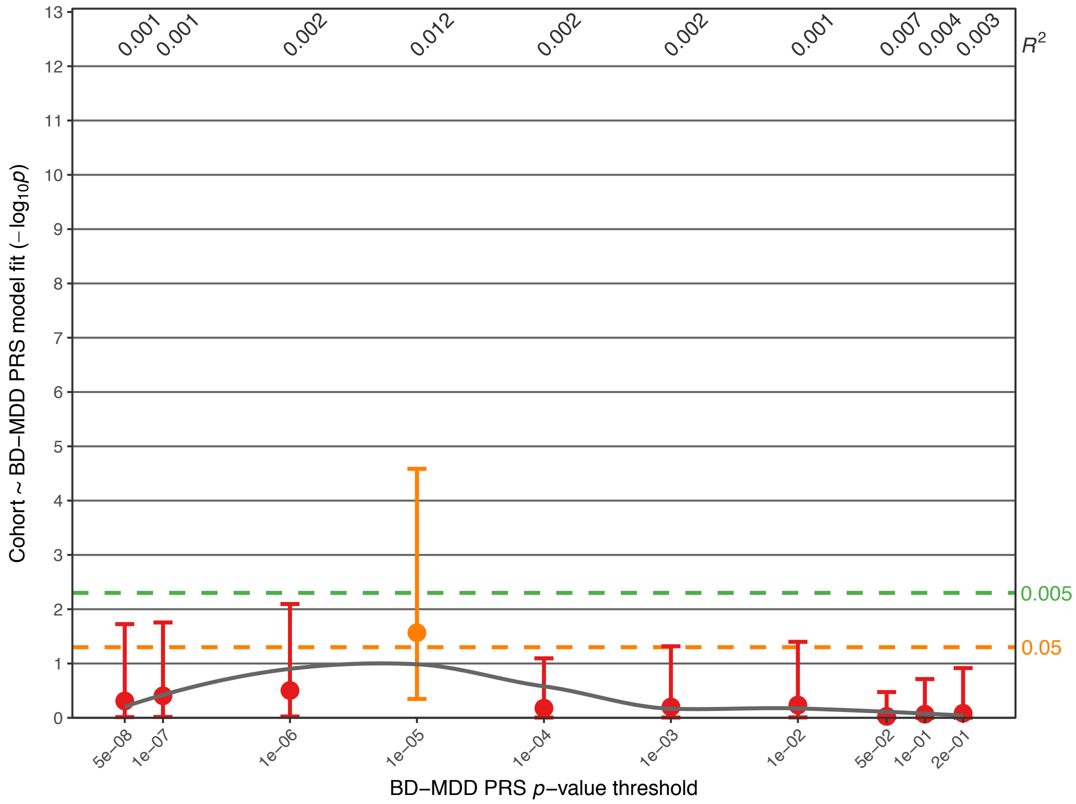

**Supplementary Fig. S3G:** Association of the SCZ-BD GWIS PRS.

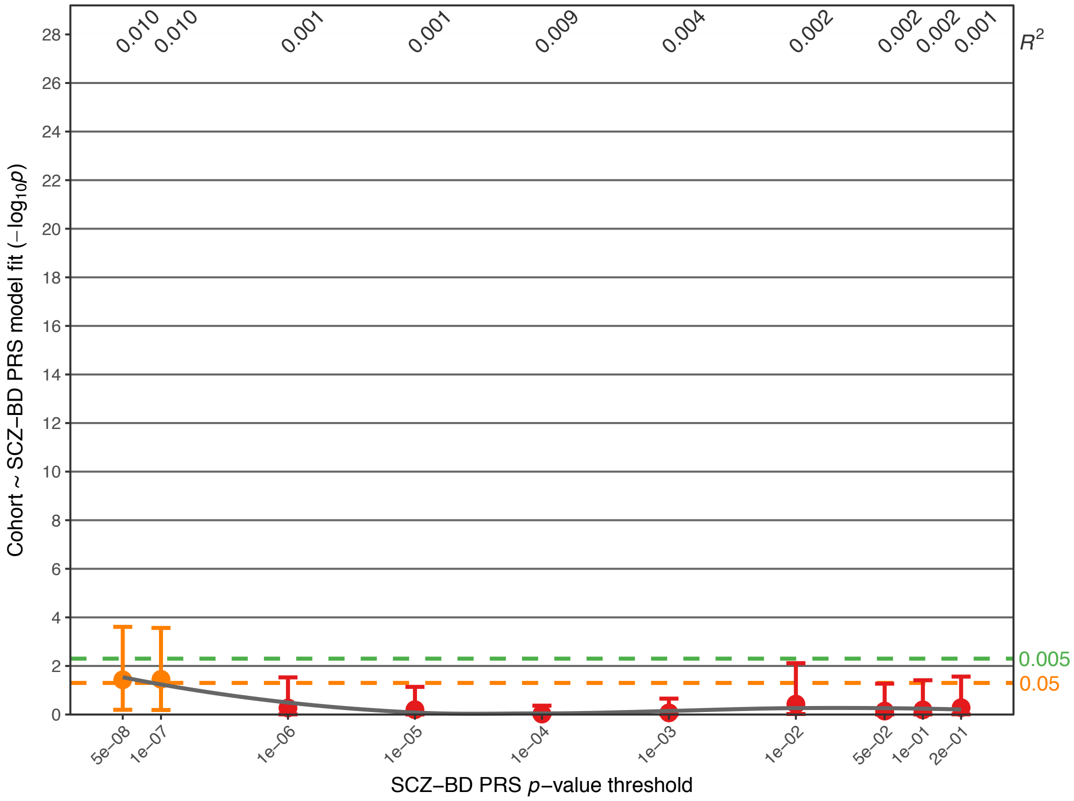


**Supplementary Fig. S3H:** Association of the MDD-BD GWIS PRS.

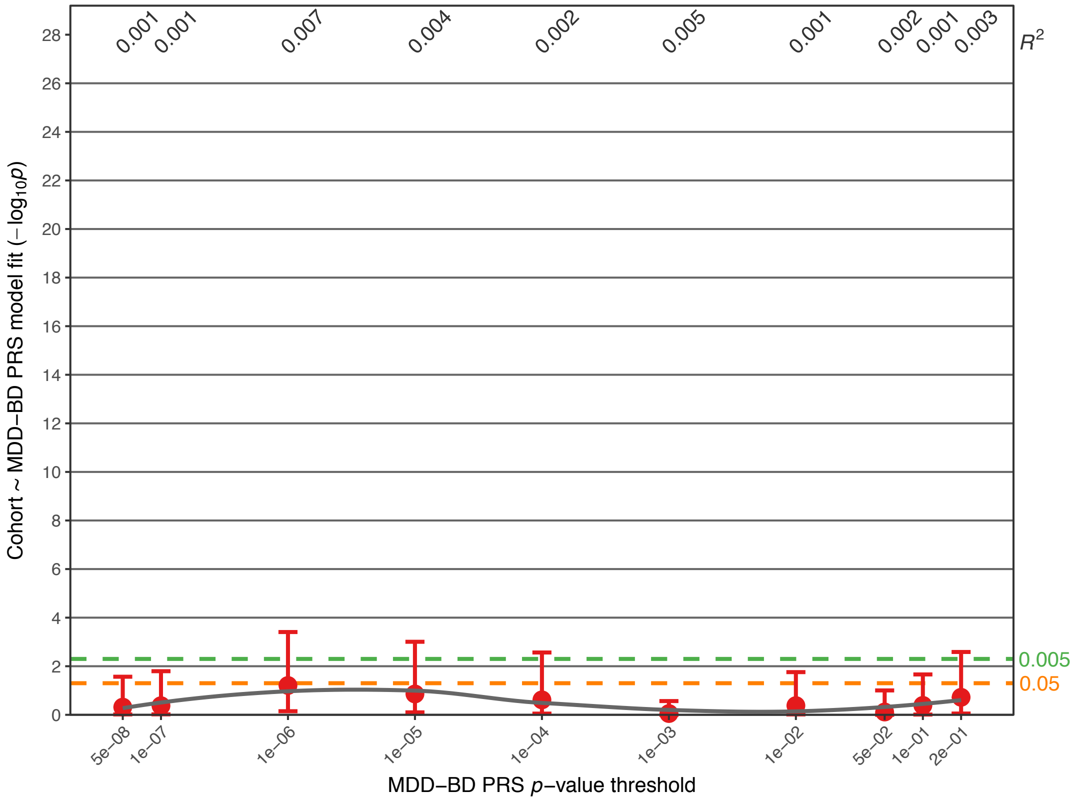

**Supplementary Fig. S3I:** Association of the LOAD PRS.

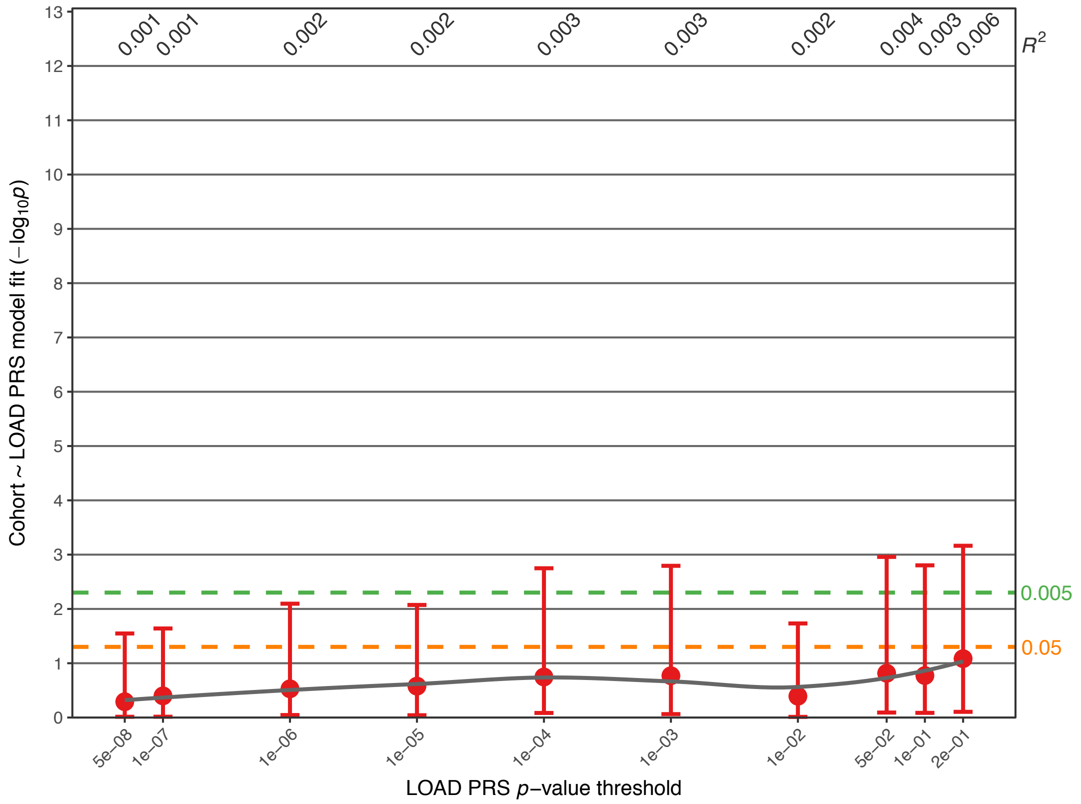


**Supplementary Fig. S4:** Association analysis comparing PRS in $FAM_{unaffected}$ and $CC_{controls}$. Further details of the plots are described in the legend for Fig. 1. Full association test statistics are shown in Supplementary Table S4.

**Supplementary Fig. S4A:** Association of the BD PRS (data is identical to Fig. 1E).

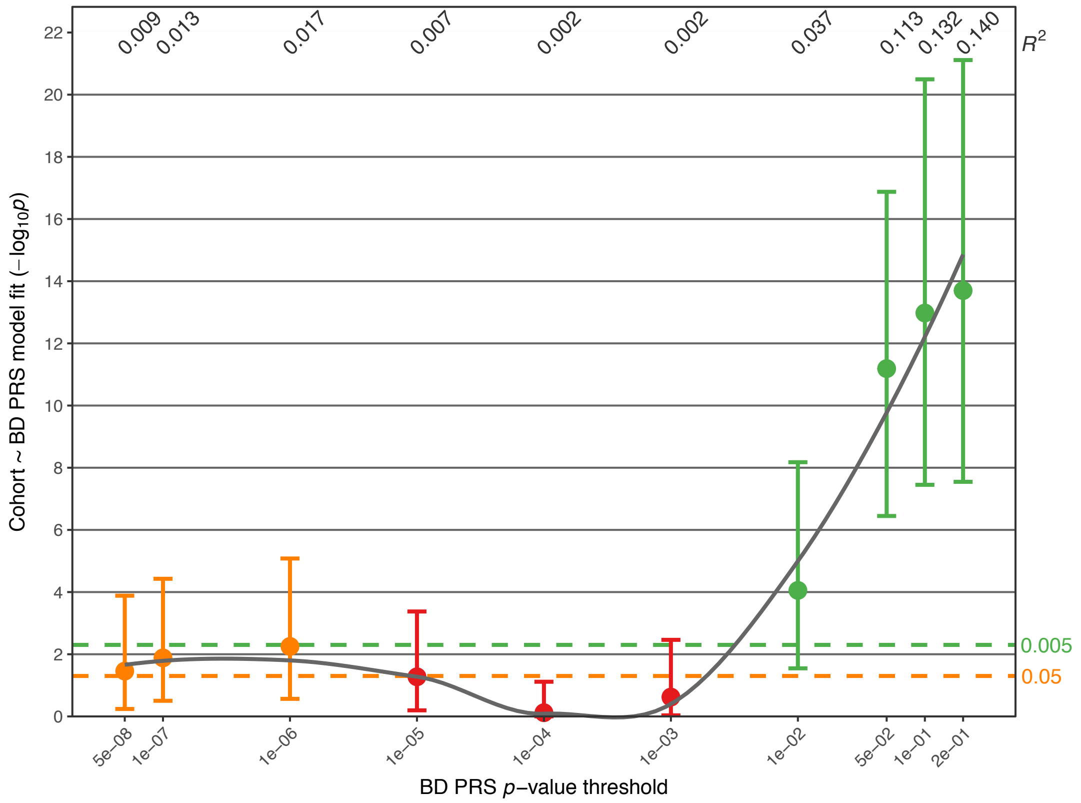

**Supplementary Fig. S4B:** Association of the SCZ PRS.

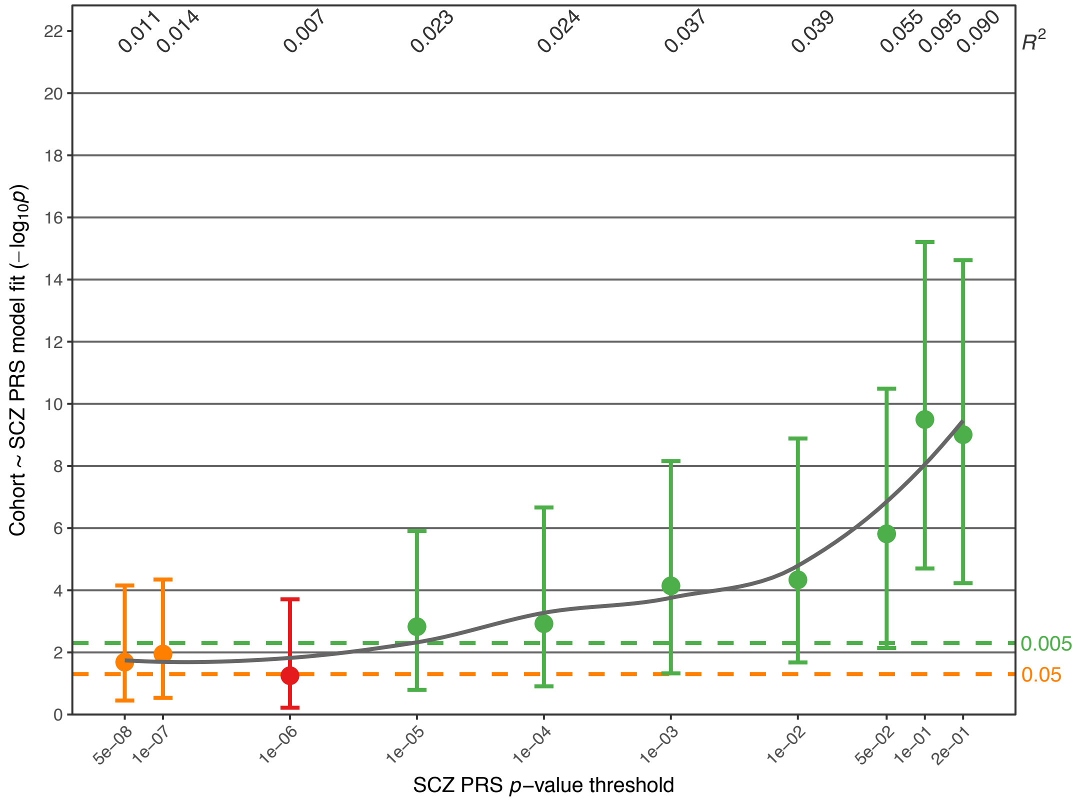


**Supplementary Fig. S4C:** Association of the MDD PRS.

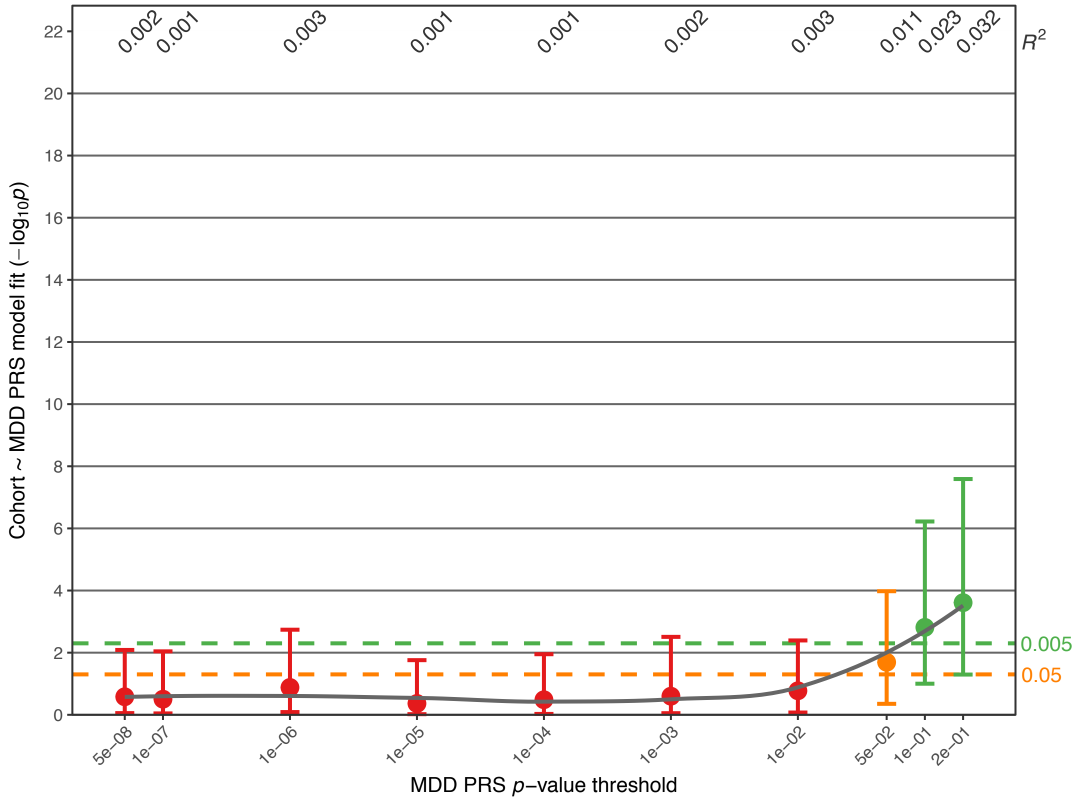

**Supplementary Fig. S4D:** Association of the *Shared* PRS.

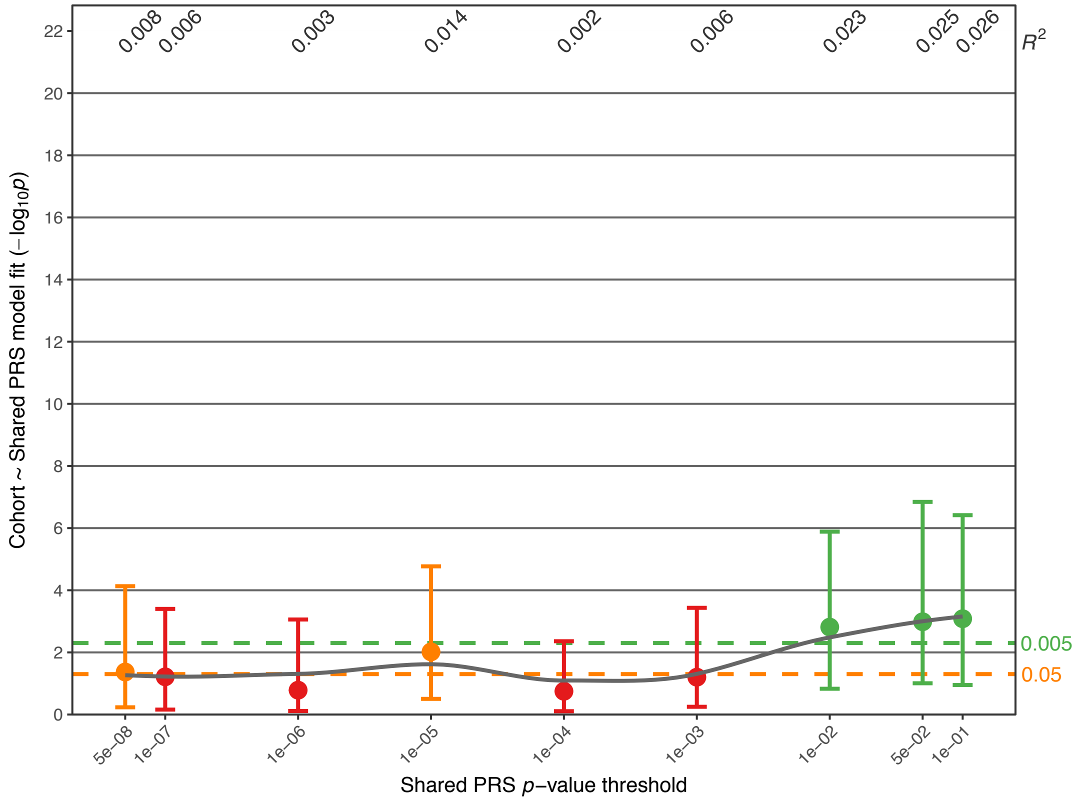


**Supplementary Fig. S4E:** Association of the BD-SCZ GWIS PRS.

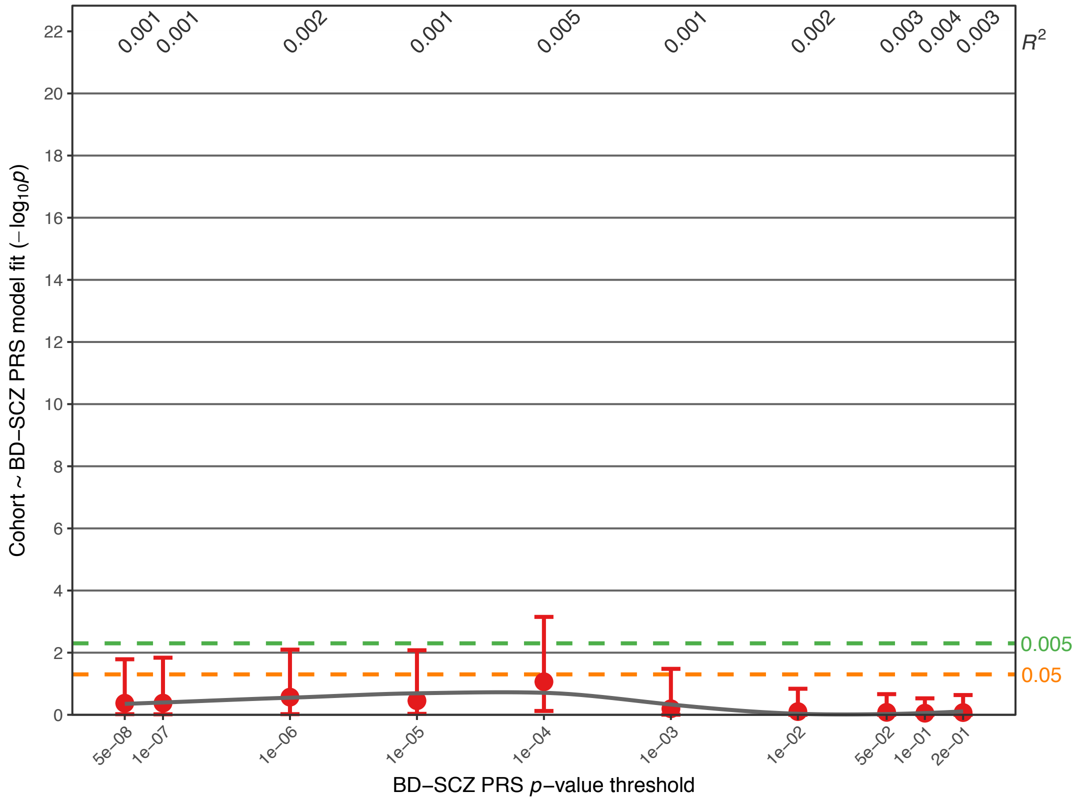

**Supplementary Fig. S4F:** Association of the BD-MDD GWIS PRS.

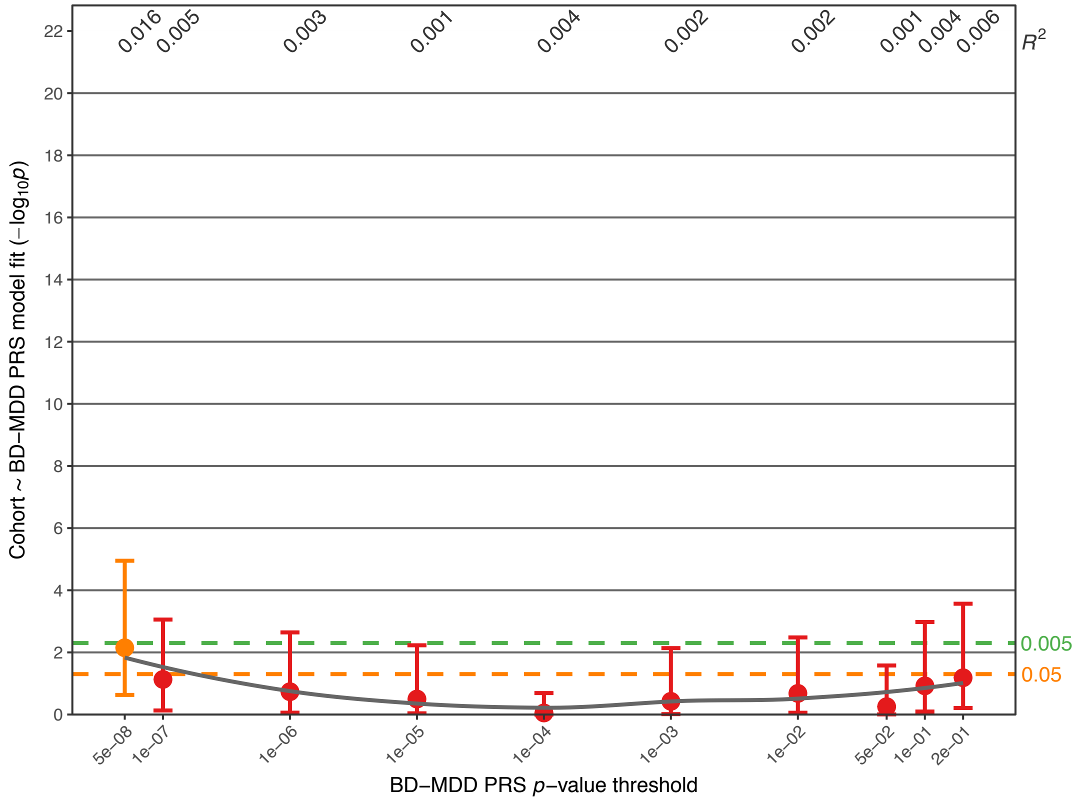


**Supplementary Fig. S4G:** Association of the SCZ-BD GWIS PRS.

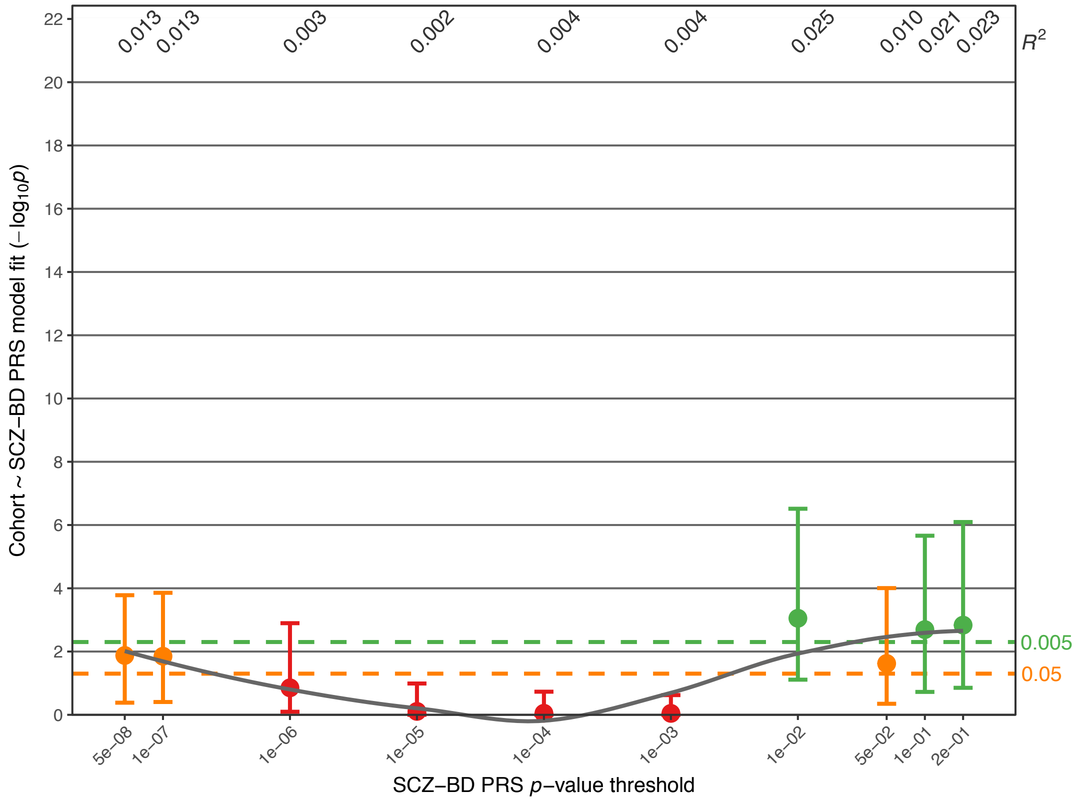

**Supplementary Fig. S4H:** Association of the MDD-BD GWIS PRS.

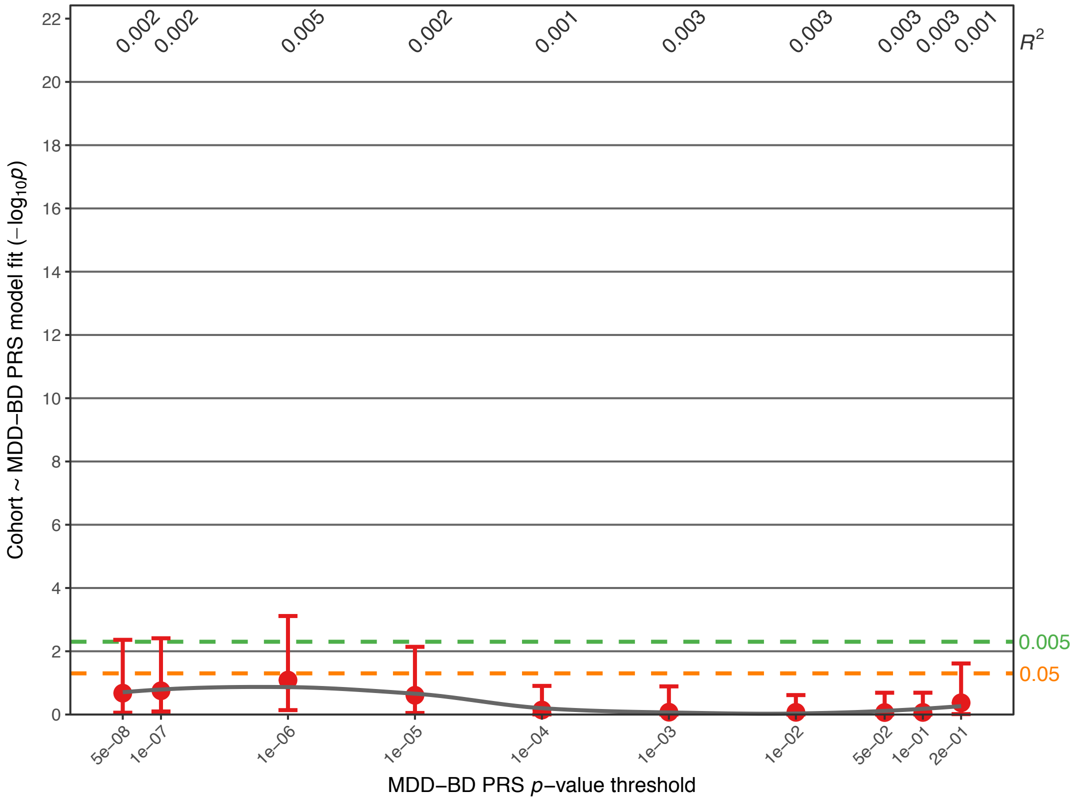


**Supplementary Fig. S4I:** Association of the LOAD PRS.

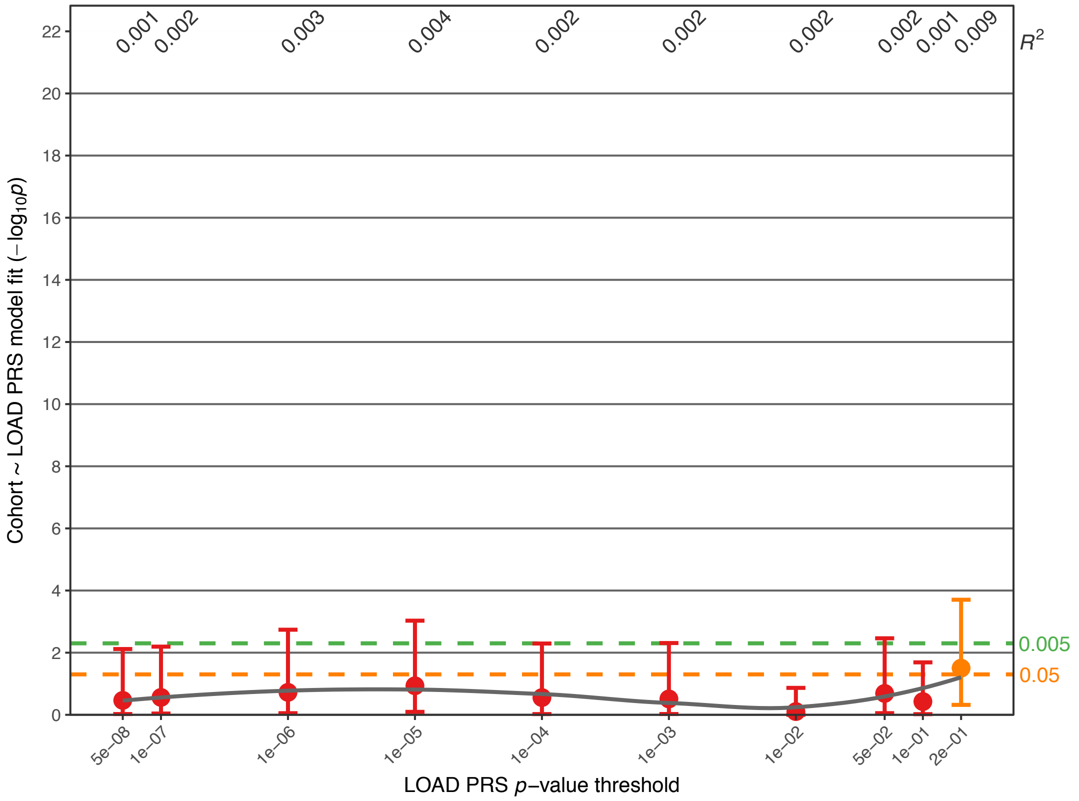

**Supplementary Fig. S5:** Association analysis comparing PRS in FAM$_{BD}$ cases and FAM$_{unaffected}$. Further details of the plots are described in the legends for Figs. 1 and 2. Full association test statistics are shown in Supplementary Table S5.

**Supplementary Fig. S5A:** Association of the BD PRS (data is identical to Fig. 2A).

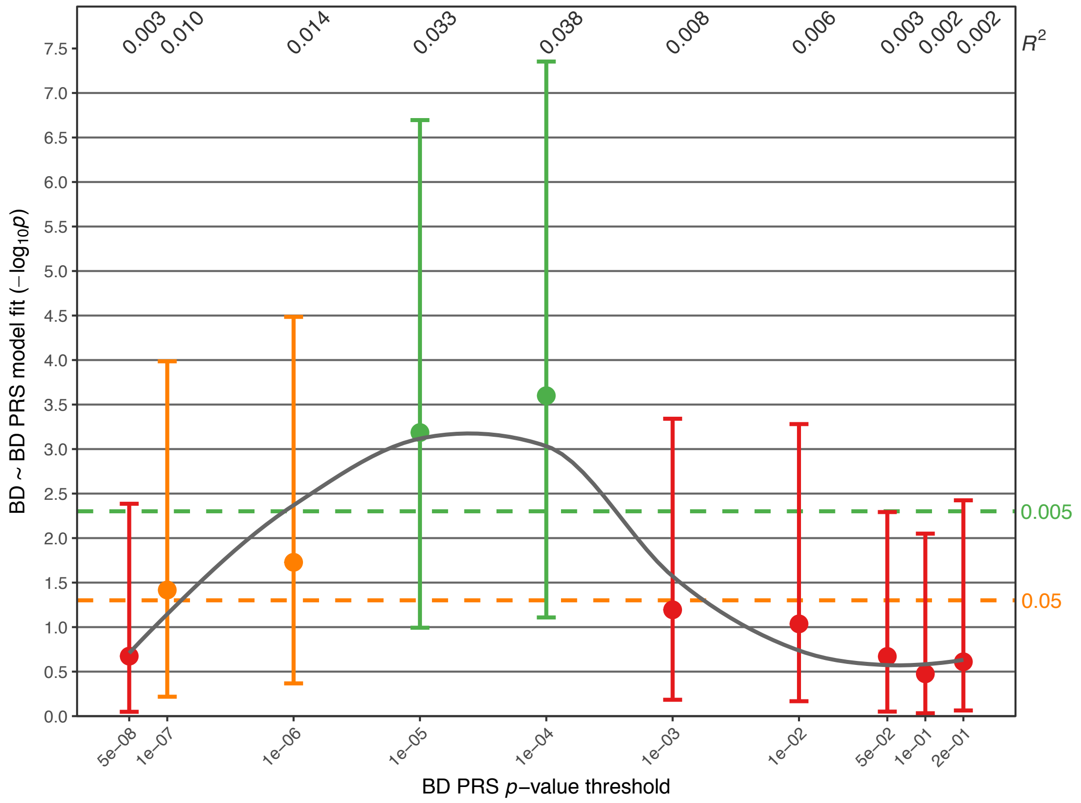


**Supplementary Fig. S5B:** Association of the SCZ PRS.

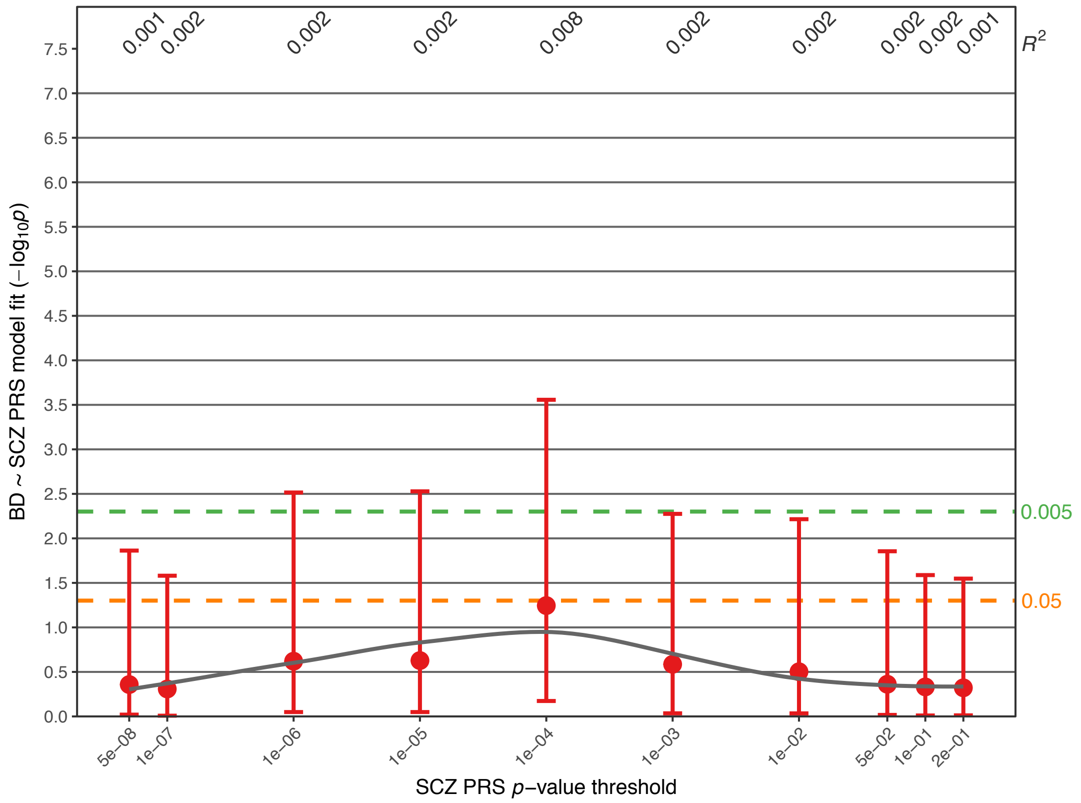

**Supplementary Fig. S5C:** Association of the MDD PRS.

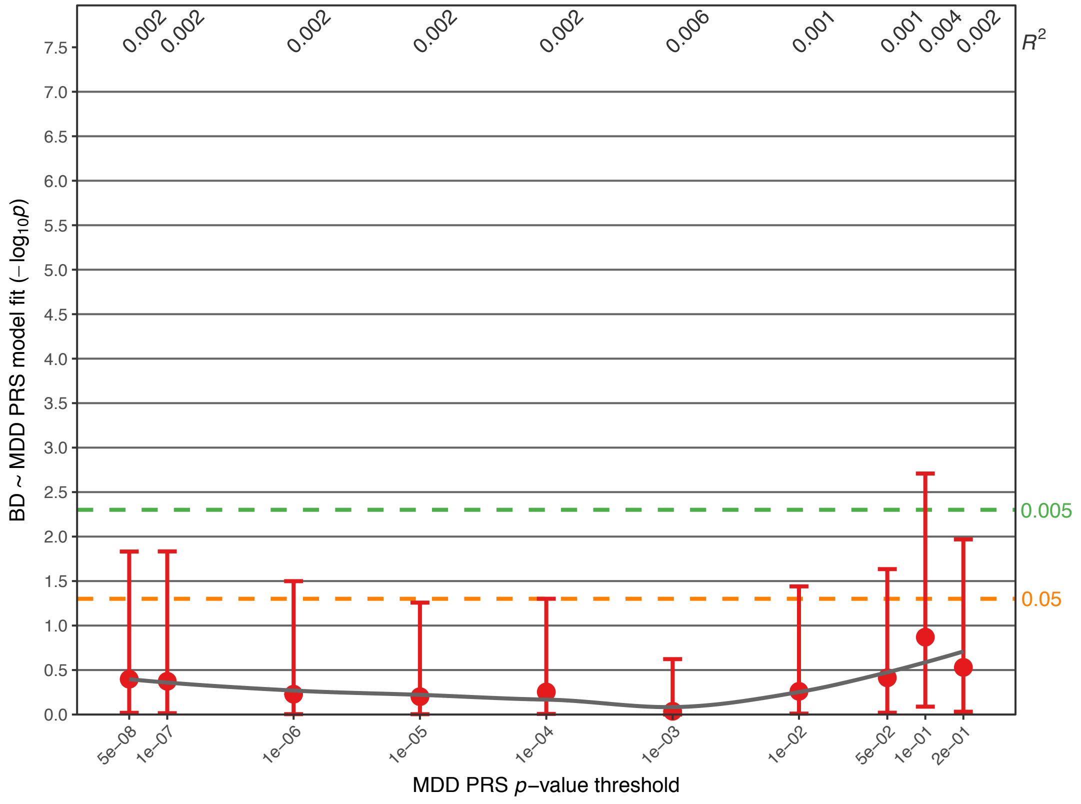


**Supplementary Fig. S5D:** Association of the *Shared* PRS.

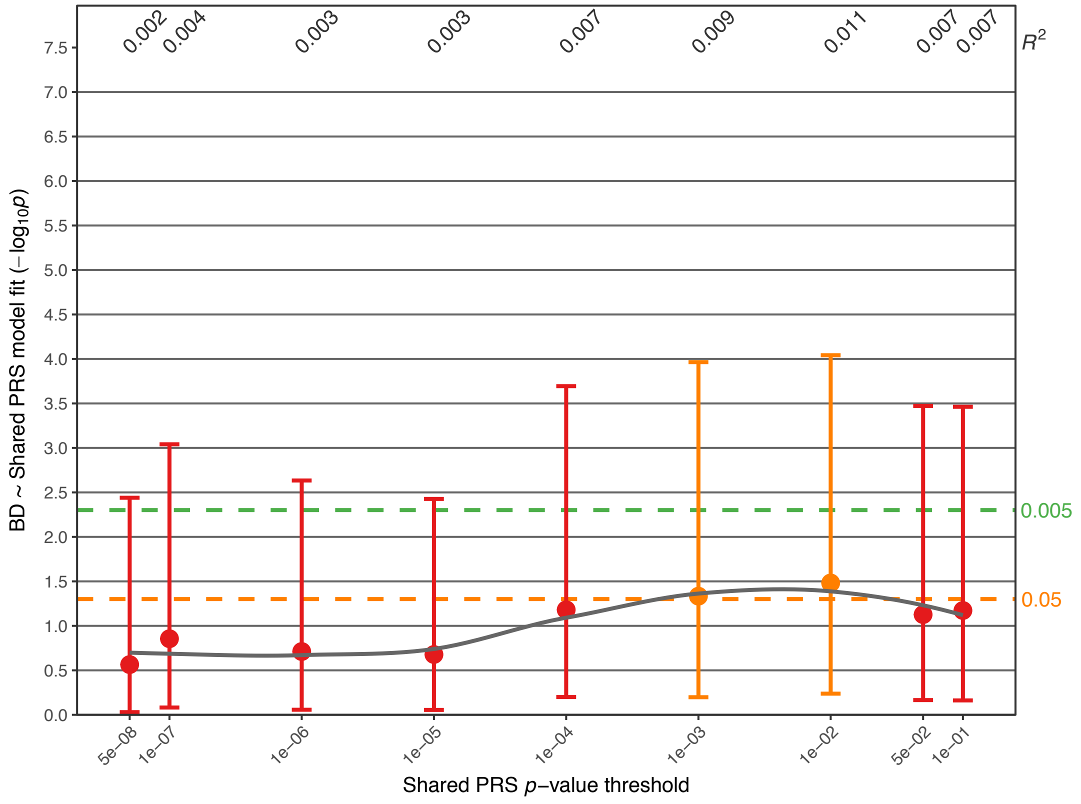

**Supplementary Fig. S5E:** Association of the BD-SCZ GWIS PRS.

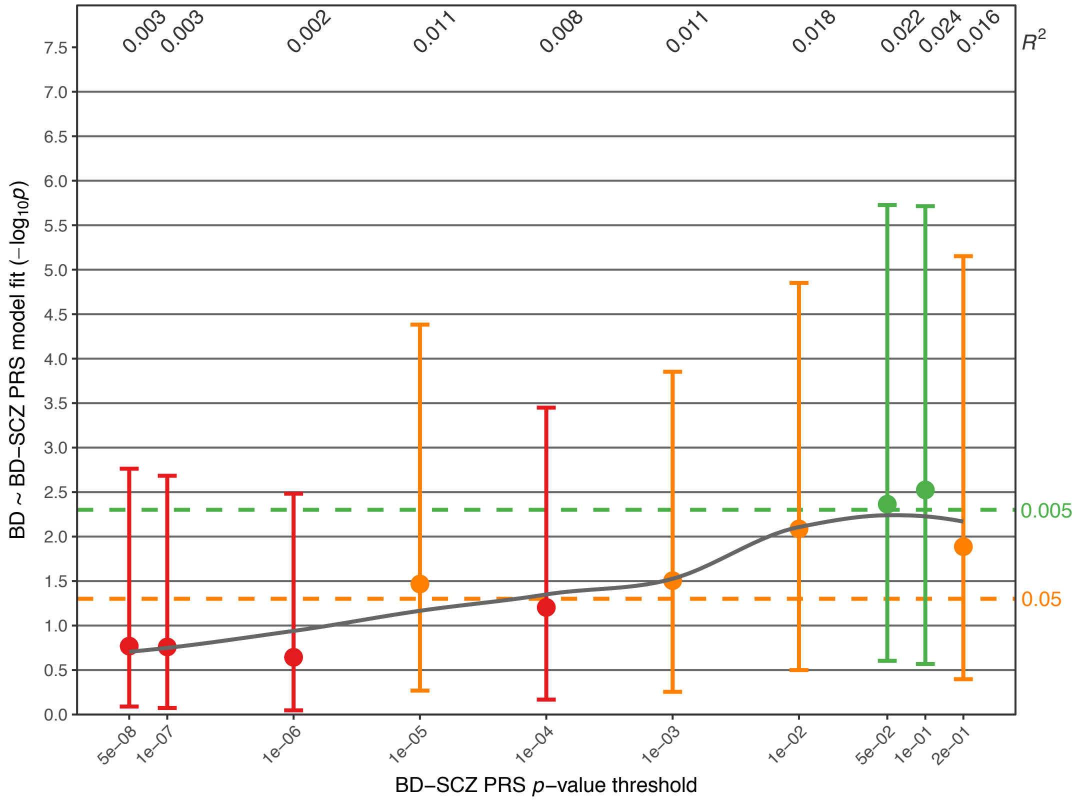


**Supplementary Fig. S5F:** Association of the BD-MDD GWIS PRS.

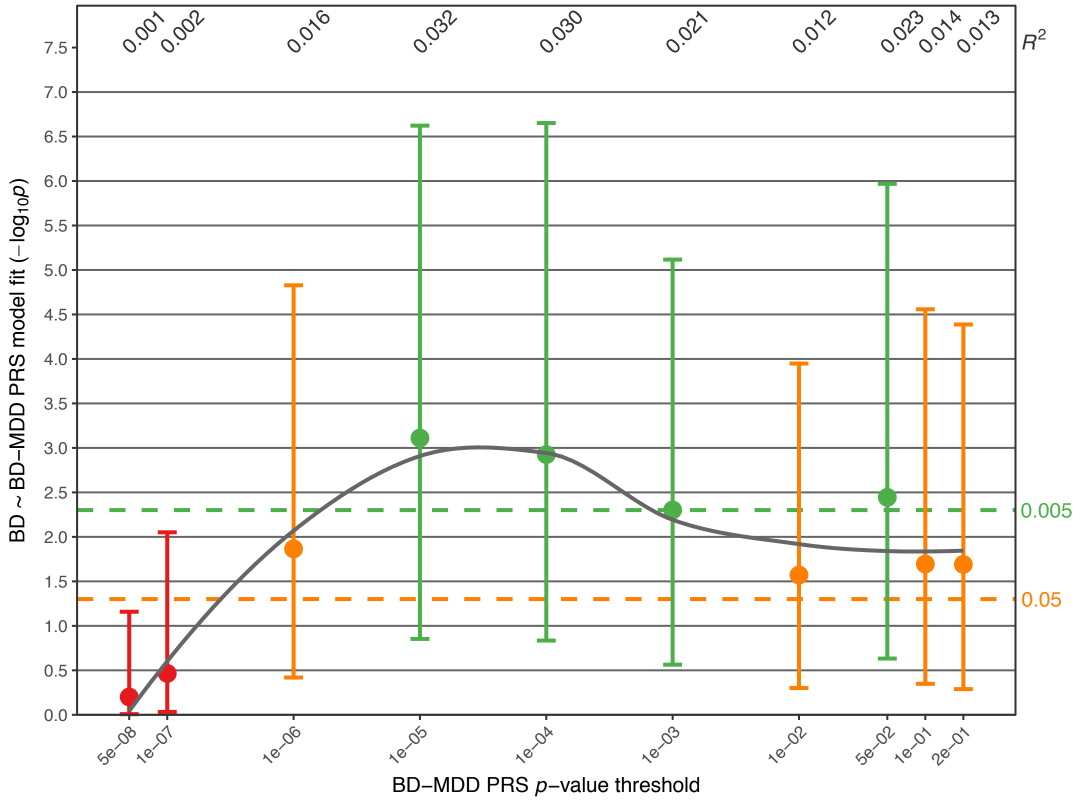

**Supplementary Fig. S5G:** Association of the SCZ-BD GWIS PRS.

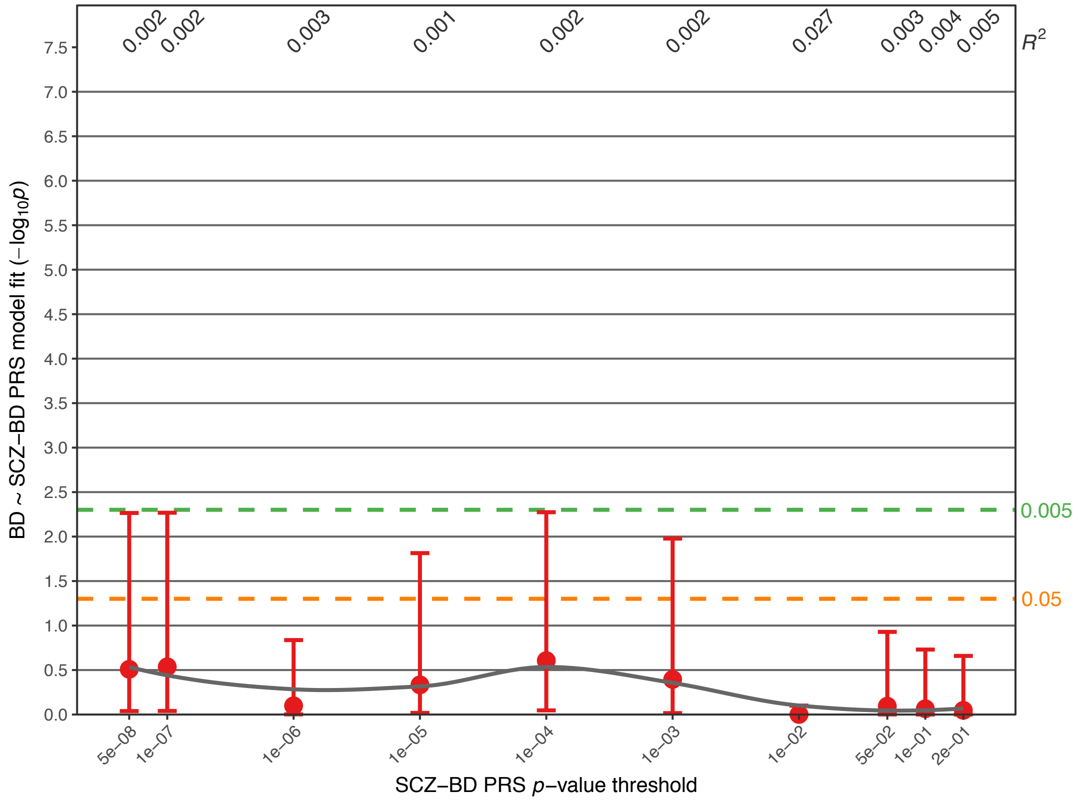


**Supplementary Fig. S5H:** Association of the MDD-BD GWIS PRS.

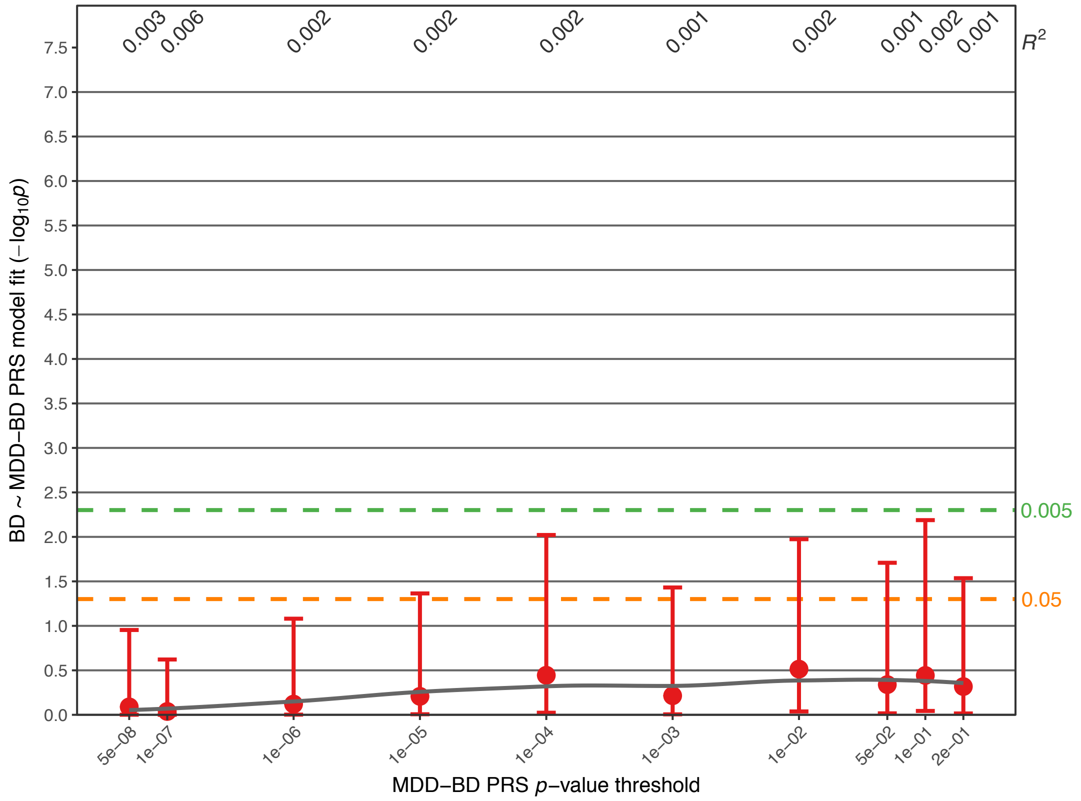

**Supplementary Fig. S5I:** Association of the LOAD PRS.

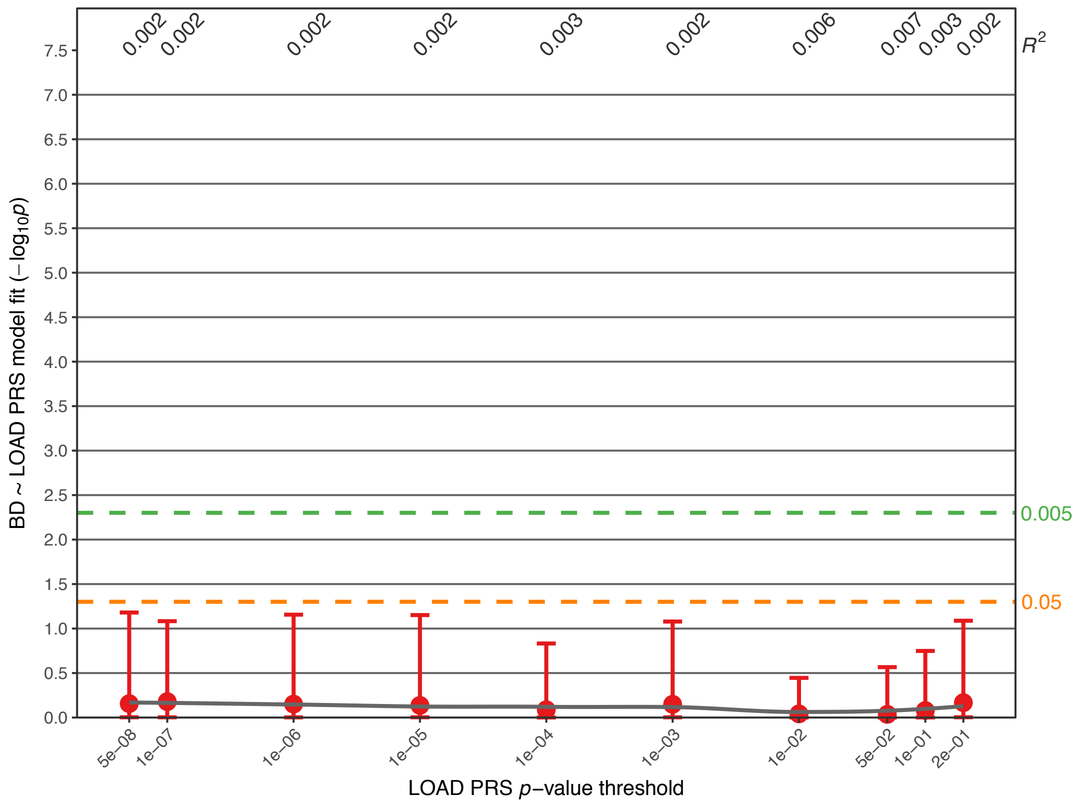

**Supplementary Fig. S6:** Analysis of assortative mating. Further details regarding the analysis and plots are described in the legend for Fig. 2C. Full association test statistics are shown in Supplementary Table S6.

**Supplementary Fig. S6A:** Association analysis comparing the BD PRS in unaffected married-in family members and $CC_{controls}$.

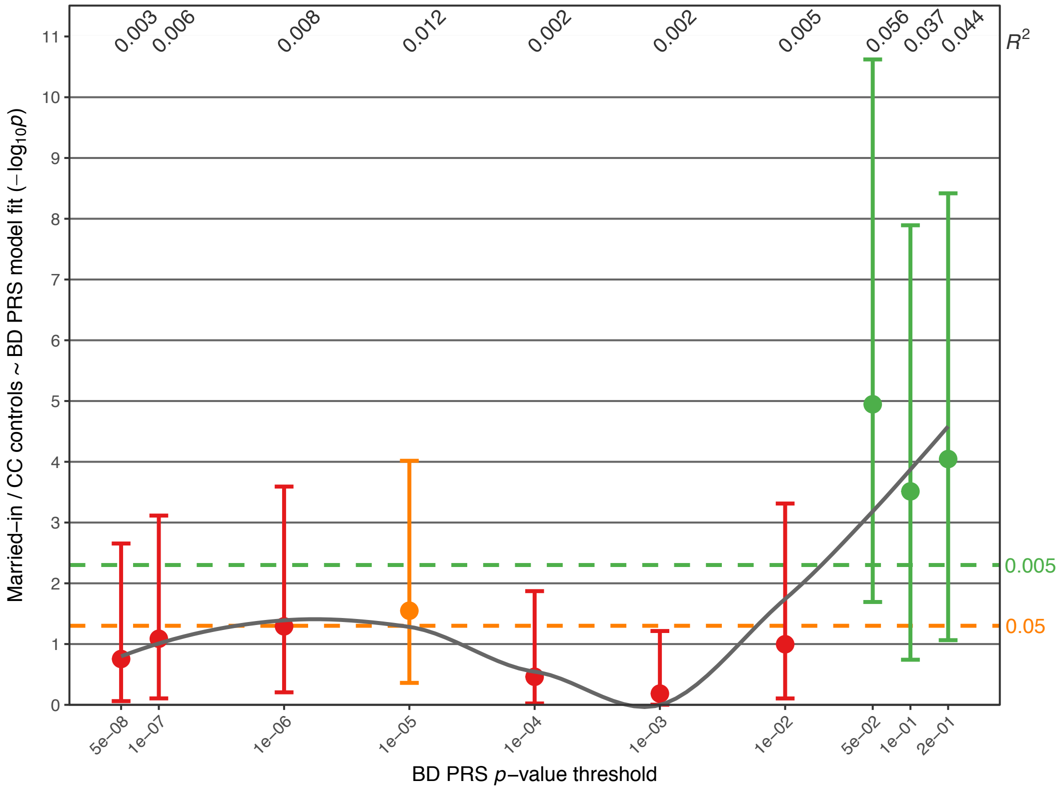


**Supplementary Fig. S6B:** Association analysis comparing the BD PRS in unaffected married-in family members to $FAM_{unaffected}$.

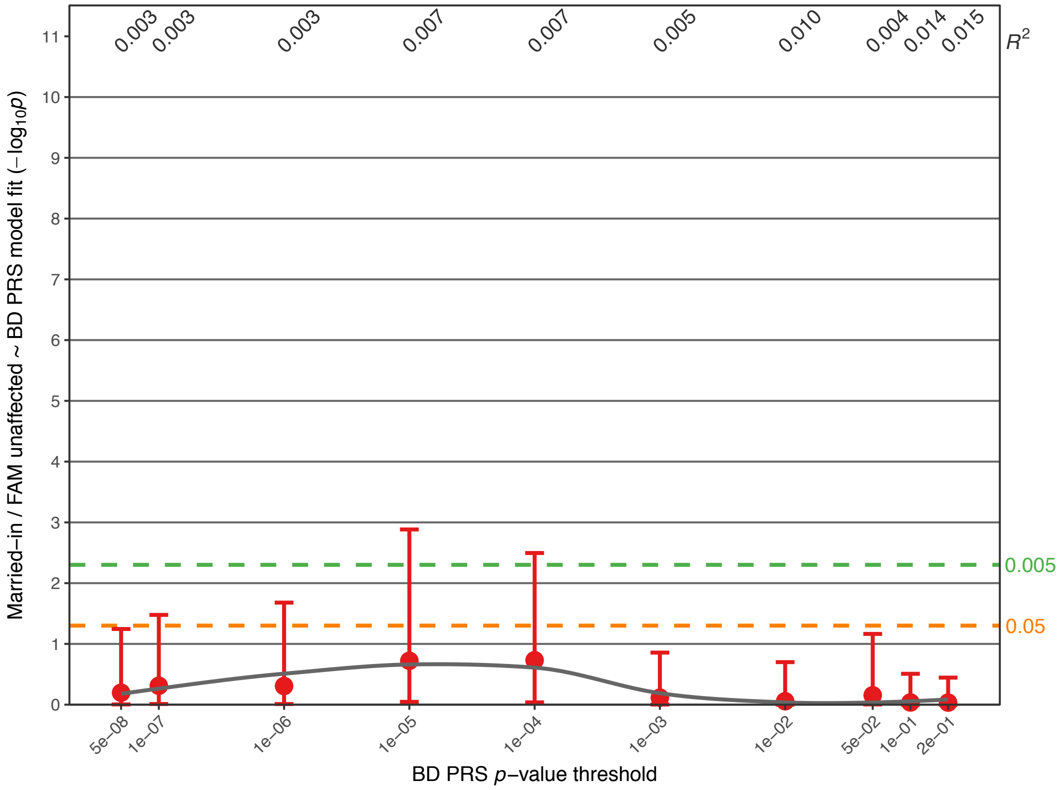

**Supplementary Fig. S7:** Analysis of anticipation in the FAM sample. Further details regarding the analysis and plots are described in the legend for Fig. 2D. Full association test statistics are shown in Supplementary Table S7.

**Supplementary Fig. S7A:** Association of the BD PRS with generation.

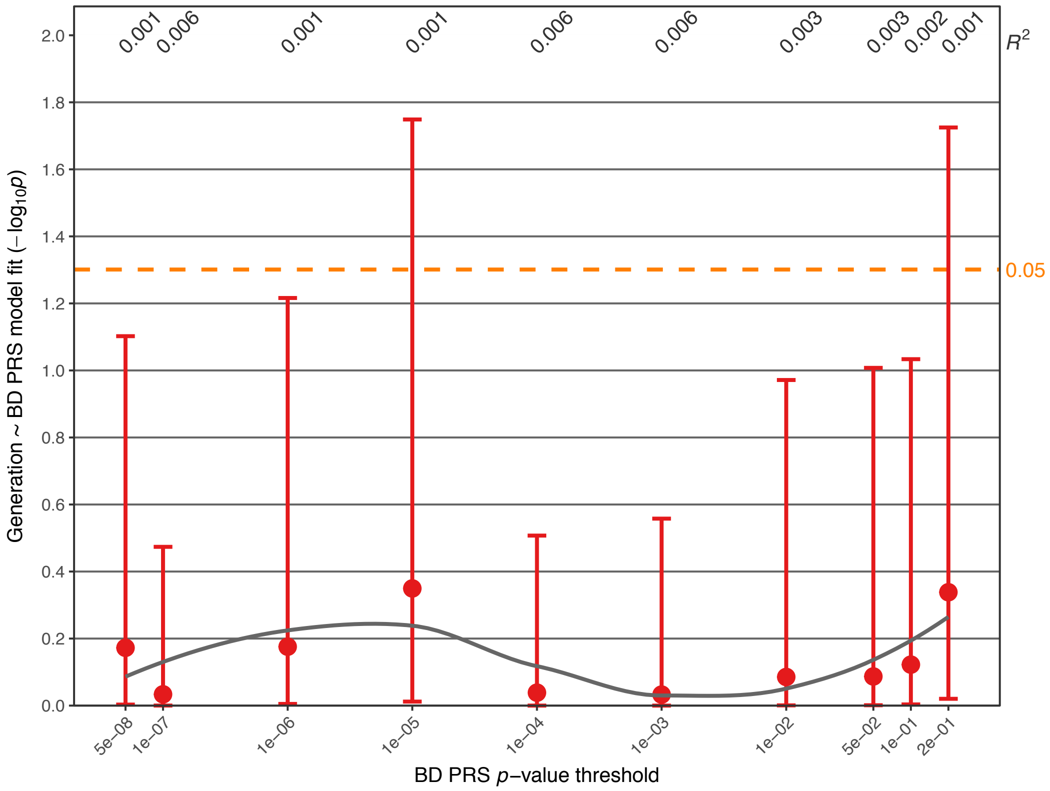


**Supplementary Fig. S7B:** Association analysis comparing the age at onset for BD across generations. The age at onset did not decrease over generations ($p$=0.54, Supplementary Table S7). Covariates were sex and age. One-sided $p$-values were calculated, following the hypothesis that the age at onset decreases across generations. The y-axis shows the residuals from the linear model.

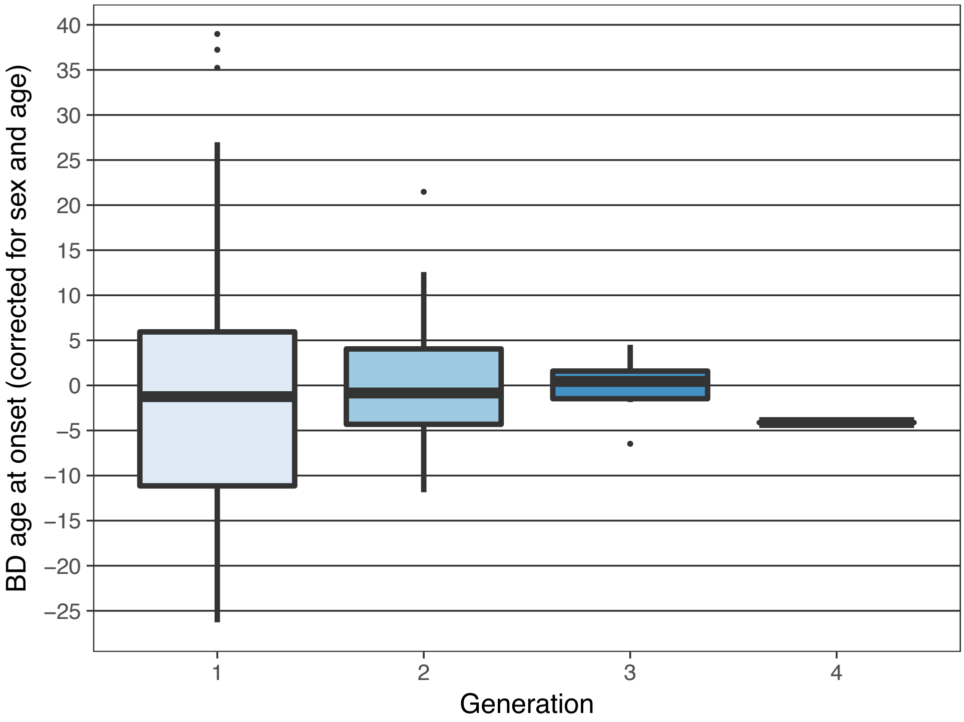

**Supplementary Fig. S8:** Boxplots of PRS at different *p*-value thresholds, including $FAM_{MDD}$ cases. The following indiduals are not shown in these plots: Family members with a history of substance abuse, married-in family members, and $CC_{BD}$ cases.

**Supplementary Fig. S8A:** Boxplots of BD PRS.

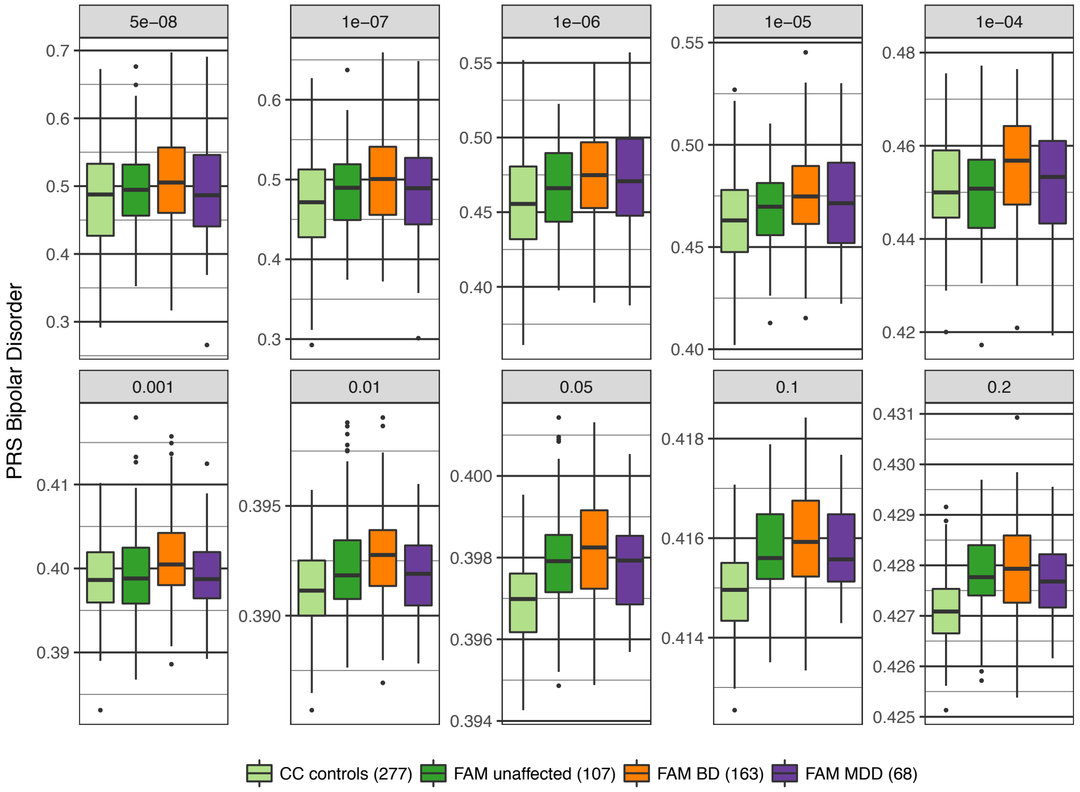


**Supplementary Fig. S8B:** Boxplots of SCZ PRS.

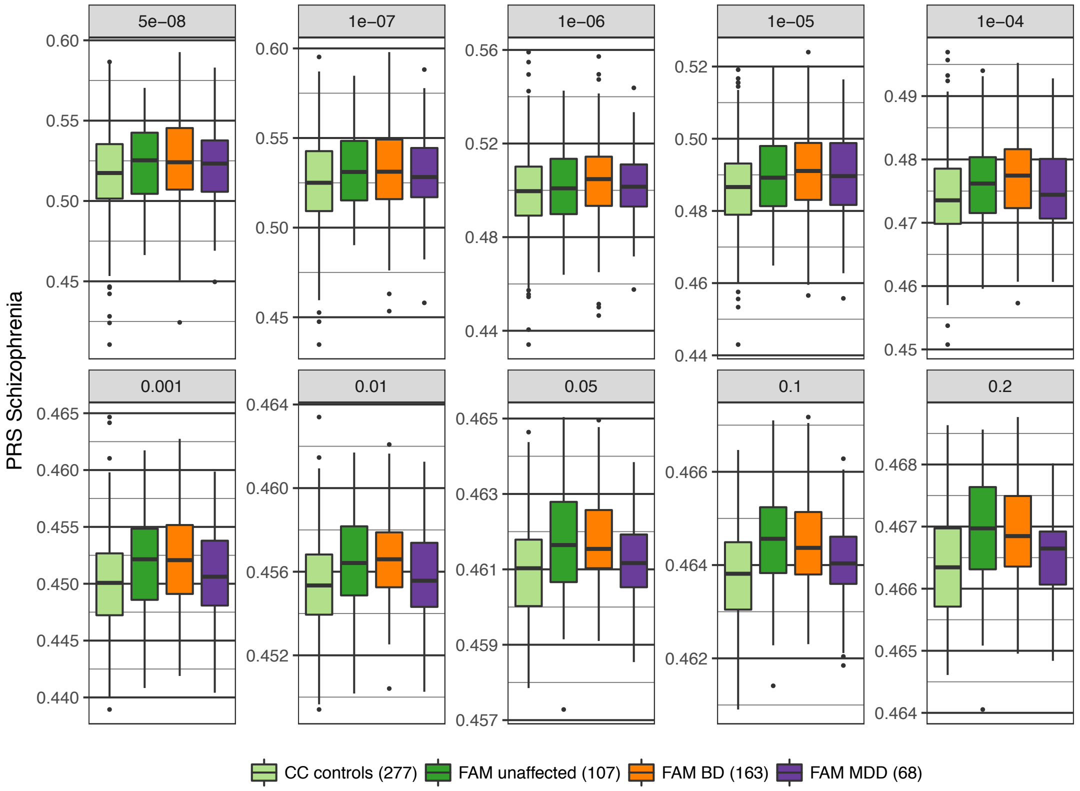

**Supplementary Fig. S8C:** Boxplots of MDD PRS.

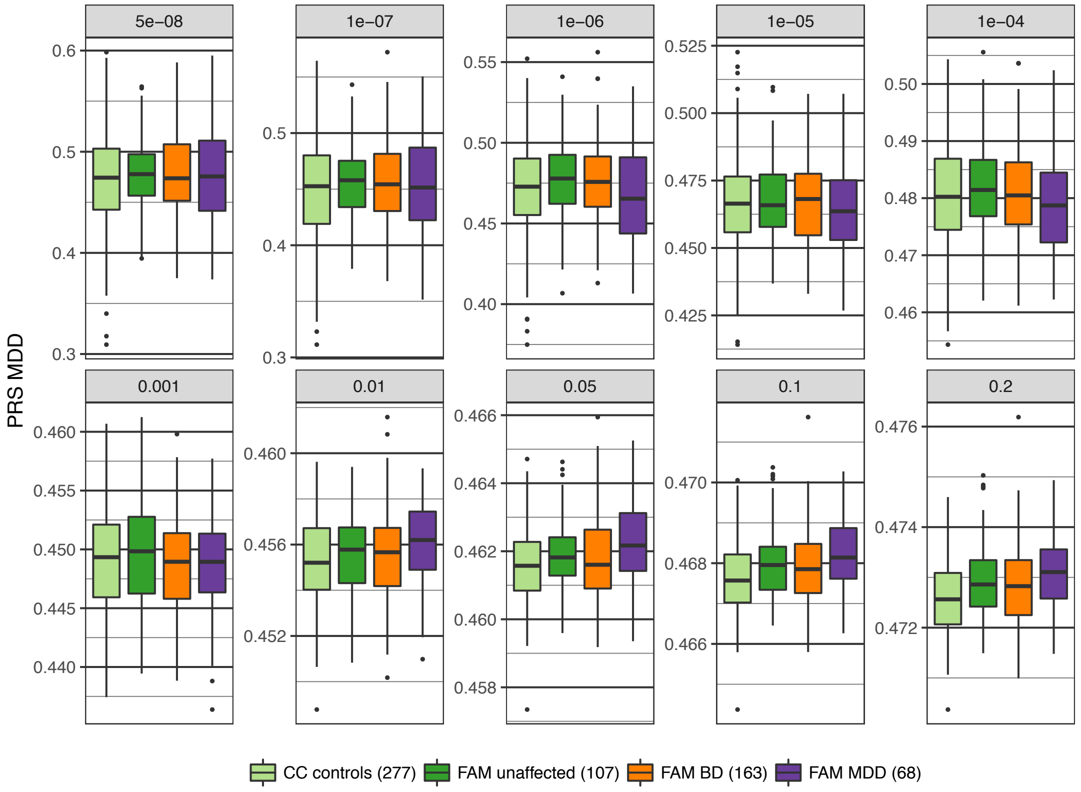


**Supplementary Fig. S8D:** Boxplots of the BD+SCZ+MDD *Shared* PRS.

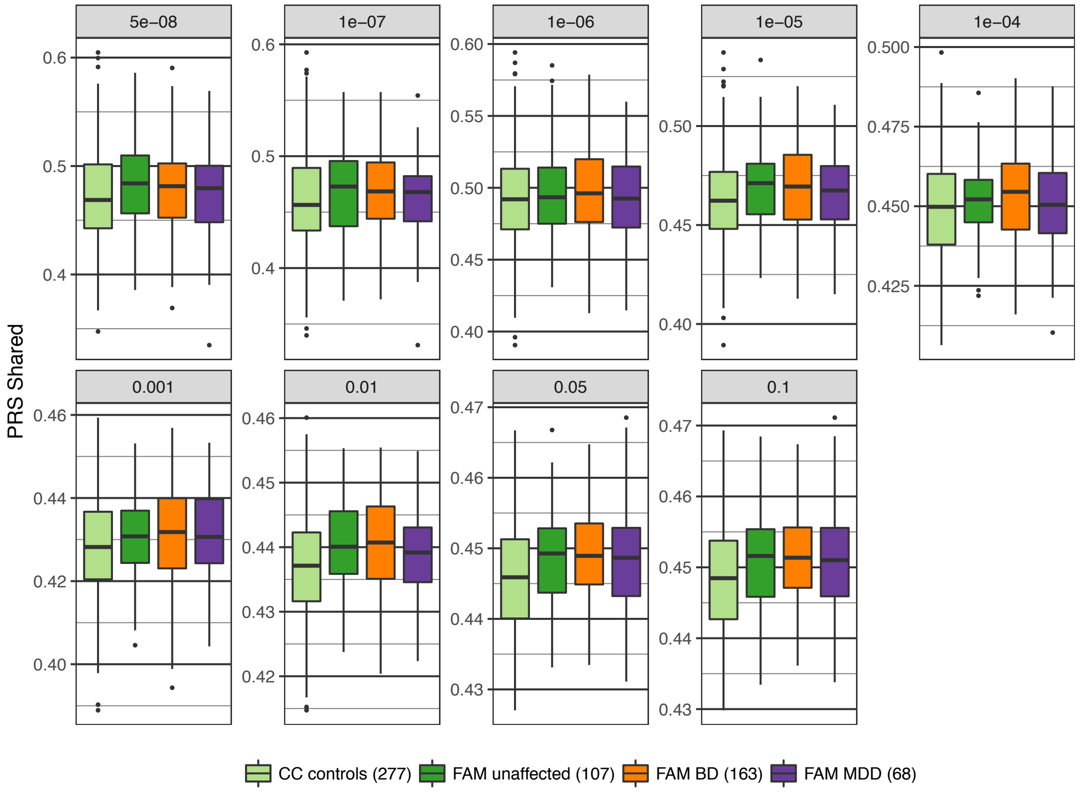

**Supplementary Fig. S8E:** Boxplots of the MDD-BD GWIS PRS.

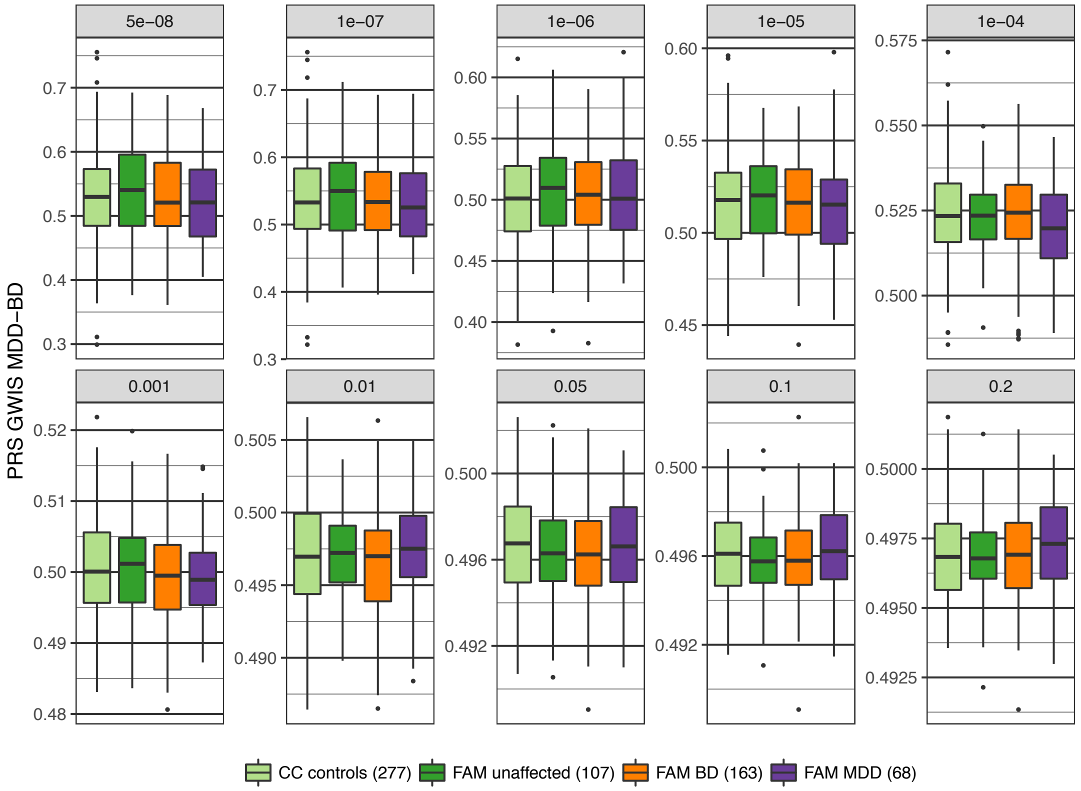


**Supplementary Fig. S8F:** Boxplots of the MDD-SCZ GWIS PRS.

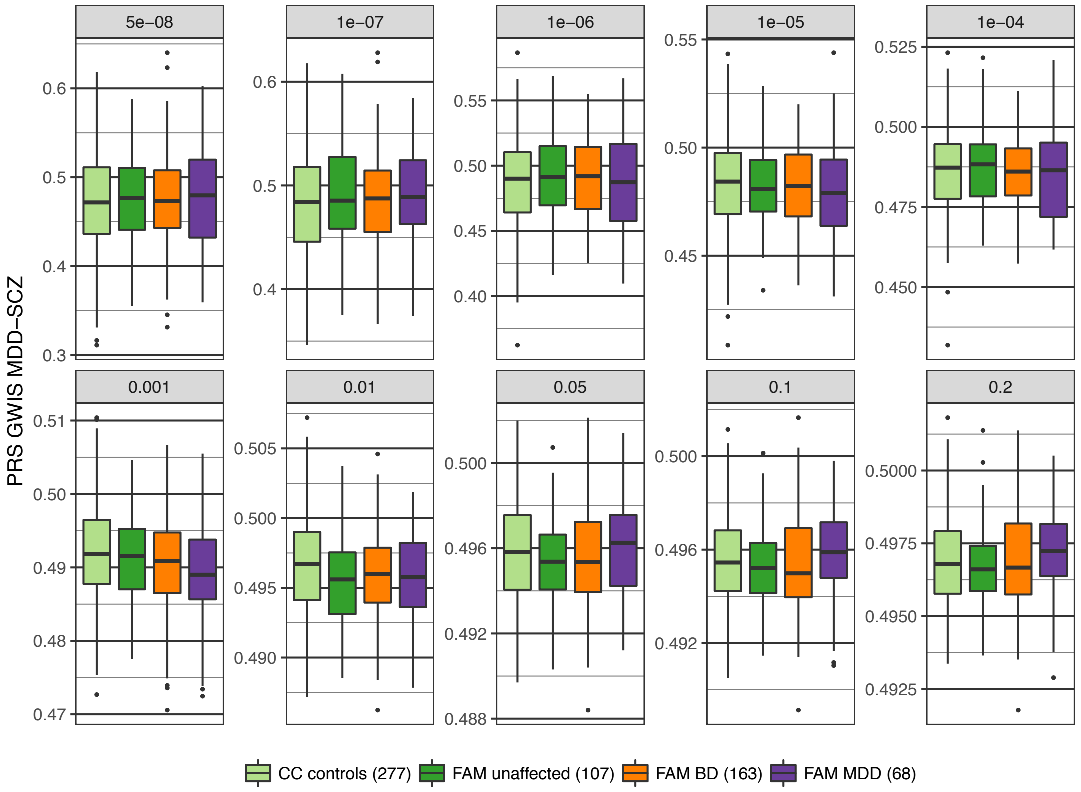

**Supplementary Fig. S8G:** Boxplots of the BD-MDD GWIS PRS.

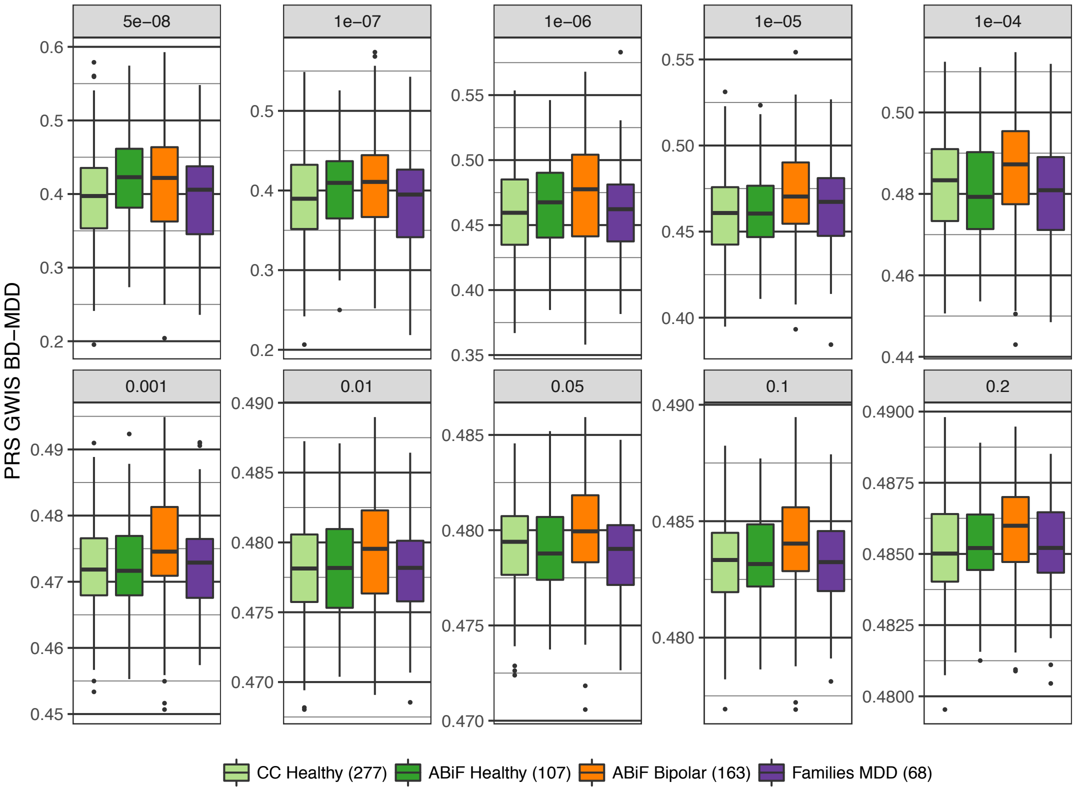


**Supplementary Fig. S8H:** Boxplots of the SCZ-MDD GWIS PRS.

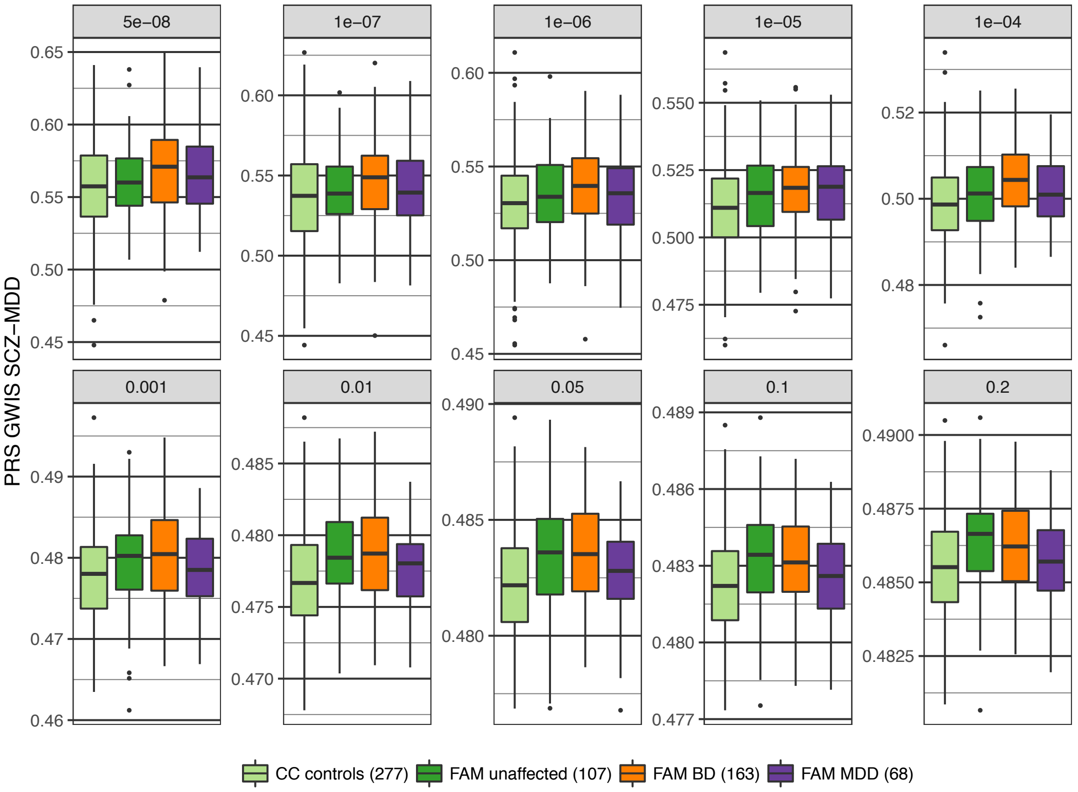

**Supplementary Fig. S8I:** Boxplots of the SCZ-BD GWIS PRS.

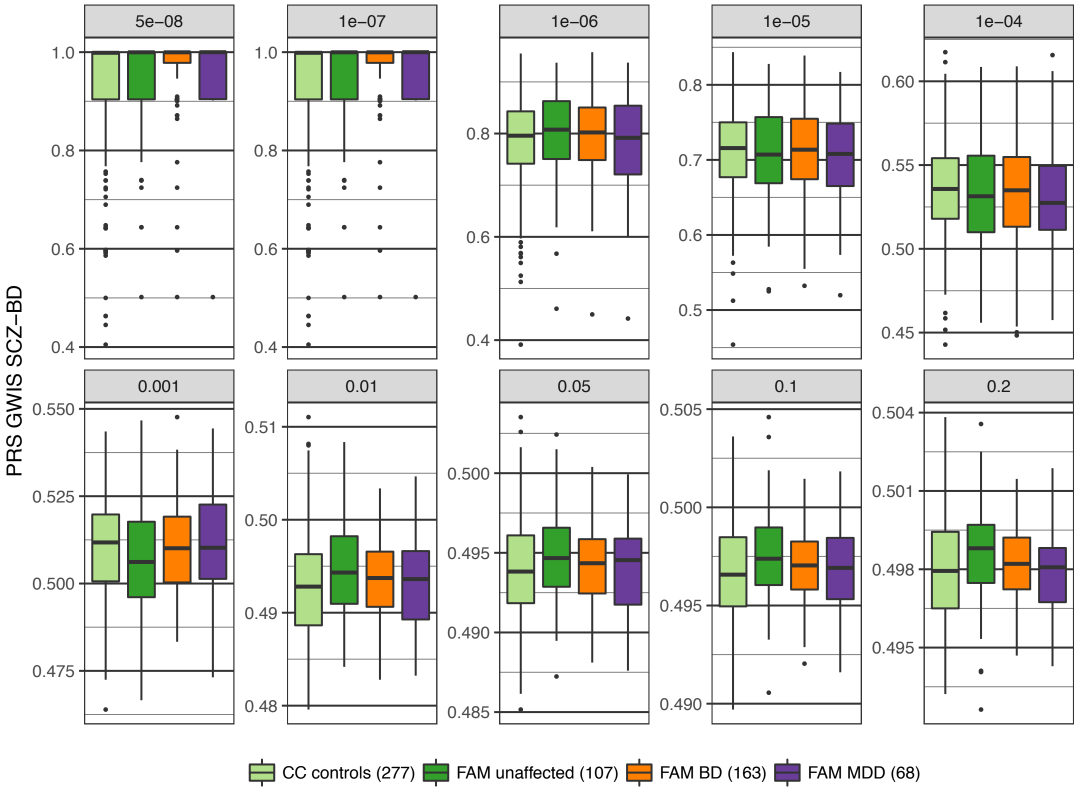


**Supplementary Fig. S8J:** Boxplots of the LOAD (Alzheimer) PRS.

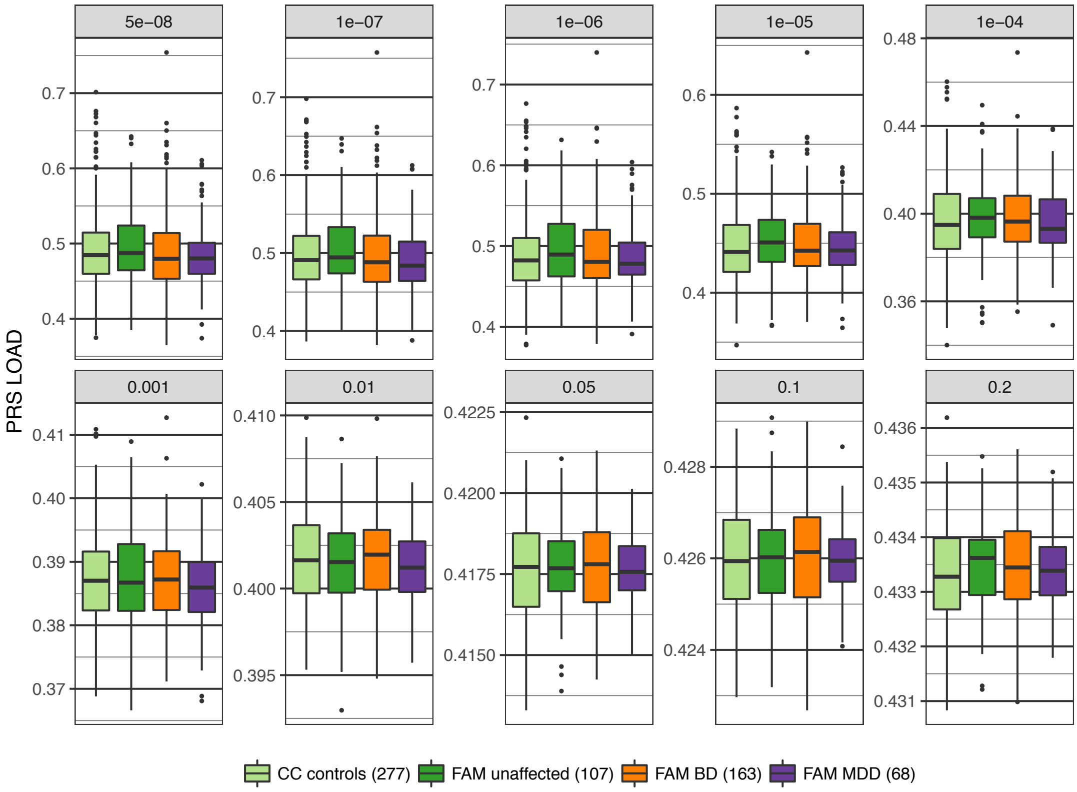

**Supplementary Fig. S9:** Association analysis comparing PRS in $FAM_{MDD}$ cases and $CC_{controls}$. Further details of the plots are described in the legend for Fig. 1. Full association test statistics are shown in Supplementary Table S8.

**Supplementary Fig. S9A:** Top-associated *p*-value thresholds for the tested PRS. The column to the left shows PRS and $p_{PRS}$ thresholds.

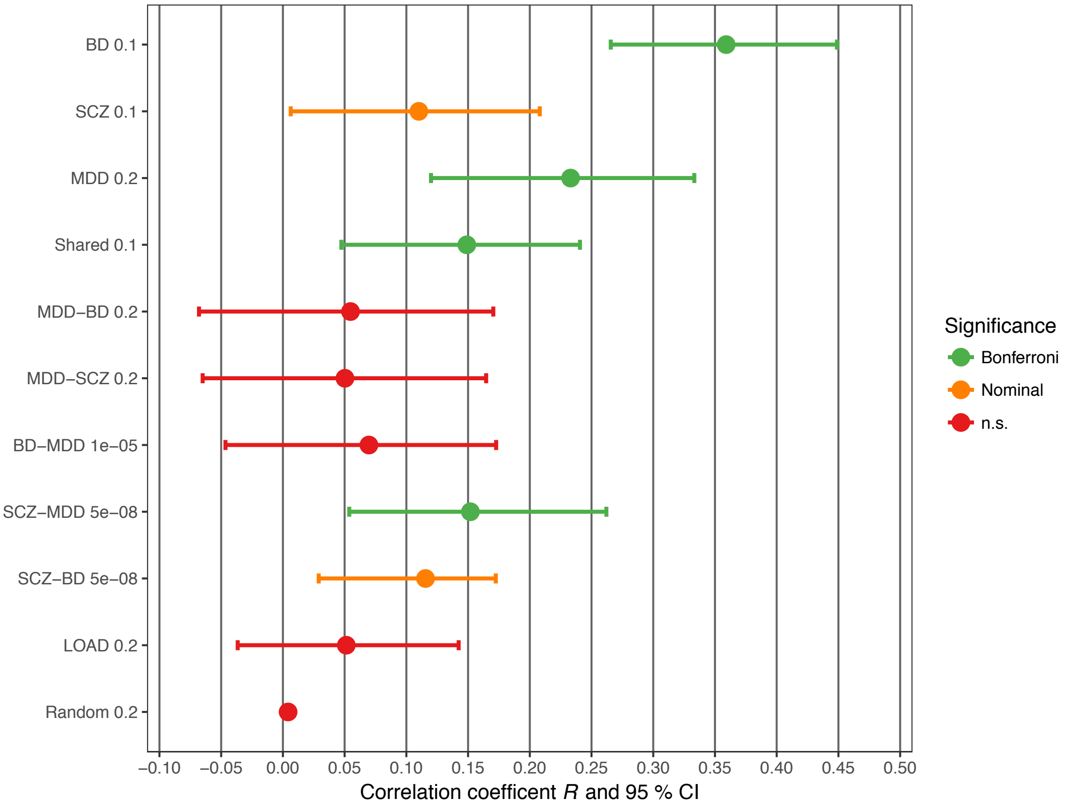


**Supplementary Fig. S9B:** Association of the BD PRS (all *p*-value thresholds).

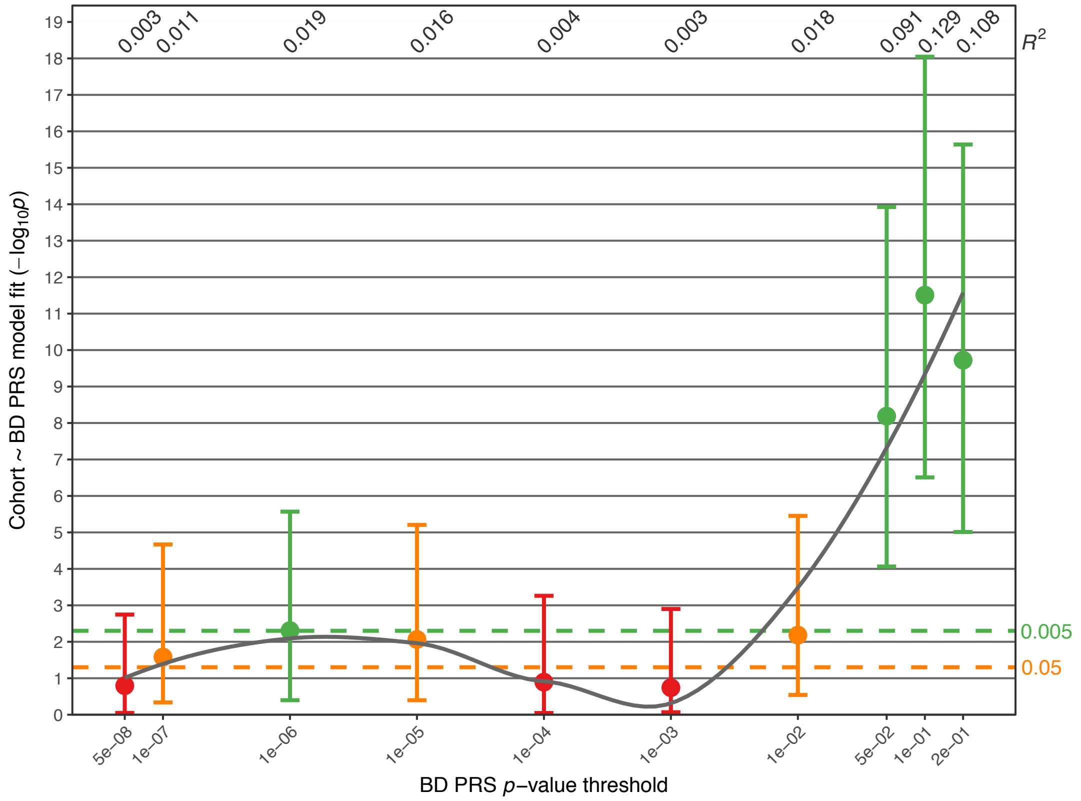

**Supplementary Fig. S9C:** Association of the SCZ PRS.

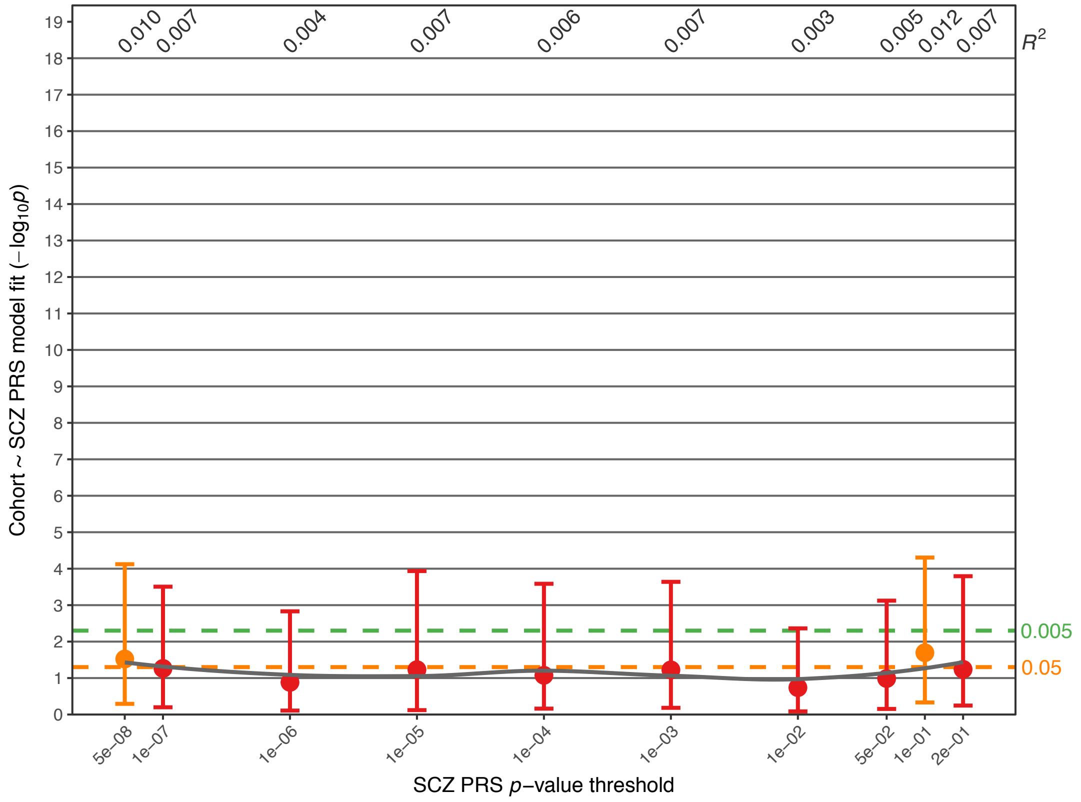


**Supplementary Fig. S9D:** Association of the MDD PRS.

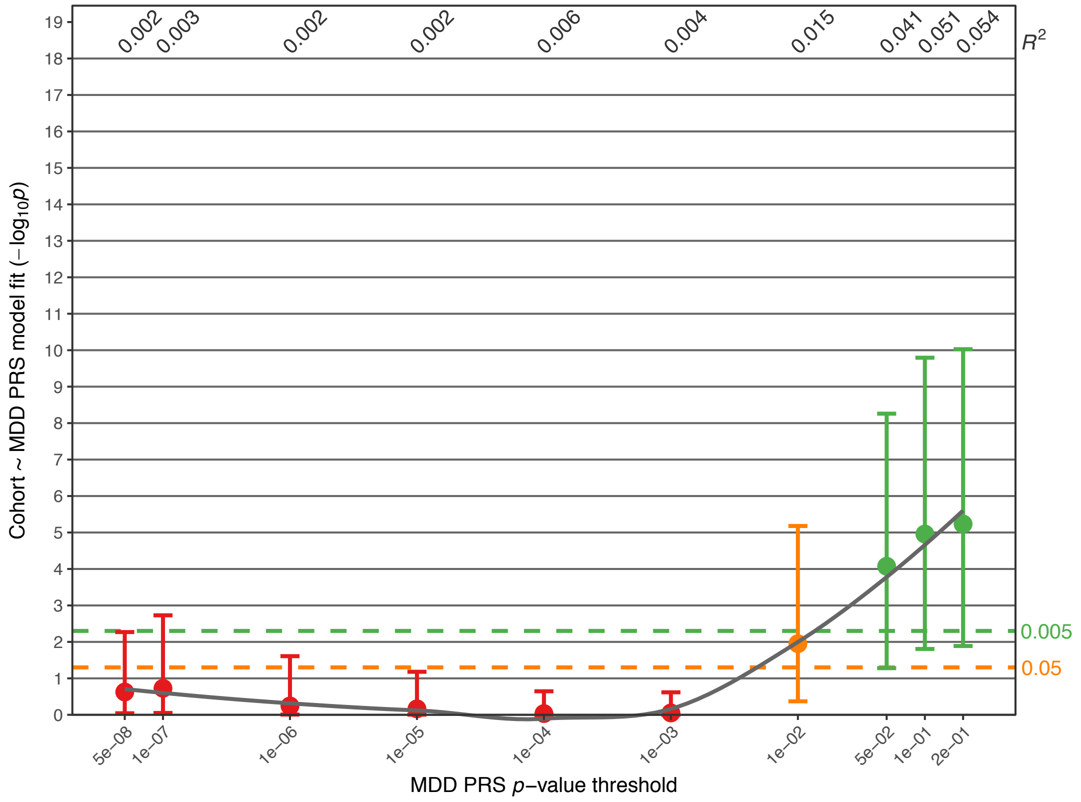

**Supplementary Fig. S9E:** Association of the *Shared* PRS.

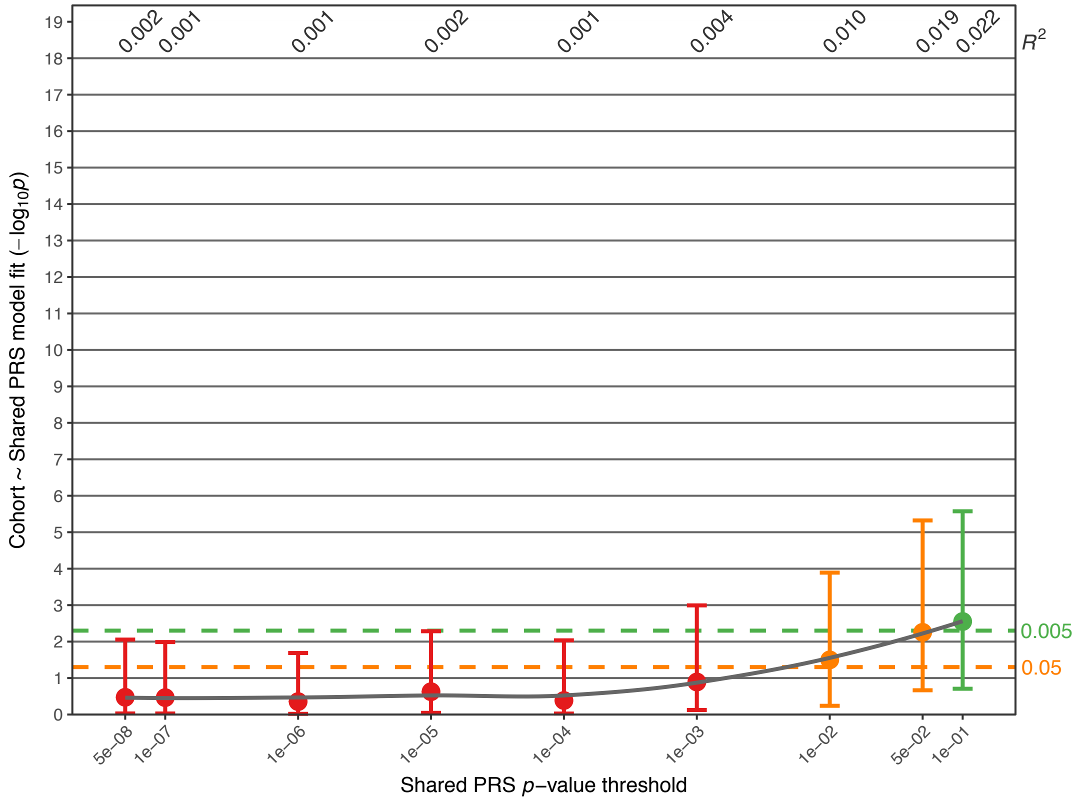


**Supplementary Fig. S9F:** Association of the MDD-BD GWIS PRS.

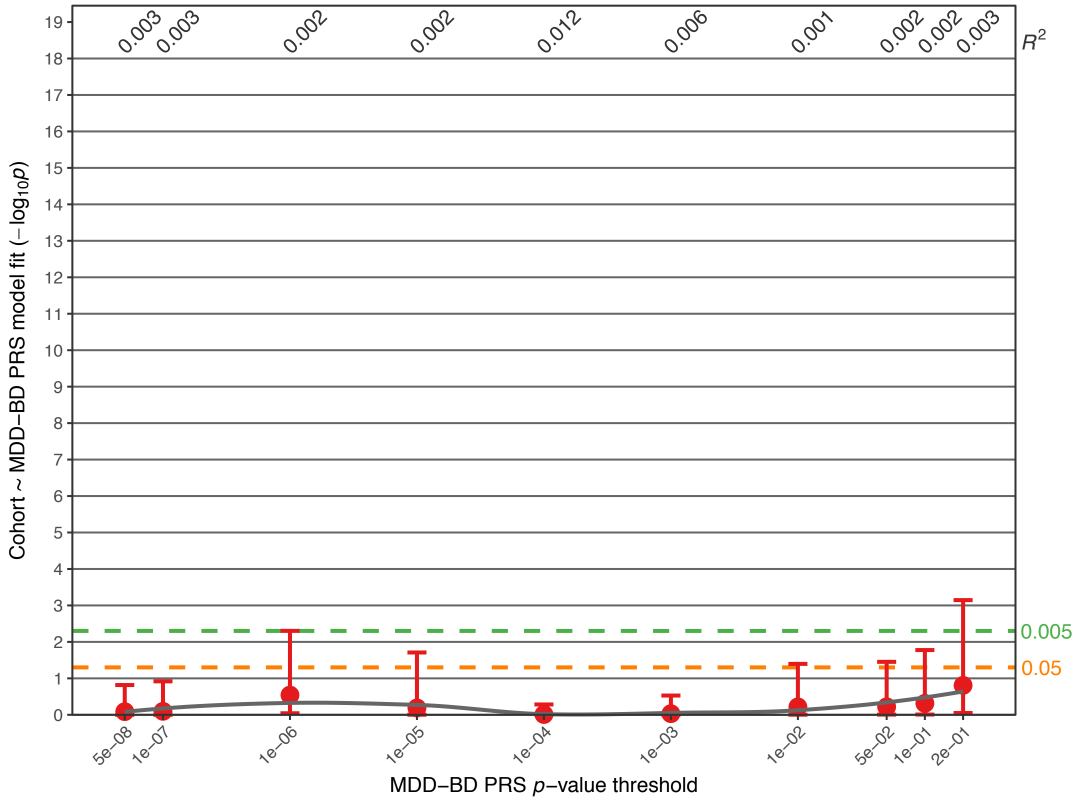

**Supplementary Fig. S9G:** Association of the MDD-SCZ GWIS PRS.

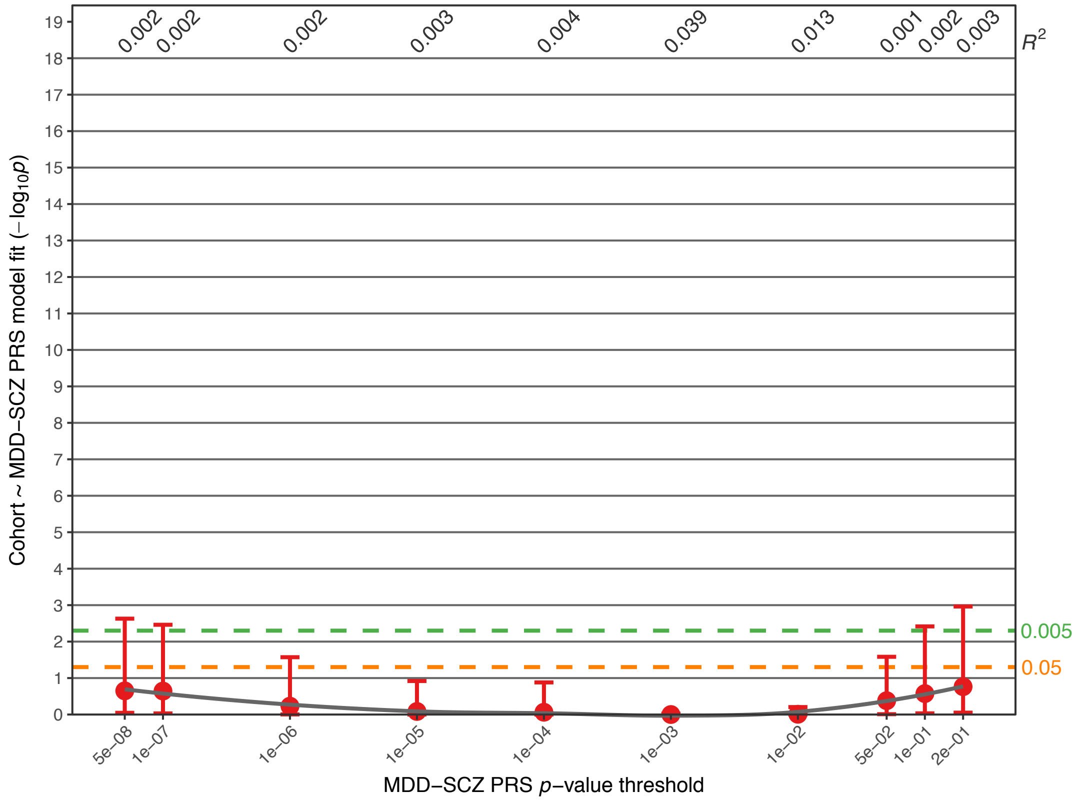


**Supplementary Fig. S9H:** Association of the BD-MDD GWIS PRS.

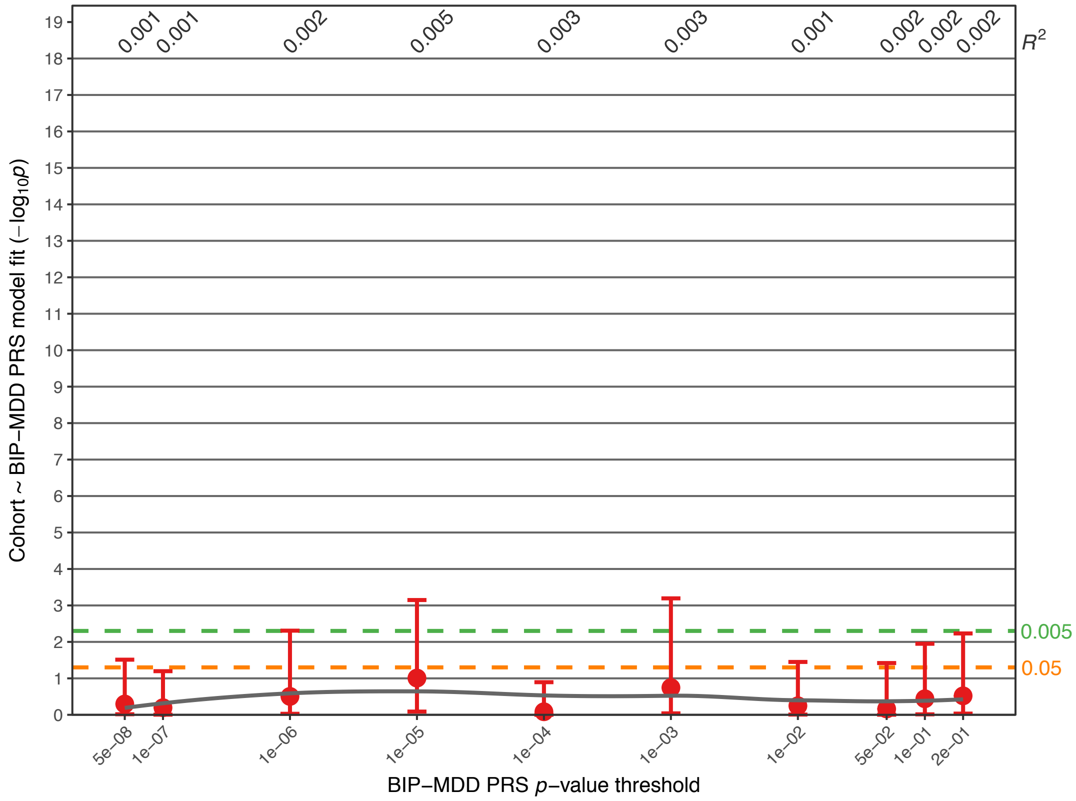

**Supplementary Fig. S9I:** Association of the SCZ-MDD GWIS PRS.

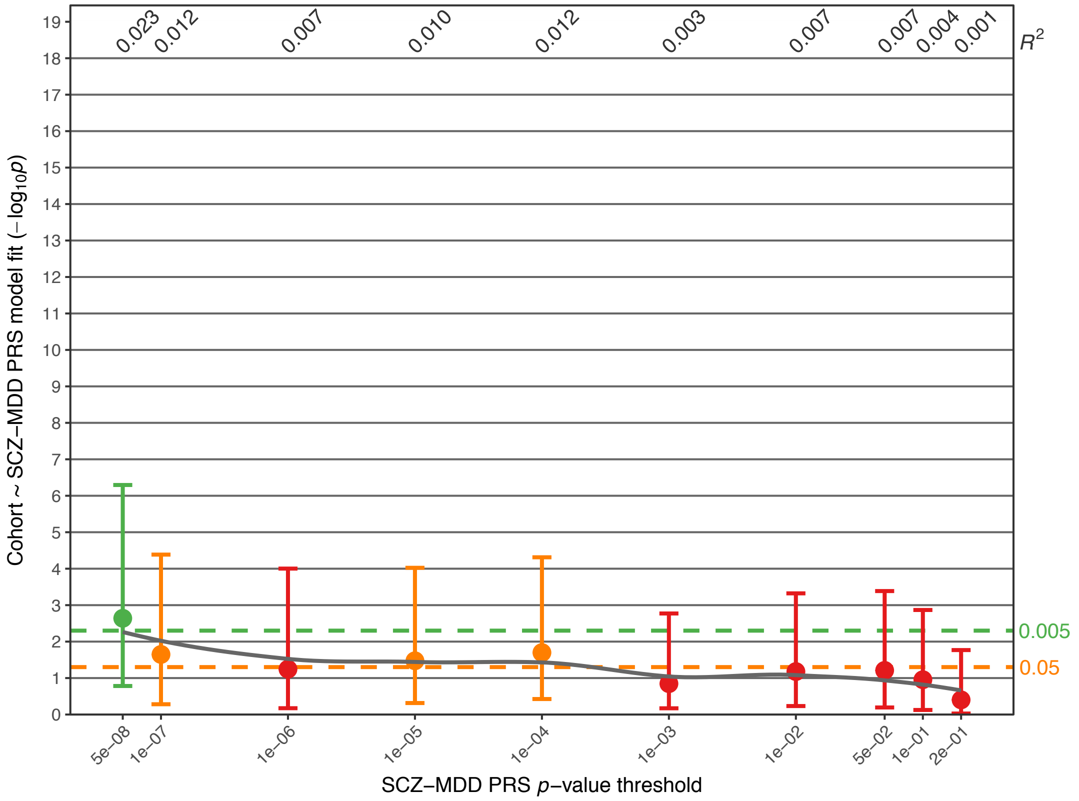


**Supplementary Fig. S9J:** Association of the SCZ-BD GWIS PRS.

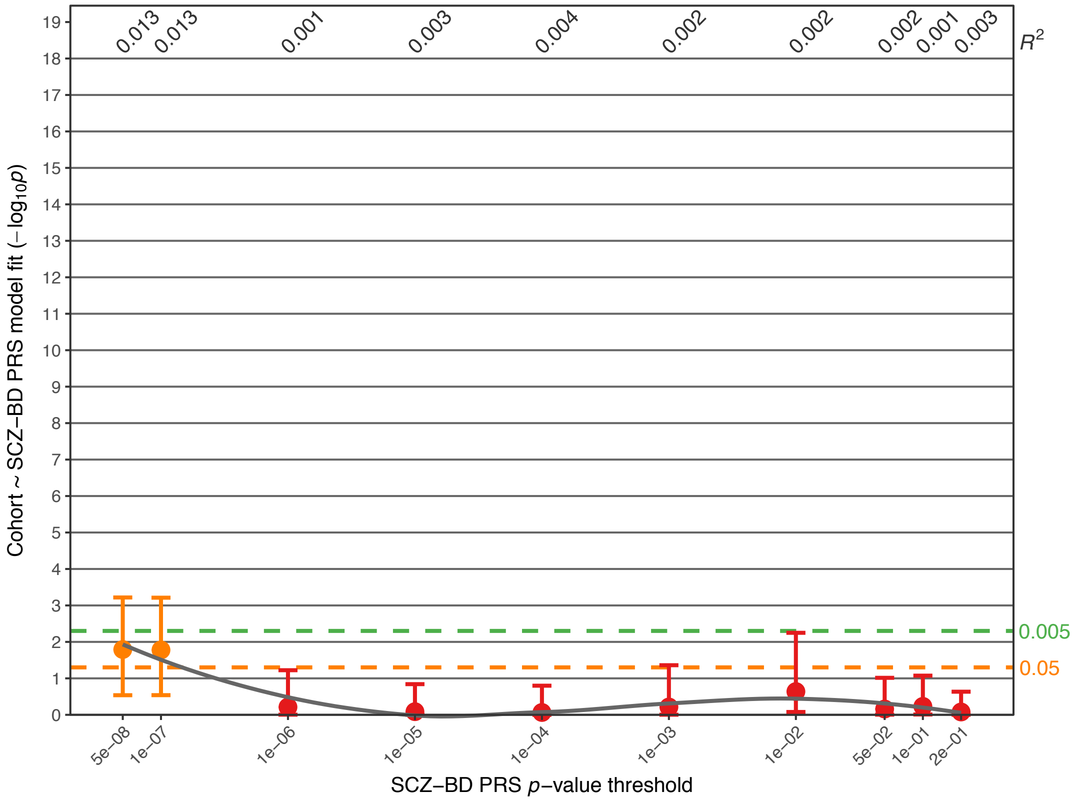

**Supplementary Fig. S9K:** Association of the LOAD PRS.

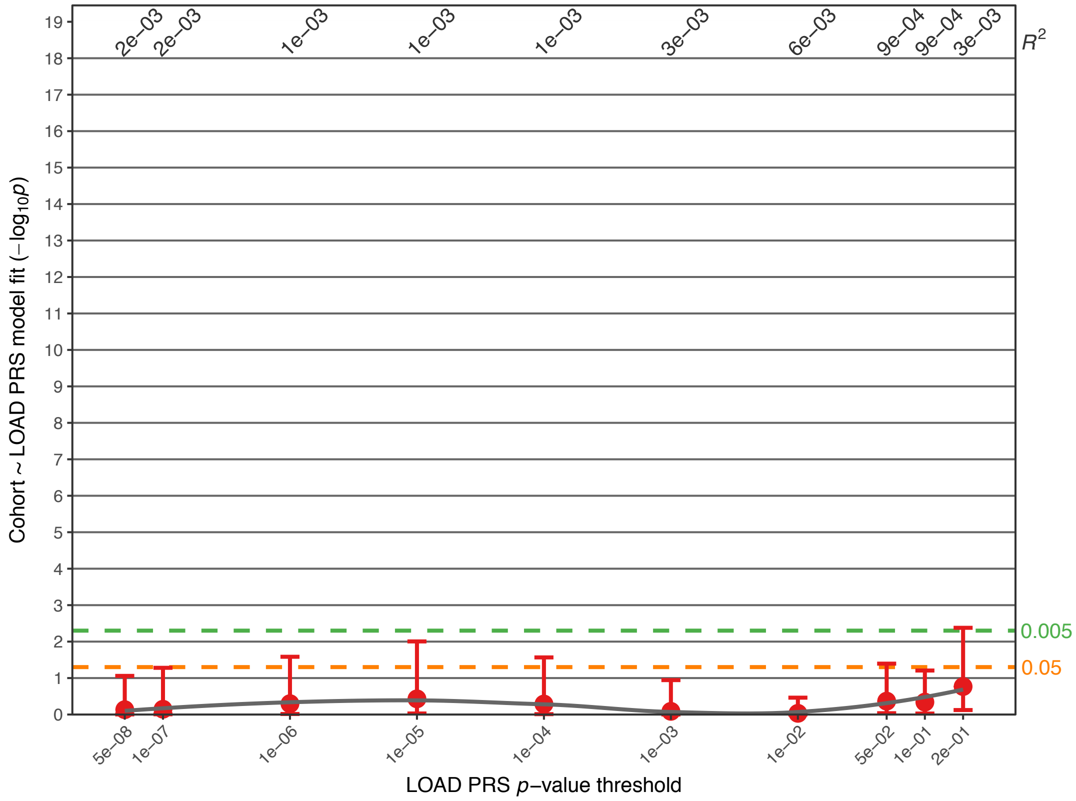


**Supplementary Fig. S10:** Association analysis comparing PRS in $FAM_{MDD}$ cases and $FAM_{unaffected}$. Further details of the plots are described in the legends for Figs. 1 and 2. Full association test statistics are shown in Supplementary Table S9.
**Supplementary Fig. S10A:** Top-associated $p_{PRS}$ thresholds for the tested PRS. The column to the left shows PRS and $p_{PRS}$ thresholds.

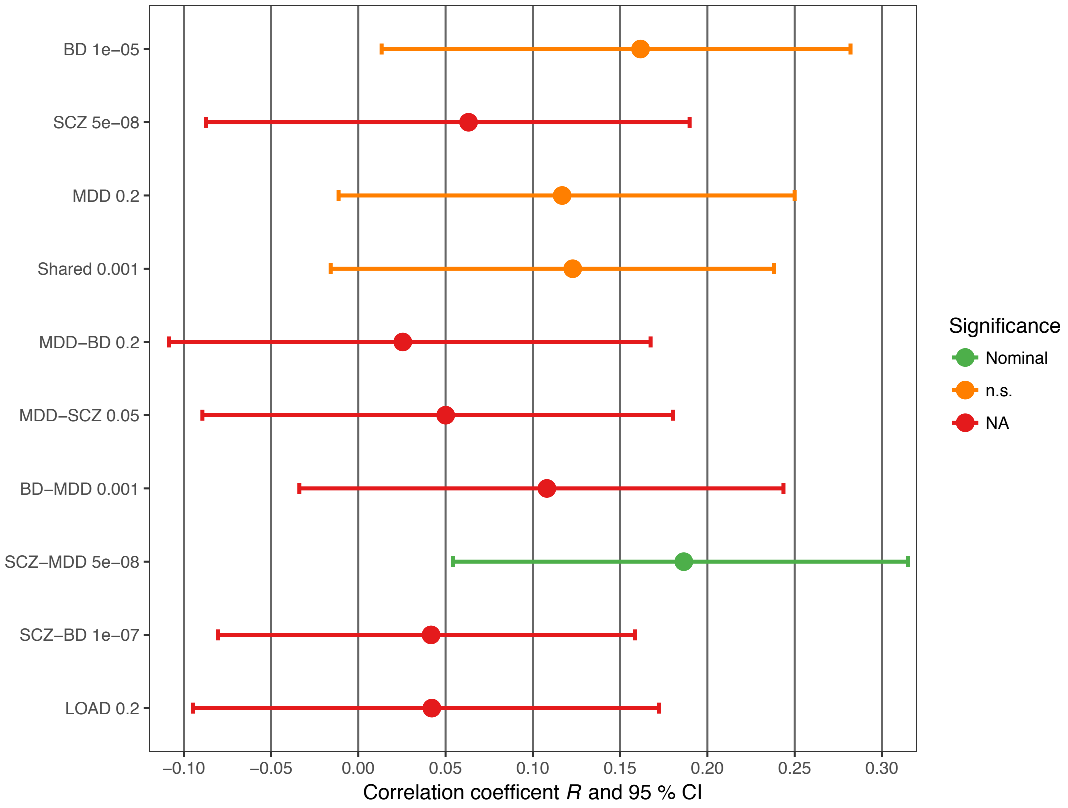

**Supplementary Fig. S10B:** Association of the BD PRS (all *p*-value thresholds).

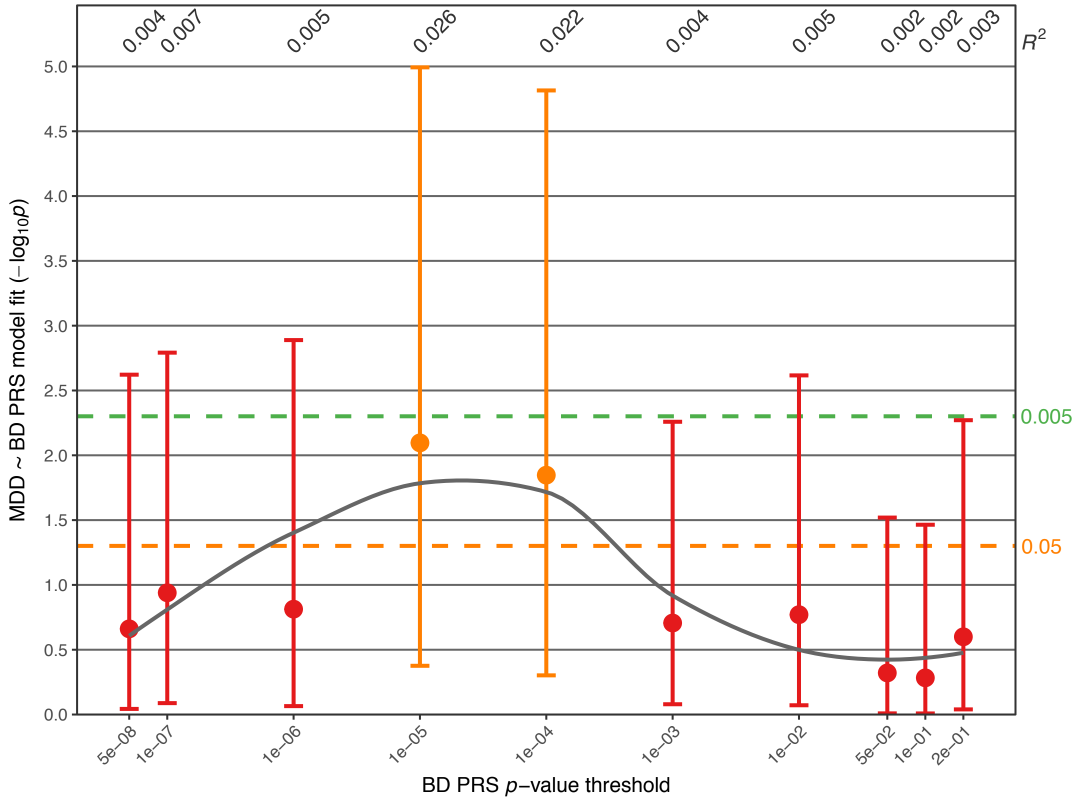


**Supplementary Fig. S10C:** Association of the SCZ PRS.

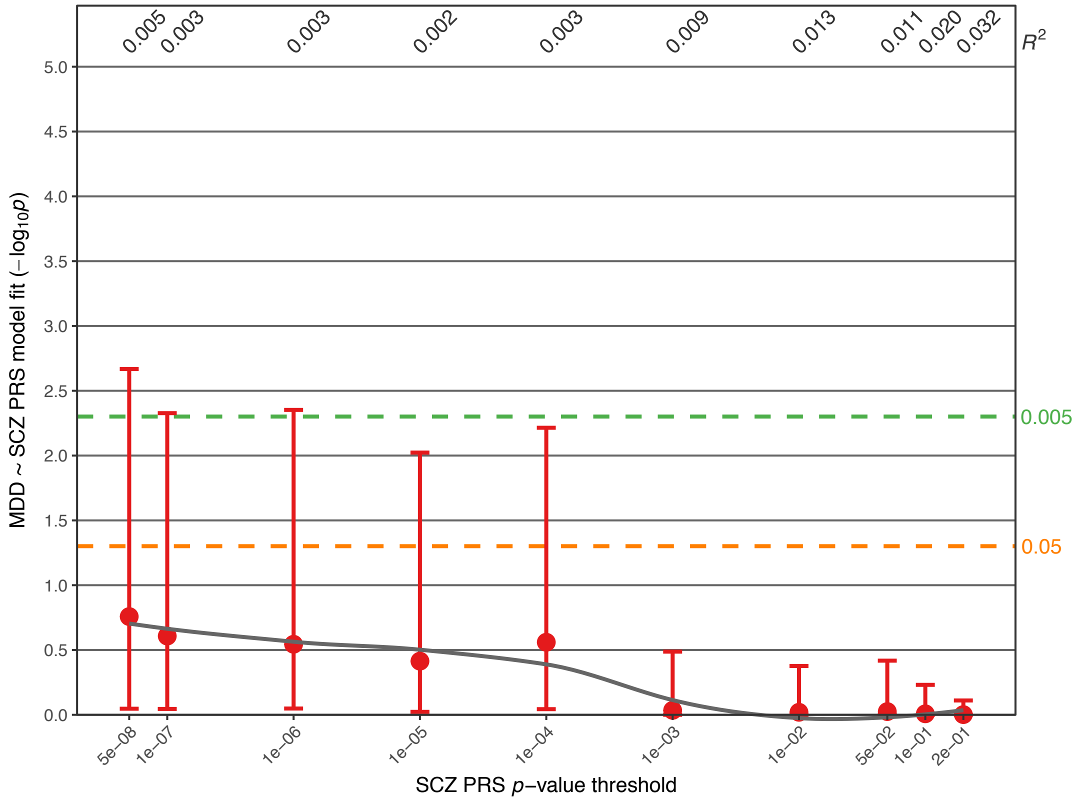

**Supplementary Fig. S10D:** Association of the MDD PRS.

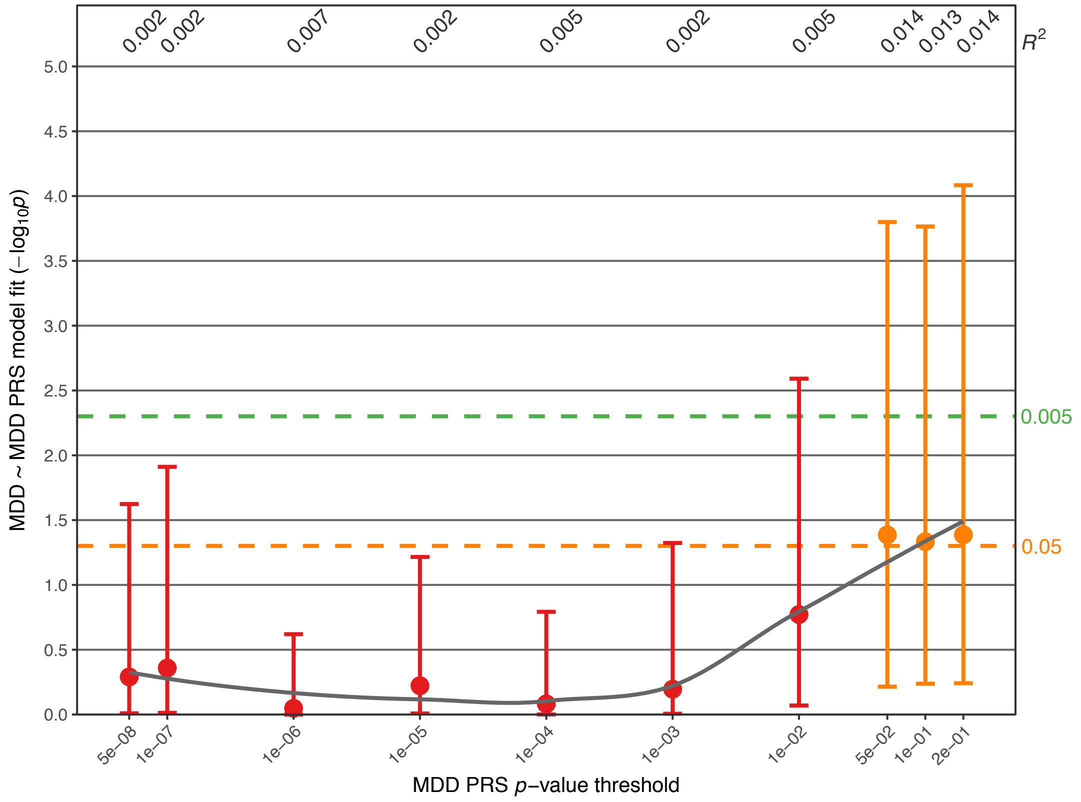


**Supplementary Fig. S10E:** Association of the *Shared* PRS.

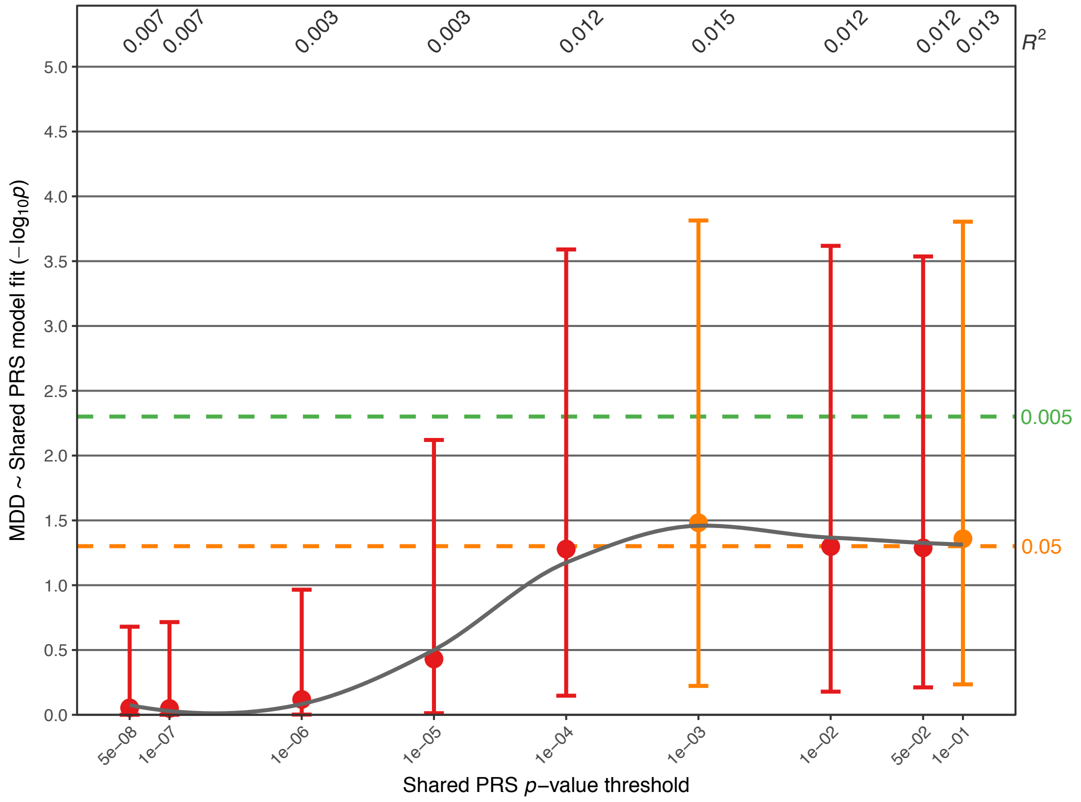

**Supplementary Fig. S10F:** Association of the MDD-BD GWIS PRS.

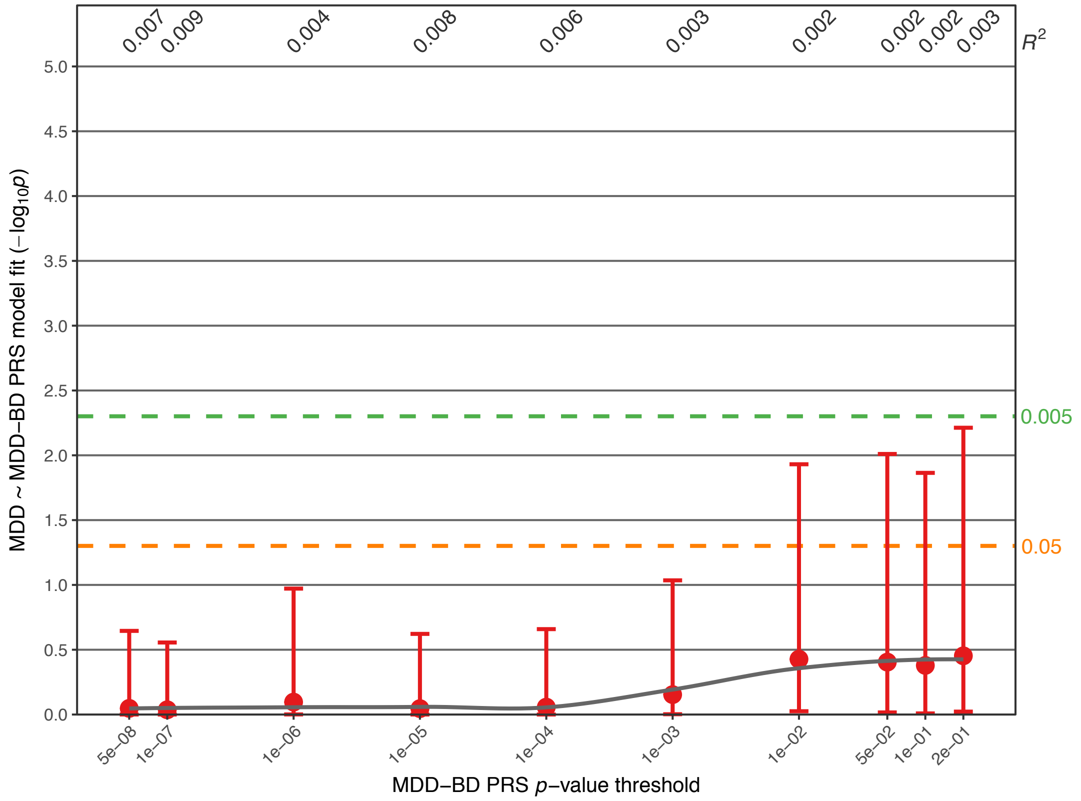


**Supplementary Fig. S10G:** Association of the MDD-SCZ GWIS PRS.

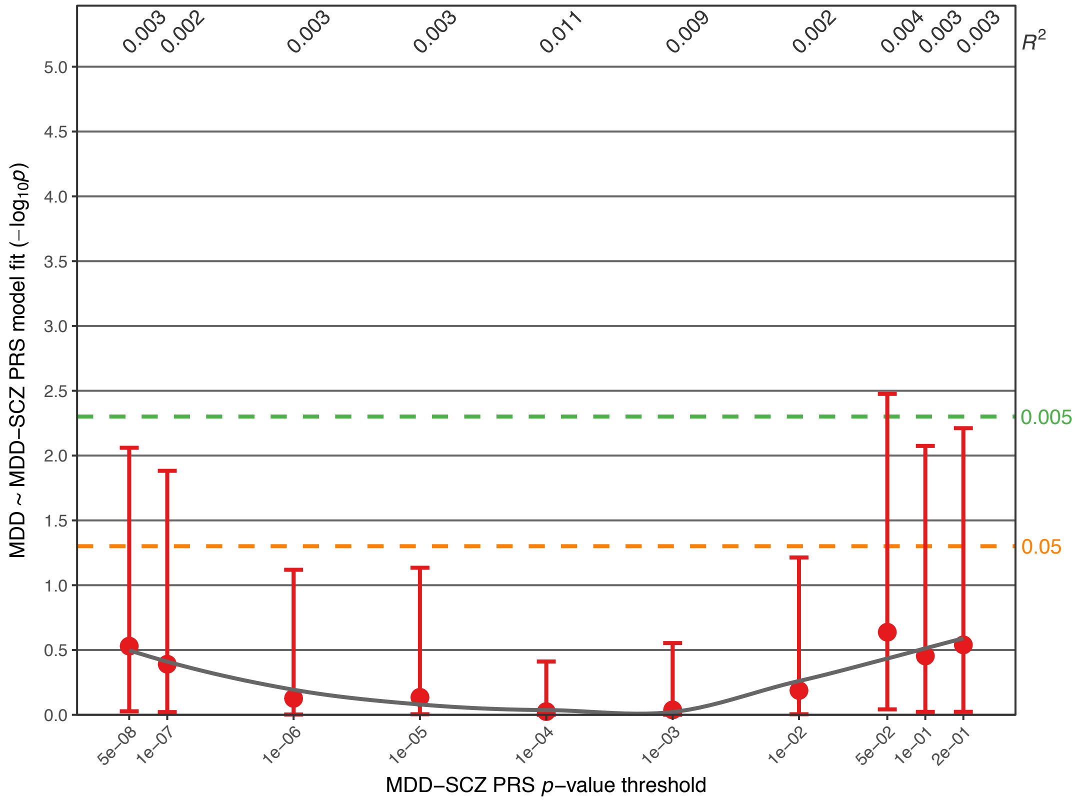

**Supplementary Fig. S10H:** Association of the BD-MDD GWIS PRS.

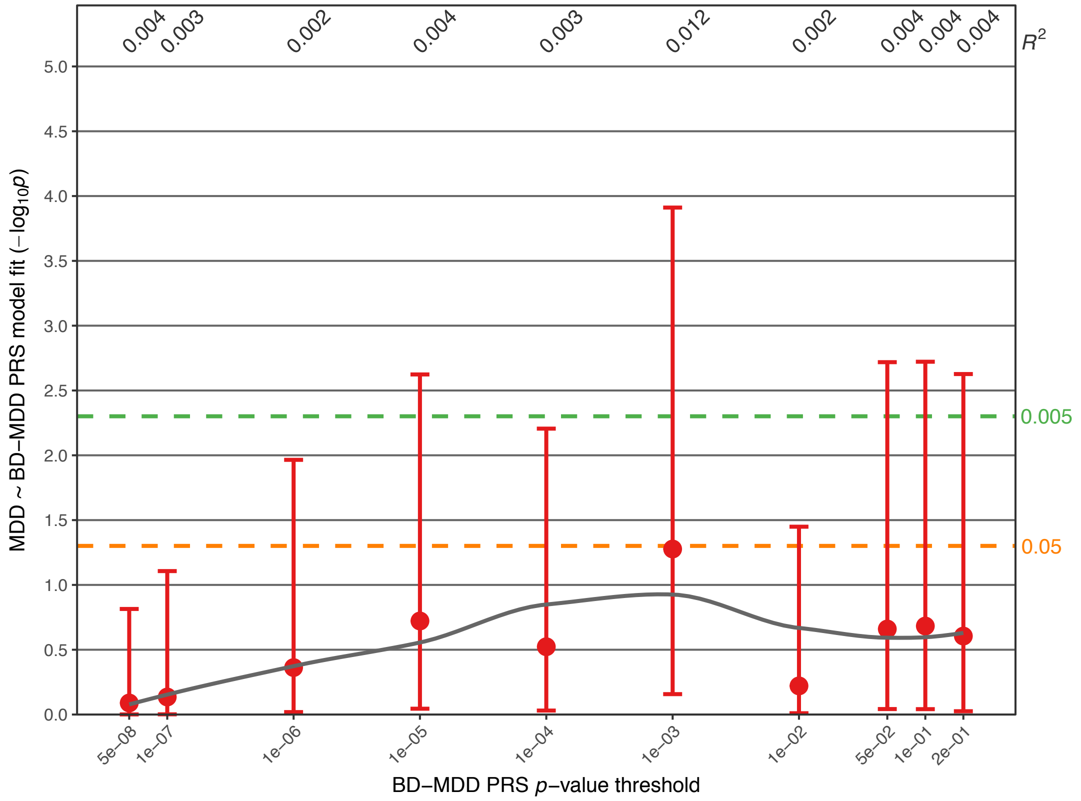


**Supplementary Fig. S10I:** Association of the SCZ-MDD GWIS PRS.

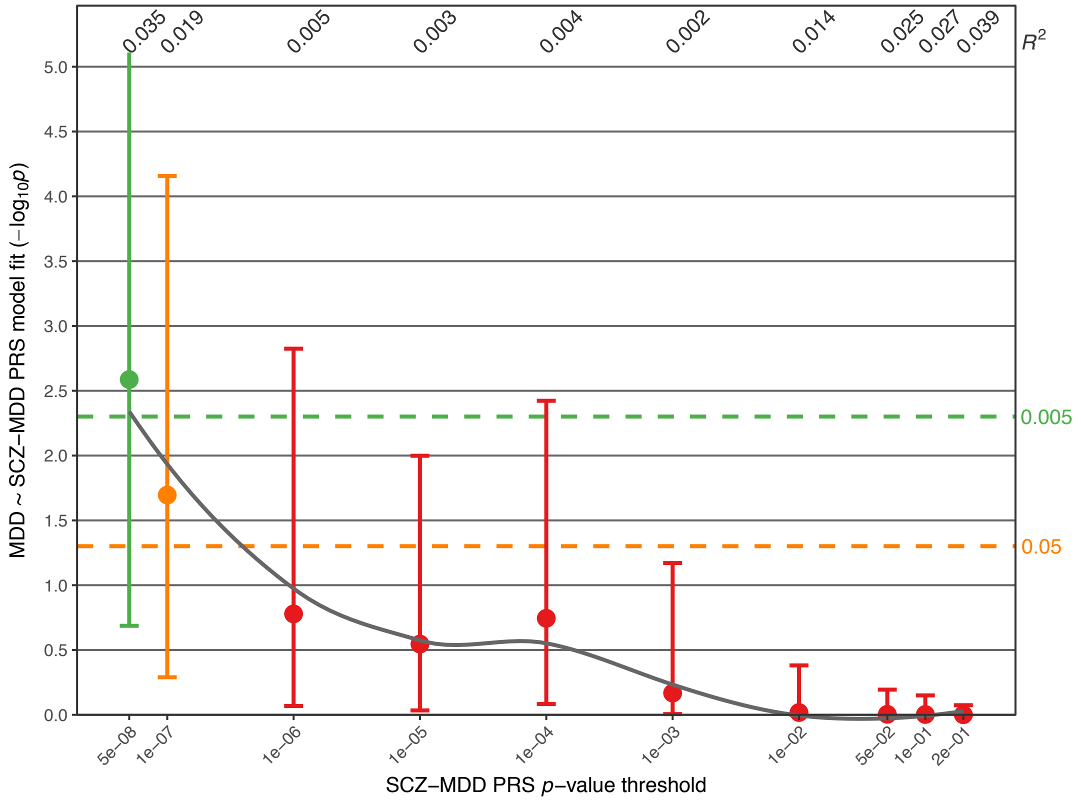

**Supplementary Fig. S10J:** Association of the SCZ-BD GWIS PRS.

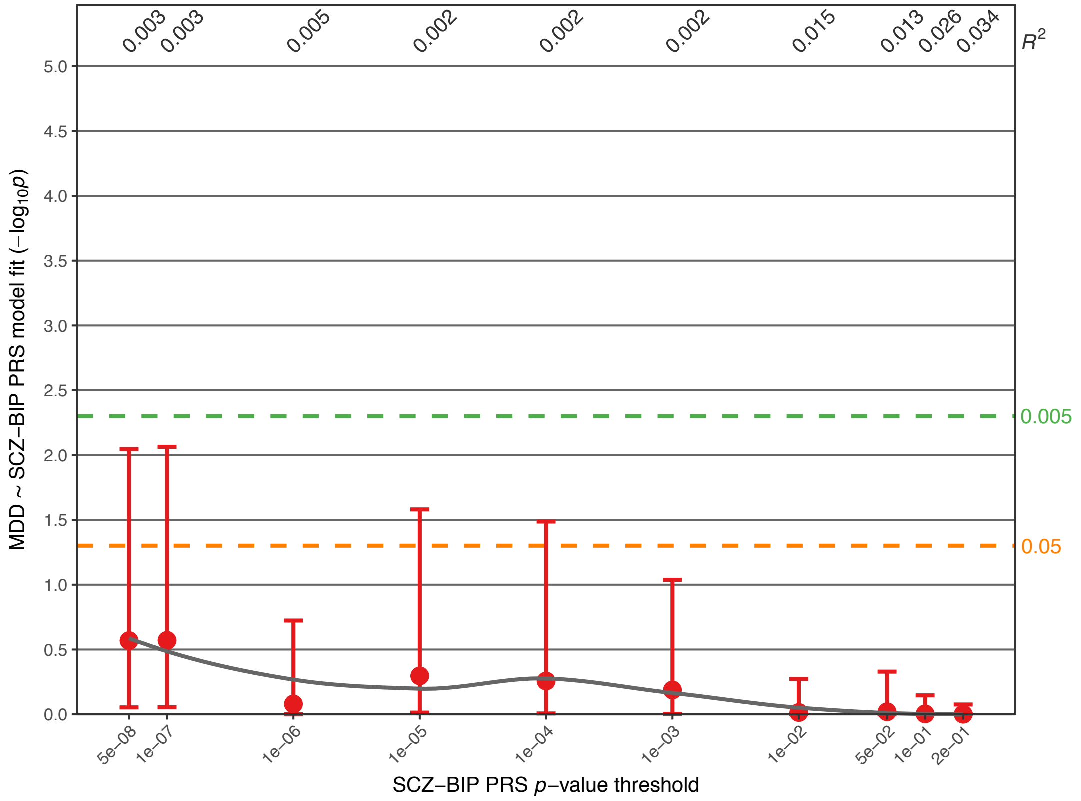


**Supplementary Fig. S10K:** Association of the LOAD PRS.

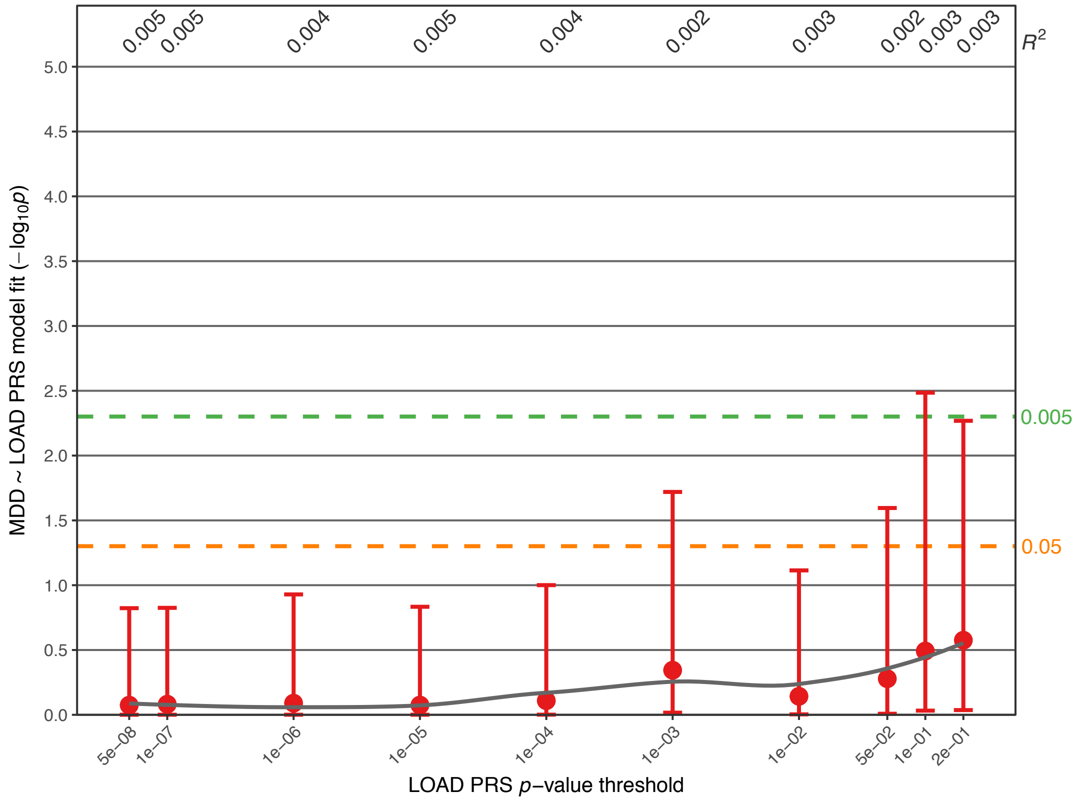

**Supplementary Fig. S11:** Population substructure analysis. Details regarding the generation of MDS components and the population substructure analysis are described above in the Supplementary Methods. The axes have been scaled to show standard deviations.

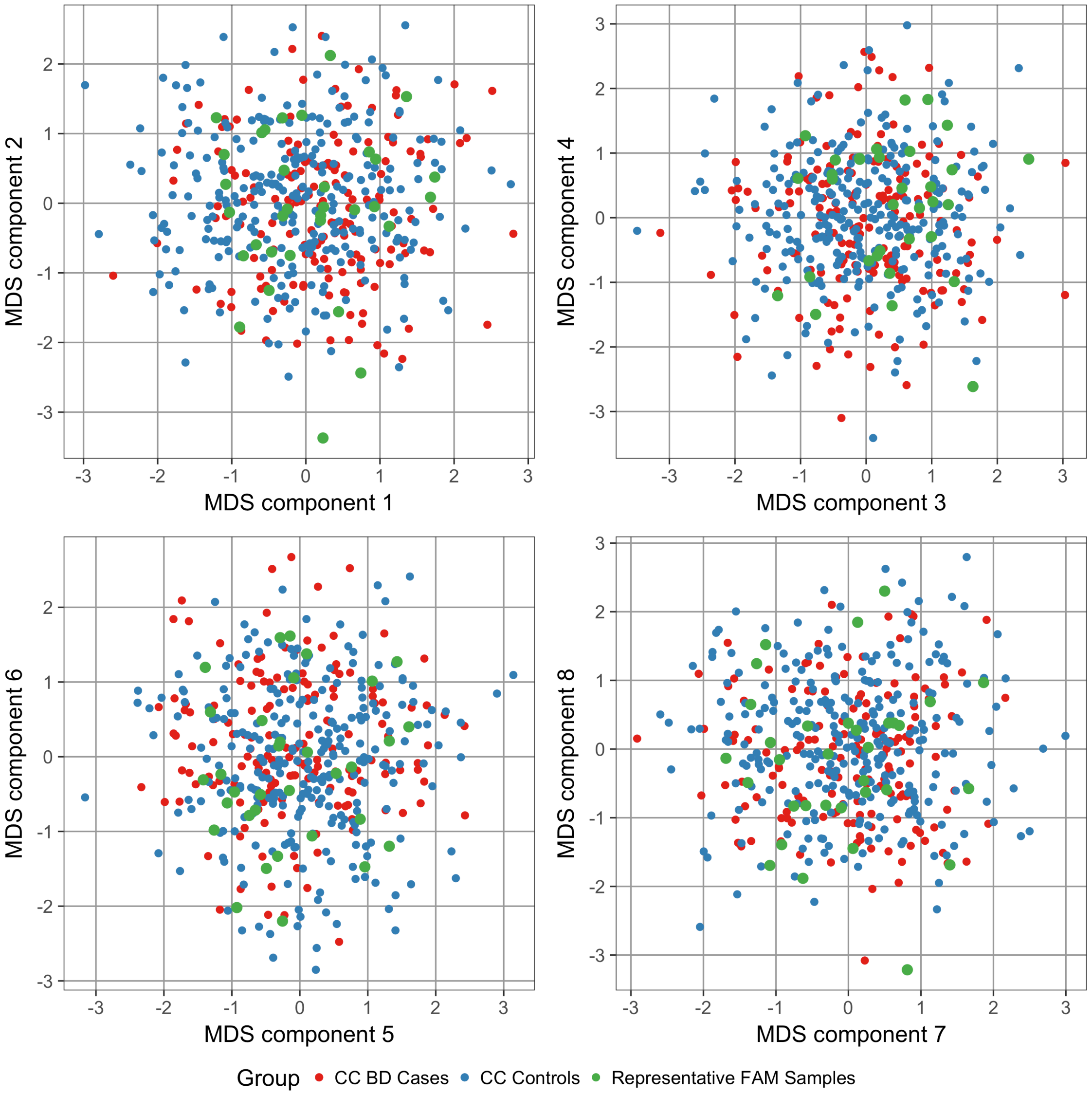

**Supplementary Fig. S12:** Association analysis comparing BD PRS in sporadic $CC_{BD}$ cases and $CC_{controls}$. Details of the plot are described in the legend for Fig. 1. Covariate used: Sex. Full association test statistics including *p*-values are shown in Supplementary Table S10.

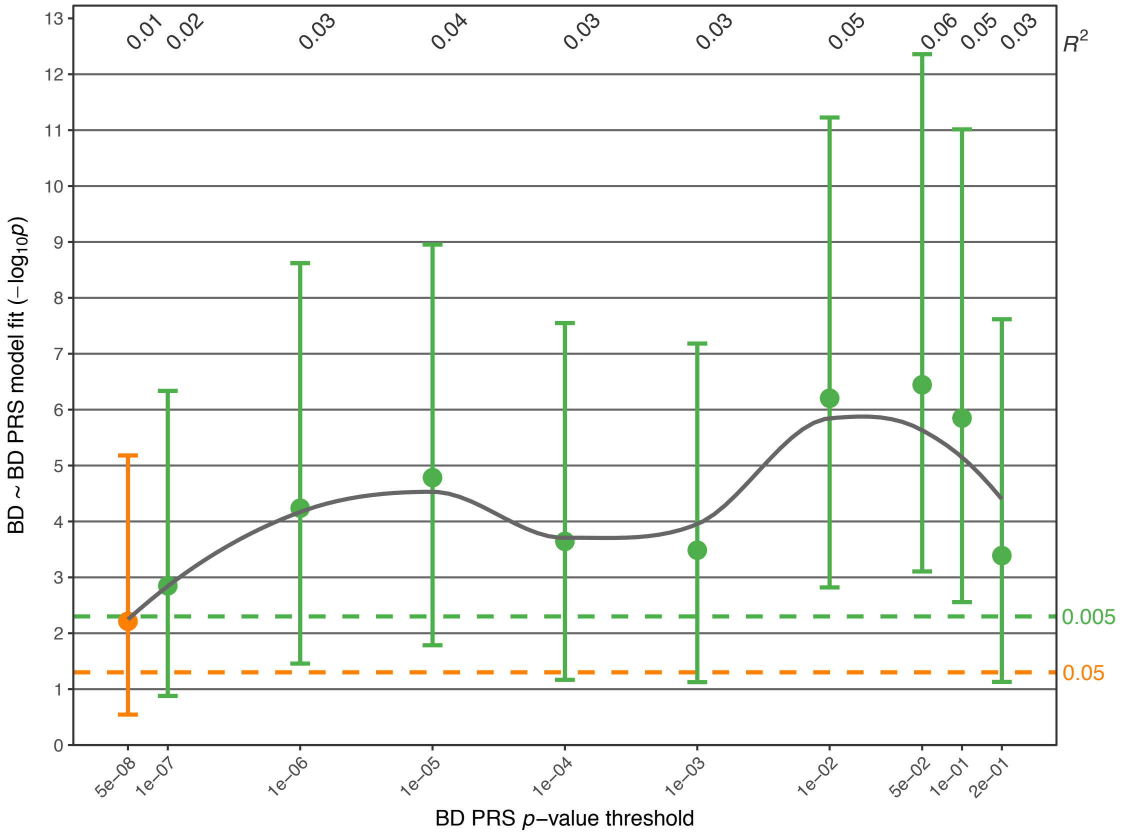

### IGAP Supplementary Methods and Acknowledgments

#### IGAP Methods for the LOAD GWAS

International Genomics of Alzheimer's Project (IGAP) is a large two-stage study based upon genome-wide association studies (GWAS) on individuals of European ancestry. In stage 1, IGAP used genotyped and imputed data on 7,055,881 single nucleotide polymorphisms (SNPs) to meta-analyse four previously-published GWAS datasets consisting of 17,008 Alzheimer's disease cases and 37,154 controls (The European Alzheimer's disease Initiative – EADI the Alzheimer Disease Genetics Consortium – ADGC The Cohorts for Heart and Aging Research in Genomic Epidemiology consortium – CHARGE The Genetic and Environmental Risk in AD consortium – GERAD). In stage 2, 11,632 SNPs were genotyped and tested for association in an independent set of 8,572 Alzheimer's disease cases and 11,312 controls. Finally, a meta-analysis was performed combining results from stages 1 & 2.
The present study used GWAS summary statistics from stage 1 for the calculation of PRS.


#### IGAP Acknowledgments

We thank the International Genomics of Alzheimer's Project (IGAP) for providing summary results data for these analyses. The investigators within IGAP contributed to the design and implementation of IGAP and/or provided data but did not participate in analysis or writing of this report. IGAP was made possible by the generous participation of the control subjects, the patients, and their families. The i–Select chips was funded by the French National Foundation on Alzheimer's disease and related disorders. EADI was supported by the LABEX (laboratory of excellence program investment for the future) DISTALZ grant, Inserm, Institut Pasteur de Lille, Université de Lille 2 and the Lille University Hospital. GERAD was supported by the Medical Research Council (Grant n° 503480), Alzheimer's Research UK (Grant n° 503176), the Wellcome Trust (Grant n° 082604/2/07/Z) and German Federal Ministry of Education and Research (BMBF): Competence Network Dementia (CND) grant n° 01GI0102, 01GI0711, 01GI0420. CHARGE was partly supported by the NIH/NIA grant R01 AG033193 and the NIA AG081220 and AGES contract N01–AG–12100, the NHLBI grant R01 HL105756, the Icelandic Heart Association, and the Erasmus Medical Center and Erasmus University. ADGC was supported by the NIH/NIA grants: U01 AG032984, U24 AG021886, U01 AG016976, and the Alzheimer's Association grant ADGC–10–196728.


#### Authors of the Bipolar Disorder Working Group of the Psychiatric Genomics Consortium


Eli A Stahl 1,2,3† &
Gerome Breen 4,5†
Andreas J Forstner 6,7,8,9,10†
Andrew McQuillin 11†
Stephan Ripke 12,13,14†
Vassily Trubetskoy 13
Manuel Mattheisen 15,16,17,18,19
Yunpeng Wang 20,21
Jonathan R I Coleman 4,5
Héléna A Gaspar 4,5
Christiaan A de Leeuw 22
Stacy Steinberg 23
Jennifer M Whitehead Pavlides 24
Maciej Trzaskowski 25
Tune H Pers 3,26
Peter A Holmans 27
Liam Abbott 12
Esben Agerbo 19,28,29

Huda Akil 30
Diego Albani 31
Ney Alliey-Rodriguez 32
Thomas D Als 15,16,19
Adebayo Anjorin 33
Verneri Antilla 14
Swapnil Awasthi 13
Judith A Badner 34
Marie Bækvad-Hansen 19,35
Jack D Barchas 36
Nicholas Bass 11
Michael Bauer 37
Richard Belliveau 12
Sarah E Bergen 38
Carsten Bøcker Pedersen 19,28,29
Erlend Bøen 39
Marco Boks 40
James Boocock 41
Monika Budde 42
William Bunney 43
Margit Burmeister 44
Jonas Bybjerg-Grauholm 19,35
William Byerley 45
Miquel Casas 46,47,48,49
Felecia Cerrato 12
Pablo Cervantes 50
Kimberly Chambert 12
Alexander W Charney 2
Danfeng Chen 12
Claire Churchhouse 12,14
Toni-Kim Clarke 51
William Coryell 52
David W Craig 53
Cristiana Cruceanu 50,54
Piotr M Czerski 55
Anders M Dale 56,57,58,59
Simone de Jong 4,5
Franziska Degenhardt 8,9
Jurgen Del-Favero 60
J Raymond DePaulo 61
Srdjan Djurovic 62,63
Amanda L Dobbyn 1,2
Ashley Dumont 12
Torbjørn Elvsåshagen 64,65
Valentina Escott-Price 27
Chun Chieh Fan 59
Sascha B Fischer 6,10
Matthew Flickinger 66
Tatiana M Foroud 67
Liz Forty 27
Josef Frank 68
Christine Fraser 27
Nelson B Freimer 69
Louise Frisén 70,71,72
Katrin Gade 42,73
Diane Gage 12
Julie Garnham 74
Claudia Giambartolomei 41
Marianne Giørtz Pedersen 19,28,29
Jaqueline Goldstein 12
Scott D Gordon 75
Katherine Gordon-Smith 76
Elaine K Green 77
Melissa J Green 78
Tiffany A Greenwood 58
Jakob Grove 15,16,19,79
Weihua Guan 80
José Guzman Parra 81
Marian L Hamshere 27
Martin Hautzinger 82
Urs Heilbronner 42
Stefan Herms 6,8,9,10
Maria Hipolito 83
Per Hoffmann 6,8,9,10
Dominic Holland 56,84
Laura Huckins 1,2
Stéphane Jamain 85,86
Jessica S Johnson 1,2
Anders Juréus 38
Radhika Kandaswamy 4
Robert Karlsson 38
James L Kennedy 87,88,89,90
Sarah Kittel-Schneider 91
James A Knowles 92,93
Manolis Kogevinas 94
Anna C Koller 8,9
Ralph Kupka 95,96,97
Catharina Lavebratt 70
Jacob Lawrence 98
William B Lawson 83
Markus Leber 99
Phil H Lee 12,14,100
Shawn E Levy 101
Jun Z Li 102
Chunyu Liu 103
Susanne Lucae 104
Anna Maaser 8,9
Donald J MacIntyre 105,106

Pamela B Mahon 61,107
Wolfgang Maier 108
Lina Martinsson 71
Steve McCarroll 12,109
Peter McGuffin 4
Melvin G McInnis 110
James D McKay 111
Helena Medeiros 93
Sarah E Medland 75
Fan Meng 30,110
Lili Milani 112
Grant W Montgomery 25
Derek W Morris 113,114
Thomas W Mühleisen 6,115
Niamh Mullins 4
Hoang Nguyen 1,2
Caroline M Nievergelt 58,116
Annelie Nordin Adolfsson 117
Evaristus A Nwulia 83
Claire O'Donovan 74
Loes M Olde Loohuis 69
Anil P S Ori 69
Lilijana Oruc 118
Urban Ösby 119
Roy H Perlis 120,121
Amy Perry 76
Andrea Pfennig 37
James B Potash 61
Shaun M Purcell 2,107
Eline J Regeer 122
Andreas Reif 91
Céline S Reinbold 6,10
John P Rice 123
Alexander L Richards 27
Fabio Rivas 81
Margarita Rivera 4,124
Panos Roussos 1,2,125
Douglas M Ruderfer 126
Euijung Ryu 127
Cristina Sánchez-Mora 46,47,49
Alan F Schatzberg 128
William A Scheftner 129
Nicholas J Schork 130
Cynthia Shannon Weickert 78,131
Tatyana Shehktman 58
Paul D Shilling 58
Engilbert Sigurdsson 132
Claire Slaney 74
Olav B Smeland 56,133,134
Janet L Sobell 135
Christine Søholm Hansen 19,35
Anne T Spijker 136
David St Clair 137
Michael Steffens 138
John S Strauss 89,139
Fabian Streit 68
Jana Strohmaier 68
Szabolcs Szelinger 140
Robert C Thompson 110
Thorgeir E Thorgeirsson 23
Jens Treutlein 68
Helmut Vedder 141
Weiqing Wang 1,2
Stanley J Watson 110
Thomas W Weickert 78,131
Stephanie H Witt 68
Simon Xi 142
Wei Xu 143,144
Allan H Young 145
Peter Zandi 146
Peng Zhang 147
Sebastian Zollner 110
Rolf Adolfsson 117
Ingrid Agartz 17,39,148
Martin Alda 74,149
Lena Backlund 71
Bernhard T Baune 150
Frank Bellivier 151,152,153,154
Wade H Berrettini 155
Joanna M Biernacka 127
Douglas H R Blackwood 51
Michael Boehnke 66
Anders D Børglum 15,16,19
Aiden Corvin 114
Nicholas Craddock 27
Mark J Daly 12,14
Udo Dannlowski 156
Tõnu Esko 3,109,112,157
Bruno Etain 151,153,154,158
Mark Frye 159
Janice M Fullerton 131,160
Elliot S Gershon 32,161
Michael Gill 114
Fernando Goes 61
Maria Grigoroiu-Serbanescu 162
Joanna Hauser 55
David M Hougaard 19,35
Christina M Hultman 38

Ian Jones 27
Lisa A Jones 76
René S Kahn 2,40
George Kirov 27
Mikael Landén 38,163
Marion Leboyer 86,151,164
Cathryn M Lewis 4,5,165
Qingqin S Li 166
Jolanta Lissowska 167
Nicholas G Martin 75,168
Fermin Mayoral 81
Susan L McElroy 169
Andrew M McIntosh 51,170
Francis J McMahon 171
Ingrid Melle 172,173
Andres Metspalu 112,174
Philip B Mitchell 78
Gunnar Morken 175,176
Ole Mors 19,177
Preben Bo Mortensen 15,19,28,29
Bertram Müller-Myhsok 54,178,179
Richard M Myers 101
Benjamin M Neale 3,12,14
Vishwajit Nimgaonkar 180
Merete Nordentoft 19,181
Markus M Nöthen 8,9
Michael C O'Donovan 27
Ketil J Oedegaard 182,183
Michael J Owen 27
Sara A Paciga 184
Carlos Pato 93,185
Michele T Pato 93
Danielle Posthuma 22,186
Josep Antoni Ramos-Quiroga 46,47,48,49
Marta Ribasés 46,47,49
Marcella Rietschel 68
Guy A Rouleau 187,188
Martin Schalling 70
Peter R Schofield 131,160
Thomas G Schulze 42,61,68,73,171
Alessandro Serretti 189
Jordan W Smoller 12,190,191
Hreinn Stefansson 23
Kari Stefansson 23,192
Eystein Stordal 193,194
Patrick F Sullivan 38,195,196
Gustavo Turecki 197
Arne E Vaaler 198
Eduard Vieta 199
John B Vincent 139
Thomas Werge 19,200,201
John I Nurnberger 202
Naomi R Wray 24,25
Arianna Di Florio 27,196
Howard J Edenberg 203
Sven Cichon 6,8,10,115
Roel A Ophoff 40,41,69
Laura J Scott 66
Ole A Andreassen 133,134
John Kelsoe 58*&
Pamela Sklar 1,2*^
† Equal contribution, * Co-last authors
^ deceased.

1 Department of Genetics and Genomic Sciences, Icahn School of Medicine at Mount Sinai, New York, NY, US
2 Department of Psychiatry, Icahn School of Medicine at Mount Sinai, New York, NY, US
3 Medical and Population Genetics, Broad Institute, Cambridge, MA, US
4 MRC Social, Genetic and Developmental Psychiatry Centre, King's College London, London, GB
5 NIHR BRC for Mental Health, King's College London, London, GB
6 Human Genomics Research Group, Department of Biomedicine, University of Basel, Basel, CH
7 Department of Psychiatry (UPK), University of Basel, Basel, CH
8 Institute of Human Genetics, University of Bonn, School of Medicine & University Hospital Bonn, Bonn, DE
9 Department of Genomics, Life&Brain Center, University of Bonn, Bonn, DE
10 Institute of Medical Genetics and Pathology, University Hospital Basel, Basel, CH
11 Division of Psychiatry, University College London, London, GB

12 Stanley Center for Psychiatric Research, Broad Institute, Cambridge, MA, US
13 Department of Psychiatry and Psychotherapy, Charité - Universitätsmedizin, Berlin, DE
14 Analytic and Translational Genetics Unit, Massachusetts General Hospital, Boston, MA, US
15 iSEQ, Center for Integrative Sequencing, Aarhus University, Aarhus, DK
16 Department of Biomedicine - Human Genetics, Aarhus University, Aarhus, DK
17 Department of Clinical Neuroscience, Centre for Psychiatry Research, Karolinska Institutet, Stockholm, SE
18 Department of Psychiatry, Psychosomatics and Psychotherapy, Center of Mental Health, University Hospital Würzburg, Würzburg, DE
19 iPSYCH, The Lundbeck Foundation Initiative for Integrative Psychiatric Research, DK
20 Institute of Biological Psychiatry, Mental Health Centre Sct. Hans, Copenhagen, DK
21 Institute of Clinical Medicine, University of Oslo, Oslo, NO
22 Department of Complex Trait Genetics, Center for Neurogenomics and Cognitive Research, Amsterdam Neuroscience, Vrije Universiteit Amsterdam, Amsterdam, NL
23 deCODE Genetics / Amgen, Reykjavik, IS
24 Queensland Brain Institute, The University of Queensland, Brisbane, QLD, AU
25 Institute for Molecular Bioscience, The University of Queensland, Brisbane, QLD, AU
26 Division of Endocrinology and Center for Basic and Translational Obesity Research, Boston Children's Hospital, Boston, MA, US
27 Medical Research Council Centre for Neuropsychiatric Genetics and Genomics, Division of Psychological Medicine and Clinical Neurosciences, Cardiff University, Cardiff, GB
28 National Centre for Register-Based Research, Aarhus University, Aarhus, DK
29 Centre for Integrated Register-based Research, Aarhus University, Aarhus, DK
30 Molecular & Behavioral Neuroscience Institute, University of Michigan, Ann Arbor, MI, US
31 NEUROSCIENCE, Istituto Di Ricerche Farmacologiche Mario Negri, Milano, IT
32 Department of Psychiatry and Behavioral Neuroscience, University of Chicago, Chicago, IL, US
33 Psychiatry, Berkshire Healthcare NHS Foundation Trust, Bracknell, GB
34 Psychiatry, Rush University Medical Center, Chicago, IL, US
35 Center for Neonatal Screening, Department for Congenital Disorders, Statens Serum Institut, Copenhagen, DK
36 Department of Psychiatry, Weill Cornell Medical College, New York, NY, US
37 Department of Psychiatry and Psychotherapy, University Hospital Carl Gustav Carus, Technische Universität Dresden, Dresden, DE
38 Department of Medical Epidemiology and Biostatistics, Karolinska Institutet, Stockholm, SE
39 Department of Psychiatric Research, Diakonhjemmet Hospital, Oslo, NO
40 Psychiatry, UMC Utrecht Hersencentrum Rudolf Magnus, Utrecht, NL
41 Human Genetics, University of California Los Angeles, Los Angeles, CA, US
42 Institute of Psychiatric Phenomics and Genomics (IPPG), University Hospital, LMU Munich, Munich, DE

43 Department of Psychiatry and Human Behavior, University of California, Irvine, Irvine, CA, US
44 Molecular & Behavioral Neuroscience Institute and Department of Computational Medicine & Bioinformatics, University of Michigan, Ann Arbor, MI, US
45 Psychiatry, University of California San Francisco, San Francisco, CA, US
46 Instituto de Salud Carlos III, Biomedical Network Research Centre on Mental Health (CIBERSAM), Madrid, ES
47 Department of Psychiatry, Hospital Universitari Vall d´Hebron, Barcelona, ES
48 Department of Psychiatry and Forensic Medicine, Universitat Autònoma de Barcelona, Barcelona, ES
49 Psychiatric Genetics Unit, Group of Psychiatry Mental Health and Addictions, Vall d´Hebron Research Institut (VHIR), Universitat Autònoma de Barcelona, Barcelona, ES
50 Department of Psychiatry, Mood Disorders Program, McGill University Health Center, Montreal, QC, CA
51 Division of Psychiatry, University of Edinburgh, Edinburgh, GB
52 University of Iowa Hospitals and Clinics, Iowa City, IA, US
53 Translational Genomics, USC, Phoenix, AZ, US
54 Department of Translational Research in Psychiatry, Max Planck Institute of Psychiatry, Munich, DE
55 Department of Psychiatry, Laboratory of Psychiatric Genetics, Poznan University of Medical Sciences, Poznan, PL
56 Department of Neurosciences, University of California San Diego, La Jolla, CA, US
57 Department of Radiology, University of California San Diego, La Jolla, CA, US
58 Department of Psychiatry, University of California San Diego, La Jolla, CA, US
59 Department of Cognitive Science, University of California San Diego, La Jolla, CA, US
60 Applied Molecular Genomics Unit, VIB Department of Molecular Genetics, University of Antwerp, Antwerp, Belgium
61 Department of Psychiatry and Behavioral Sciences, Johns Hopkins University School of Medicine, Baltimore, MD, US
62 Department of Medical Genetics, Oslo University Hospital Ullevål, Oslo, NO
63 NORMENT, KG Jebsen Centre for Psychosis Research, Department of Clinical Science, University of Bergen, Bergen, NO
64 Department of Neurology, Oslo University Hospital, Oslo, NO
65 NORMENT, KG Jebsen Centre for Psychosis Research, Oslo University Hospital, Oslo, NO
66 Center for Statistical Genetics and Department of Biostatistics, University of Michigan, Ann Arbor, MI, US
67 Department of Medical & Molecular Genetics, Indiana University, Indianapolis, IN, US
68 Department of Genetic Epidemiology in Psychiatry, Central Institute of Mental Health, Medical Faculty Mannheim, Heidelberg University, Mannheim, DE
69 Center for Neurobehavioral Genetics, University of California Los Angeles, Los Angeles, CA, US
70 Department of Molecular Medicine and Surgery, Karolinska Institutet and Center for Molecular Medicine, Karolinska University Hospital, Stockholm, SE
71 Department of Clinical Neuroscience, Karolinska Institutet and Center for Molecular Medicine, Karolinska University Hospital, Stockholm, SE

72 Child and Adolescent Psychiatry Research Center, Stockholm, SE
73 Department of Psychiatry and Psychotherapy, University Medical Center Göttingen, Göttingen, DE
74 Department of Psychiatry, Dalhousie University, Halifax, NS, CA
75 Genetics and Computational Biology, QIMR Berghofer Medical Research Institute, Brisbane, QLD, AU
76 Department of Psychological Medicine, University of Worcester, Worcester, GB
77 School of Biomedical and Healthcare Sciences, Plymouth University Peninsula Schools of Medicine and Dentistry, Plymouth, GB
78 School of Psychiatry, University of New South Wales, Sydney, NSW, AU
79 Bioinformatics Research Centre, Aarhus University, Aarhus, DK
80 Biostatistics, University of Minnesota System, Minneapolis, MN, US
81 Mental Health Department, University Regional Hospital, Biomedicine Institute (IBIMA), Málaga, ES
82 Department of Psychology, Eberhard Karls Universität Tübingen, Tubingen, DE
83 Department of Psychiatry and Behavioral Sciences, Howard University Hospital, Washington, DC, US
84 Center for Multimodal Imaging and Genetics, University of California San Diego, La Jolla, CA, US
85 Psychiatrie Translationnelle, Inserm U955, Créteil, FR
86 Faculté de Médecine, Université Paris Est, Créteil, FR
87 Campbell Family Mental Health Research Institute, Centre for Addiction and Mental Health, Toronto, ON, CA
88 Neurogenetics Section, Centre for Addiction and Mental Health, Toronto, ON, CA
89 Department of Psychiatry, University of Toronto, Toronto, ON, CA
90 Institute of Medical Sciences, University of Toronto, Toronto, ON, CA
91 Department of Psychiatry, Psychosomatic Medicine and Psychotherapy, University Hospital Frankfurt, Frankfurt am Main, DE
92 Cell Biology, SUNY Downstate Medical Center College of Medicine, Brooklyn, NY, US
93 Institute for Genomic Health, SUNY Downstate Medical Center College of Medicine, Brooklyn, NY, US
94 ISGlobal, Barcelona, ES
95 Psychiatry, Altrecht, Utrecht, NL
96 Psychiatry, GGZ inGeest, Amsterdam, NL
97 Psychiatry, VU medisch centrum, Amsterdam, NL
98 Psychiatry, North East London NHS Foundation Trust, Ilford, GB
99 Clinic for Psychiatry and Psychotherapy, University Hospital Cologne, Cologne, DE
100 Psychiatric and Neurodevelopmental Genetics Unit, Massachusetts General Hospital, Boston, MA, US
101 HudsonAlpha Institute for Biotechnology, Huntsville, AL, US
102 Department of Human Genetics, University of Michigan, Ann Arbor, MI, US
103 Psychiatry, University of Illinois at Chicago College of Medicine, Chicago, IL, US
104 Max Planck Institute of Psychiatry, Munich, DE
105 Mental Health, NHS 24, Glasgow, GB
106 Division of Psychiatry, Centre for Clinical Brain Sciences, University of Edinburgh, Edinburgh, GB
107 Psychiatry, Brigham and Women's Hospital, Boston, MA, US
108 Department of Psychiatry and Psychotherapy, University of Bonn, Bonn, DE

109 Department of Genetics, Harvard Medical School, Boston, MA, US
110 Department of Psychiatry, University of Michigan, Ann Arbor, MI, US
111 Genetic Cancer Susceptibility Group, International Agency for Research on Cancer, Lyon, FR
112 Estonian Genome Center, University of Tartu, Tartu, EE
113 Discipline of Biochemistry, Neuroimaging and Cognitive Genomics (NICOG) Centre, National University of Ireland, Galway, Galway, IE
114 Neuropsychiatric Genetics Research Group, Dept of Psychiatry and Trinity Translational Medicine Institute, Trinity College Dublin, Dublin, IE
115 Institute of Neuroscience and Medicine (INM-1), Research Centre Jülich, Jülich, DE
116 Research/Psychiatry, Veterans Affairs San Diego Healthcare System, San Diego, CA, US
117 Department of Clinical Sciences, Psychiatry, Umeå University Medical Faculty, Umeå, SE
118 Department of Clinical Psychiatry, Psychiatry Clinic, Clinical Center University of Sarajevo, Sarajevo, BA
119 Department of Neurobiology, Care sciences, and Society, Karolinska Institutet and Center for Molecular Medicine, Karolinska University Hospital, Stockholm, SE
120 Psychiatry, Harvard Medical School, Boston, MA, US
121 Division of Clinical Research, Massachusetts General Hospital, Boston, MA, US
122 Outpatient Clinic for Bipolar Disorder, Altrecht, Utrecht, NL
123 Department of Psychiatry, Washington University in Saint Louis, Saint Louis, MO, US
124 Department of Biochemistry and Molecular Biology II, Institute of Neurosciences, Center for Biomedical Research, University of Granada, Granada, ES
125 Department of Neuroscience, Icahn School of Medicine at Mount Sinai, New York, NY, US
126 Medicine, Psychiatry, Biomedical Informatics, Vanderbilt University Medical Center, Nashville, TN, US
127 Department of Health Sciences Research, Mayo Clinic, Rochester, MN, US
128 Psychiatry and Behavioral Sciences, Stanford University School of Medicine, Stanford, CA, US
129 Rush University Medical Center, Chicago, IL, US
130 Scripps Translational Science Institute, La Jolla, CA, US
131 Neuroscience Research Australia, Sydney, NSW, AU
132 Faculty of Medicine, Department of Psychiatry, School of Health Sciences, University of Iceland, Reykjavik, IS
133 Div Mental Health and Addiction, Oslo University Hospital, Oslo, NO
134 NORMENT, University of Oslo, Oslo, NO
135 Psychiatry and the Behavioral Sciences, University of Southern California, Los Angeles, CA, US
136 Mood Disorders, PsyQ, Rotterdam, NL
137 Institute for Medical Sciences, University of Aberdeen, Aberdeen, UK
138 Research Division, Federal Institute for Drugs and Medical Devices (BfArM), Bonn, DE
139 Centre for Addiction and Mental Health, Toronto, ON, CA
140 Neurogenomics, TGen, Los Angeles, AZ, US
141 Psychiatry, Psychiatrisches Zentrum Nordbaden, Wiesloch, DE

142 Computational Sciences Center of Emphasis, Pfizer Global Research and Development, Cambridge, MA, US
143 Department of Biostatistics, Princess Margaret Cancer Centre, Toronto, ON, CA
144 Dalla Lana School of Public Health, University of Toronto, Toronto, ON, CA
145 Psychological Medicine, Institute of Psychiatry, Psychology & Neuroscience, King's College London, London, GB
146 Department of Mental Health, Johns Hopkins University Bloomberg School of Public Health, Baltimore, MD, US
147 Institute of Genetic Medicine, Johns Hopkins University School of Medicine, Baltimore, MD, US
148 NORMENT, KG Jebsen Centre for Psychosis Research, Division of Mental Health and Addiction, Institute of Clinical Medicine and Diakonhjemmet Hospital, University of Oslo, Oslo, NO
149 National Institute of Mental Health, Klecany, CZ
150 Discipline of Psychiatry, University of Adelaide, Adelaide, SA, AU
151 Department of Psychiatry and Addiction Medicine, Assistance Publique - Hôpitaux de Paris, Paris, FR
152 Paris Bipolar and TRD Expert Centres, FondaMental Foundation, Paris, FR
153 UMR-S1144 Team 1: Biomarkers of relapse and therapeutic response in addiction and mood disorders, INSERM, Paris, FR
154 Psychiatry, Université Paris Diderot, Paris, FR
155 Psychiatry, University of Pennsylvania, Philadelphia, PA, US
156 Department of Psychiatry, University of Münster, Münster, DE
157 Division of Endocrinology, Children's Hospital Boston, Boston, MA, US
158 Centre for Affective Disorders, Institute of Psychiatry, Psychology and Neuroscience, London, GB
159 Department of Psychiatry & Psychology, Mayo Clinic, Rochester, MN, US
160 School of Medical Sciences, University of New South Wales, Sydney, NSW, AU
161 Department of Human Genetics, University of Chicago, Chicago, IL, US
162 Biometric Psychiatric Genetics Research Unit, Alexandru Obregia Clinical Psychiatric Hospital, Bucharest, RO
163 Institute of Neuroscience and Physiology, University of Gothenburg, Gothenburg, SE
164 INSERM, Paris, FR
165 Department of Medical & Molecular Genetics, King's College London, London, GB
166 Neuroscience Therapeutic Area, Janssen Research and Development, LLC, Titusville, NJ, US
167 Cancer Epidemiology and Prevention, M. Sklodowska-Curie Cancer Center and Institute of Oncology, Warsaw, PL
168 School of Psychology, The University of Queensland, Brisbane, QLD, AU
169 Research Institute, Lindner Center of HOPE, Mason, OH, US
170 Centre for Cognitive Ageing and Cognitive Epidemiology, University of Edinburgh, Edinburgh, GB
171 Human Genetics Branch, Intramural Research Program, National Institute of Mental Health, Bethesda, MD, US
172 Division of Mental Health and Addiction, Oslo University Hospital, Oslo, NO
173 Division of Mental Health and Addiction, University of Oslo, Institute of Clinical Medicine, Oslo, NO
174 Institute of Molecular and Cell Biology, University of Tartu, Tartu, EE

175 Mental Health, Faculty of Medicine and Health Sciences, Norwegian University of Science and Technology - NTNU, Trondheim, NO
176 Psychiatry, St Olavs University Hospital, Trondheim, NO
177 Psychosis Research Unit, Aarhus University Hospital, Risskov, DK
178 Munich Cluster for Systems Neurology (SyNergy), Munich, DE
179 University of Liverpool, Liverpool, GB
180 Psychiatry and Human Genetics, University of Pittsburgh, Pittsburgh, PA, US
181 Mental Health Services in the Capital Region of Denmark, Mental Health Center Copenhagen, University of Copenhagen, Copenhagen, DK
182 Division of Psychiatry, Haukeland Universitetssjukehus, Bergen, NO
183 Faculty of Medicine and Dentistry, University of Bergen, Bergen, NO
184 Human Genetics and Computational Biomedicine, Pfizer Global Research and Development, Groton, CT, US
185 College of Medicine Institute for Genomic Health, SUNY Downstate Medical Center College of Medicine, Brooklyn, NY, US
186 Department of Clinical Genetics, Amsterdam Neuroscience, Vrije Universiteit Medical Center, Amsterdam, NL
187 Department of Neurology and Neurosurgery, McGill University, Faculty of Medicine, Montreal, QC, CA
188 Montreal Neurological Institute and Hospital, Montreal, QC, CA
189 Department of Biomedical and NeuroMotor Sciences, University of Bologna, Bologna, IT
190 Department of Psychiatry, Massachusetts General Hospital, Boston, MA, US
191 Psychiatric and Neurodevelopmental Genetics Unit (PNGU), Massachusetts General Hospital, Boston, MA, US
192 Faculty of Medicine, University of Iceland, Reykjavik, IS
193 Department of Psychiatry, Hospital Namsos, Namsos, NO
194 Department of Neuroscience, Norges Teknisk Naturvitenskapelige Universitet Fakultet for naturvitenskap og teknologi, Trondheim, NO
195 Department of Genetics, University of North Carolina at Chapel Hill, Chapel Hill, NC, US
196 Department of Psychiatry, University of North Carolina at Chapel Hill, Chapel Hill, NC, US
197 Department of Psychiatry, McGill University, Montreal, QC, CA
198 Dept of Psychiatry, Sankt Olavs Hospital Universitetssykehuset i Trondheim, Trondheim, NO
199 Clinical Institute of Neuroscience, Hospital Clinic, University of Barcelona, IDIBAPS, CIBERSAM, Barcelona, ES
200 Institute of Biological Psychiatry, MHC Sct. Hans, Mental Health Services Copenhagen, Roskilde, DK
201 Department of Clinical Medicine, University of Copenhagen, Copenhagen, DK
202 Psychiatry, Indiana University School of Medicine, Indianapolis, IN, US
203 Biochemistry and Molecular Biology, Indiana University School of Medicine, Indianapolis, IN, US

**Authors of the Major Depressive Disorder Working Group of the Psychiatric Genomics Consortium**


Naomi R Wray* 1, 2
Stephan Ripke* 3, 4, 5
Manuel Mattheisen* 6, 7, 8, 9
Maciej Trzaskowski* 1
Enda M Byrne 1
Abdel Abdellaoui 10
Mark J Adams 11
Esben Agerbo 9, 12, 13
Tracy M Air 14
Till F M Andlauer 15, 16
Silviu-Alin Bacanu 17
Marie Bækvad-Hansen 9, 18
Aartjan T F Beekman 19
Tim B Bigdeli 17, 20
Elisabeth B Binder 15, 21
Douglas H R Blackwood 11
Julien Bryois 22
Henriette N Buttenschøn 8, 9, 23
Jonas Bybjerg-Grauholm 9, 18
Na Cai 24, 25
Enrique Castelao 26
Jane Hvarregaard Christensen 7, 8, 9
Toni-Kim Clarke 11
Jonathan R I Coleman 27
Lucía Colodro-Conde 28
Baptiste Couvy-Duchesne 2, 29
Nick Craddock 30
Gregory E Crawford 31, 32
Gail Davies 33
Ian J Deary 33
Franziska Degenhardt 34, 35
Eske M Derks 28
Nese Direk 36, 37
Conor V Dolan 10
Erin C Dunn 38, 39, 40
Thalia C Eley 27
Valentina Escott-Price 41
Farnush Farhadi Hassan Kiadeh 42
Hilary K Finucane 43, 44
Jerome C Foo 45
Andreas J Forstner 34, 35, 46, 47
Josef Frank 45
Héléna A Gaspar 27
Michael Gill 48
Fernando S Goes 49
Scott D Gordon 28
Jakob Grove 7, 8, 9, 50
Lynsey S Hall 11, 51
Christine Søholm Hansen 9, 18
Thomas F Hansen 52, 53, 54
Stefan Herms 34, 35, 47
Ian B Hickie 55
Per Hoffmann 34, 35, 47
Georg Homuth 56
Carsten Horn 57
Jouke-Jan Hottenga 10
David M Hougaard 9, 18
Marcus Ising 58
Rick Jansen 19
Ian Jones 59
Lisa A Jones 60
Eric Jorgenson 61
James A Knowles 62
Isaac S Kohane 63, 64, 65
Julia Kraft 4
Warren W. Kretzschmar 66
Jesper Krogh 67
Zoltán Kutalik 68, 69
Yihan Li 66
Penelope A Lind 28
Donald J MacIntyre 70, 71
Dean F MacKinnon 49
Robert M Maier 2
Wolfgang Maier 72
Jonathan Marchini 73
Hamdi Mbarek 10
Patrick McGrath 74
Peter McGuffin 27
Sarah E Medland 28
Divya Mehta 2, 75
Christel M Middeldorp 10, 76, 77
Evelin Mihailov 78
Yuri Milaneschi 19
Lili Milani 78
Francis M Mondimore 49
Grant W Montgomery 1
Sara Mostafavi 79, 80
Niamh Mullins 27
Matthias Nauck 81, 82
Bernard Ng 80
Michel G Nivard 10
Dale R Nyholt 83

Paul F O'Reilly 27
Hogni Oskarsson 84
Michael J Owen 59
Jodie N Painter 28
Carsten Bøcker Pedersen 9, 12, 13
Marianne Giørtz Pedersen 9, 12, 13
Roseann E. Peterson 17, 85
Erik Pettersson 22
Wouter J Peyrot 19
Giorgio Pistis 26
Danielle Posthuma 86, 87
Jorge A Quiroz 88
Per Qvist 7, 8, 9
John P Rice 89
Brien P. Riley 17
Margarita Rivera 27, 90
Saira Saeed Mirza 36
Robert Schoevers 91
Eva C Schulte 92, 93
Ling Shen 61
Jianxin Shi 94
Stanley I Shyn 95
Engilbert Sigurdsson 96
Grant C B Sinnamon 97
Johannes H Smit 19
Daniel J Smith 98
Hreinn Stefansson 99
Stacy Steinberg 99
Fabian Streit 45
Jana Strohmaier 45
Katherine E Tansey 100
Henning Teismann 101
Alexander Teumer 102
Wesley Thompson 9, 53, 103, 104
Pippa A Thomson 105
Thorgeir E Thorgeirsson 99
Matthew Traylor 106
Jens Treutlein 45
Vassily Trubetskoy 4
André G Uitterlinden 107
Daniel Umbricht 108
Sandra Van der Auwera 109
Albert M van Hemert 110
Alexander Viktorin 22
Peter M Visscher 1, 2
Yunpeng Wang 9, 53, 104
Bradley T. Webb 111
Shantel Marie Weinsheimer 9, 53
Jürgen Wellmann 101
Gonneke Willemsen 10
Stephanie H Witt 45
Yang Wu 1
Hualin S Xi 112
Jian Yang 2, 113
Futao Zhang 1
Volker Arolt 114
Bernhard T Baune 115
Klaus Berger 101
Dorret I Boomsma 10
Sven Cichon 34, 47, 116, 117
Udo Dannlowski 114
EJC de Geus 10, 118
J Raymond DePaulo 49
Enrico Domenici 119
Katharina Domschke 120
Tõnu Esko 5, 78
Hans J Grabe 109
Steven P Hamilton 121
Caroline Hayward 122
Andrew C Heath 89
Kenneth S Kendler 17
Stefan Kloiber 58, 123, 124
Glyn Lewis 125
Qingqin S Li 126
Susanne Lucae 58
Pamela AF Madden 89
Patrik K Magnusson 22
Nicholas G Martin 28
Andrew M McIntosh 11, 33
Andres Metspalu 78, 127
Ole Mors 9, 128
Preben Bo Mortensen 8, 9, 12, 13
Bertram Müller-Myhsok 15, 16, 129
Merete Nordentoft 9, 130
Markus M Nöthen 34, 35
Michael C O'Donovan 59
Sara A Paciga 131
Nancy L Pedersen 22
Brenda WJH Penninx 19
Roy H Perlis 38, 132
David J Porteous 105
James B Potash 133
Martin Preisig 26
Marcella Rietschel 45
Catherine Schaefer 61
Thomas G Schulze 45, 93, 134, 135, 136
Jordan W Smoller 38, 39, 40

Kari Stefansson 99, 137
Henning Tiemeier 36, 138, 139
Rudolf Uher 140
Henry Völzke 102
Myrna M Weissman 74, 141
Thomas Werge 9, 53, 142
Cathryn M Lewis 27, 143
Douglas F Levinson 144
Gerome Breen 27, 145
Anders D Børglum 7, 8, 9
Patrick F Sullivan 22, 146, 147

1 Institute for Molecular Bioscience, The University of Queensland, Brisbane, QLD, AU
2 Queensland Brain Institute, The University of Queensland, Brisbane, QLD, AU
3 Analytic and Translational Genetics Unit, Massachusetts General Hospital, Boston, MA, US
4 Department of Psychiatry and Psychotherapy, Universitätsmedizin Berlin Campus Charité Mitte, Berlin, DE
5 Medical and Population Genetics, Broad Institute, Cambridge, MA, US
6 Centre for Psychiatry Research, Department of Clinical Neuroscience, Karolinska Institutet, Stockholm, SE
7 Department of Biomedicine, Aarhus University, Aarhus, DK
8 iSEQ, Centre for Integrative Sequencing, Aarhus University, Aarhus, DK
9 iPSYCH, The Lundbeck Foundation Initiative for Integrative Psychiatric Research,, DK
10 Dept of Biological Psychology & EMGO+ Institute for Health and Care Research, Vrije Universiteit Amsterdam, Amsterdam, NL
11 Division of Psychiatry, University of Edinburgh, Edinburgh, GB
12 Centre for Integrated Register-based Research, Aarhus University, Aarhus, DK
13 National Centre for Register-Based Research, Aarhus University, Aarhus, DK
14 Discipline of Psychiatry, University of Adelaide, Adelaide, SA, AU
15 Department of Translational Research in Psychiatry, Max Planck Institute of Psychiatry, Munich, DE
16 Munich Cluster for Systems Neurology (SyNergy), Munich, DE
17 Department of Psychiatry, Virginia Commonwealth University, Richmond, VA, US
18 Center for Neonatal Screening, Department for Congenital Disorders, Statens Serum Institut, Copenhagen, DK
19 Department of Psychiatry, Vrije Universiteit Medical Center and GGZ inGeest, Amsterdam, NL
20 Virginia Institute for Psychiatric and Behavior Genetics, Richmond, VA, US
21 Department of Psychiatry and Behavioral Sciences, Emory University School of Medicine, Atlanta, GA, US
22 Department of Medical Epidemiology and Biostatistics, Karolinska Institutet, Stockholm, SE
23 Department of Clinical Medicine, Translational Neuropsychiatry Unit, Aarhus University, Aarhus, DK
24 Human Genetics, Wellcome Trust Sanger Institute, Cambridge, GB
25 Statistical genomics and systems genetics, European Bioinformatics Institute (EMBL-EBI), Cambridge, GB
26 Department of Psychiatry, University Hospital of Lausanne, Prilly, Vaud, CH
27 Social Genetic and Developmental Psychiatry Centre, King's College London, London, GB
28 Genetics and Computational Biology, QIMR Berghofer Medical Research Institute, Brisbane, QLD, AU
29 Centre for Advanced Imaging, The University of Queensland, Brisbane, QLD, AU
30 Psychological Medicine, Cardiff University, Cardiff, GB

31 Center for Genomic and Computational Biology, Duke University, Durham, NC, US
32 Department of Pediatrics, Division of Medical Genetics, Duke University, Durham, NC, US
33 Centre for Cognitive Ageing and Cognitive Epidemiology, University of Edinburgh, Edinburgh, GB
34 Institute of Human Genetics, University of Bonn, Bonn, DE
35 Life&Brain Center, Department of Genomics, University of Bonn, Bonn, DE
36 Epidemiology, Erasmus MC, Rotterdam, Zuid-Holland, NL
37 Psychiatry, Dokuz Eylul University School Of Medicine, Izmir, TR
38 Department of Psychiatry, Massachusetts General Hospital, Boston, MA, US
39 Psychiatric and Neurodevelopmental Genetics Unit (PNGU), Massachusetts General Hospital, Boston, MA, US
40 Stanley Center for Psychiatric Research, Broad Institute, Cambridge, MA, US
41 Neuroscience and Mental Health, Cardiff University, Cardiff, GB
42 Bioinformatics, University of British Columbia, Vancouver, BC, CA
43 Department of Epidemiology, Harvard T.H. Chan School of Public Health, Boston, MA, US
44 Department of Mathematics, Massachusetts Institute of Technology, Cambridge, MA, US
45 Department of Genetic Epidemiology in Psychiatry, Central Institute of Mental Health, Medical Faculty Mannheim, Heidelberg University, Mannheim, Baden-Württemberg, DE
46 Department of Psychiatry (UPK), University of Basel, Basel, CH
47 Human Genomics Research Group, Department of Biomedicine, University of Basel, Basel, CH
48 Department of Psychiatry, Trinity College Dublin, Dublin, IE
49 Psychiatry & Behavioral Sciences, Johns Hopkins University, Baltimore, MD, US
50 Bioinformatics Research Centre, Aarhus University, Aarhus, DK
51 Institute of Genetic Medicine, Newcastle University, Newcastle upon Tyne, GB
52 Danish Headache Centre, Department of Neurology, Rigshospitalet, Glostrup, DK
53 Institute of Biological Psychiatry, Mental Health Center Sct. Hans, Mental Health Services Capital Region of Denmark, Copenhagen, DK
54 iPSYCH, The Lundbeck Foundation Initiative for Psychiatric Research, Copenhagen, DK
55 Brain and Mind Centre, University of Sydney, Sydney, NSW, AU
56 Interfaculty Institute for Genetics and Functional Genomics, Department of Functional Genomics, University Medicine and Ernst Moritz Arndt University Greifswald, Greifswald, Mecklenburg-Vorpommern, DE
57 Roche Pharmaceutical Research and Early Development, Pharmaceutical Sciences, Roche Innovation Center Basel, F. Hoffmann-La Roche Ltd, Basel, CH
58 Max Planck Institute of Psychiatry, Munich, DE
59 MRC Centre for Neuropsychiatric Genetics and Genomics, Cardiff University, Cardiff, GB
60 Department of Psychological Medicine, University of Worcester, Worcester, GB
61 Division of Research, Kaiser Permanente Northern California, Oakland, CA, US
62 Psychiatry & The Behavioral Sciences, University of Southern California, Los Angeles, CA, US
63 Department of Biomedical Informatics, Harvard Medical School, Boston, MA, US
64 Department of Medicine, Brigham and Women's Hospital, Boston, MA, US
65 Informatics Program, Boston Children's Hospital, Boston, MA, US
66 Wellcome Trust Centre for Human Genetics, University of Oxford, Oxford, GB
67 Department of Endocrinology at Herlev University Hospital, University of Copenhagen, Copenhagen, DK

68 Institute of Social and Preventive Medicine (IUMSP), University Hospital of Lausanne, Lausanne, VD, CH
69 Swiss Institute of Bioinformatics, Lausanne, VD, CH
70 Division of Psychiatry, Centre for Clinical Brain Sciences, University of Edinburgh, Edinburgh, GB
71 Mental Health, NHS 24 Glasgow, GB
72 Department of Psychiatry and Psychotherapy, University of Bonn, Bonn, DE
73 Statistics, University of Oxford, Oxford, GB
74 Psychiatry, Columbia University College of Physicians and Surgeons, New York, NY, US
75 School of Psychology and Counseling, Queensland University of Technology, Brisbane, QLD, AU
76 Child and Youth Mental Health Service, Children's Health Queensland Hospital and Health Service, South Brisbane, QLD, AU
77 Child Health Research Centre, University of Queensland, Brisbane, QLD, AU
78 Estonian Genome Center, University of Tartu, Tartu, EE
79 Medical Genetics, University of British Columbia, Vancouver, BC, CA
80 Statistics, University of British Columbia, Vancouver, BC, CA
81 DZHK (German Centre for Cardiovascular Research), Partner Site Greifswald, University Medicine, University Medicine Greifswald, Greifswald, Mecklenburg-Vorpommern, DE
82 Institute of Clinical Chemistry and Laboratory Medicine, University Medicine Greifswald, Greifswald, Mecklenburg-Vorpommern, DE
83 Institute of Health and Biomedical Innovation, Queensland University of Technology, Brisbane, QLD, AU
84 Humus, Reykjavik, IS
85 Virginia Institute for Psychiatric & Behavioral Genetics, Virginia Commonwealth University, Richmond, VA, US
86 Clinical Genetics, Vrije Universiteit Medical Center, Amsterdam, NL
87 Complex Trait Genetics, Vrije Universiteit Amsterdam, Amsterdam, NL
88 Solid Biosciences, Boston, MA, US
89 Department of Psychiatry, Washington University in Saint Louis School of Medicine, Saint Louis, MO, US
90 Department of Biochemistry and Molecular Biology II, Institute of Neurosciences, Center for Biomedical Research, University of Granada, Granada, ES
91 Department of Psychiatry, University of Groningen, University Medical Center Groningen, Groningen, NL
92 Department of Psychiatry and Psychotherapy, Medical Center of the University of Munich, Campus Innenstadt, Munich, DE
93 Institute of Psychiatric Phenomics and Genomics (IPPG), Medical Center of the University of Munich, Campus Innenstadt, Munich, DE
94 Division of Cancer Epidemiology and Genetics, National Cancer Institute, Bethesda, MD, US
95 Behavioral Health Services, Kaiser Permanente Washington, Seattle, WA, US
96 Faculty of Medicine, Department of Psychiatry, University of Iceland, Reykjavik, IS
97 School of Medicine and Dentistry, James Cook University, Townsville, QLD, AU
98 Institute of Health and Wellbeing, University of Glasgow, Glasgow, GB
99 deCODE Genetics / Amgen, Reykjavik, IS
100 College of Biomedical and Life Sciences, Cardiff University, Cardiff, GB
101 Institute of Epidemiology and Social Medicine, University of Münster, Münster, Nordrhein-Westfalen, DE
102 Institute for Community Medicine, University Medicine Greifswald, Greifswald, Mecklenburg-Vorpommern, DE

103 Department of Psychiatry, University of California, San Diego, San Diego, CA, US
104 KG Jebsen Centre for Psychosis Research, Norway Division of Mental Health and Addiction, Oslo University Hospital, Oslo, NO
105 Medical Genetics Section, CGEM, IGMM, University of Edinburgh, Edinburgh, GB
106 Clinical Neurosciences, University of Cambridge, Cambridge, GB
107 Internal Medicine, Erasmus MC, Rotterdam, Zuid-Holland, NL
108 Roche Pharmaceutical Research and Early Development, Neuroscience, Ophthalmology and Rare Diseases Discovery & Translational Medicine Area, Roche Innovation Center Basel, F. Hoffmann-La Roche Ltd, Basel, CH
109 Department of Psychiatry and Psychotherapy, University Medicine Greifswald, Greifswald, Mecklenburg-Vorpommern, DE
110 Department of Psychiatry, Leiden University Medical Center, Leiden, NL
111 Virginia Institute for Psychiatric & Behavioral Genetics, Virginia Commonwealth University, Richmond, VA, US
112 Computational Sciences Center of Emphasis, Pfizer Global Research and Development, Cambridge, MA, US
113 Institute for Molecular Bioscience; Queensland Brain Institute, The University of Queensland, Brisbane, QLD, AU
114 Department of Psychiatry, University of Münster, Münster, Nordrhein-Westfalen, DE
115 Department of Psychiatry, Melbourne Medical School, University of Melbourne, Melbourne, AU
116 Institute of Medical Genetics and Pathology, University Hospital Basel, University of Basel, Basel, CH
117 Institute of Neuroscience and Medicine (INM-1), Research Center Juelich, Juelich, DE
118 Amsterdam Public Health Institute, Vrije Universiteit Medical Center, Amsterdam, NL
119 Centre for Integrative Biology, Università degli Studi di Trento, Trento, Trentino-Alto Adige, IT
120 Department of Psychiatry and Psychotherapy, Medical Center, University of Freiburg, Faculty of Medicine, University of Freiburg, Freiburg, DE
121 Psychiatry, Kaiser Permanente Northern California, San Francisco, CA, US
122 Medical Research Council Human Genetics Unit, Institute of Genetics and Molecular Medicine, University of Edinburgh, Edinburgh, GB
123 Department of Psychiatry, University of Toronto, Toronto, ON, CA
124 Centre for Addiction and Mental Health, Toronto, ON, CA
125 Division of Psychiatry, University College London, London, GB
126 Neuroscience Therapeutic Area, Janssen Research and Development, LLC, Titusville, NJ, US
127 Institute of Molecular and Cell Biology, University of Tartu, Tartu, EE
128 Psychosis Research Unit, Aarhus University Hospital, Risskov, Aarhus, DK
129 University of Liverpool, Liverpool, GB
130 Mental Health Center Copenhagen, Copenhagen Universtity Hospital, Copenhagen, DK
131 Human Genetics and Computational Biomedicine, Pfizer Global Research and Development, Groton, CT, US
132 Psychiatry, Harvard Medical School, Boston, MA, US
133 Psychiatry, University of Iowa, Iowa City, IA, US
134 Department of Psychiatry and Behavioral Sciences, Johns Hopkins University, Baltimore, MD, US
135 Department of Psychiatry and Psychotherapy, University Medical Center Göttingen, Goettingen, Niedersachsen, DE

136 Human Genetics Branch, NIMH Division of Intramural Research Programs, Bethesda, MD, US
137 Faculty of Medicine, University of Iceland, Reykjavik, IS
138 Child and Adolescent Psychiatry, Erasmus MC, Rotterdam, Zuid-Holland, NL
139 Psychiatry, Erasmus MC, Rotterdam, Zuid-Holland, NL
140 Psychiatry, Dalhousie University, Halifax, NS, CA
141 Division of Epidemiology, New York State Psychiatric Institute, New York, NY, US
142 Department of Clinical Medicine, University of Copenhagen, Copenhagen, DK
143 Department of Medical & Molecular Genetics, King's College London, London, GB
144 Psychiatry & Behavioral Sciences, Stanford University, Stanford, CA, US
145 NIHR Maudsley Biomedical Research Centre, King's College London, London, GB
146 Genetics, University of North Carolina at Chapel Hill, Chapel Hill, NC, US
147 Psychiatry, University of North Carolina at Chapel Hill, Chapel Hill, NC, US
